# *In vitro*-generated murine monocyte-derived dendritic cells habor a unique identity compared to other mononuclear phagocytes, largely conserved with their human and putative in vivo counterparts

**DOI:** 10.64898/2026.10.09.758081

**Authors:** Abel Lopez-Hernandez, Laurent Hannouche, Laurine Gil, Sylvain Bigot, Laurence Borge, Marc Barad, Cyrille Mionnet, Camille Pierini-Malosse, Romain Fenouil, Bertrand Escalière, Jean Descamps, Sergio Villazala Merino, Mauro Gaya, Pierre Milpied, Marc Dalod

## Abstract

Whether the cells differentiating *in vitro* from total murine bone marrow (BM) cultures with GM-CSF ± IL-4 are homologous to the human cells derived in vitro from purified circulating monocytes (Mo) cultured with GM-CSF and IL-4, mostly correspond to monocyte-derived dendritic cells (moDCs) or rather to macrophages (MFs), and have in vivo counterparts, is debated. It has been proposed to define cell types based on their ontogeny, leading to the proposal of excluding from the dendritic cell (DC) family the cells that do not arise from the common DC progenitors (CDP). However, recent discoveries support defining cell types rather by functions, or, as a surrogate, their instructive molecular networks. Indeed, Langerhans cells, Rorgt+ DCs, some cDC2As and human in vitro-derived moDCs do not arise from the CDP but exert hallmark DC functions instructed by specific transcriptomic modules shared with cDCs. Actually, the study of both LCs and moDCs has been key for dissecting DC biology. Yet, the existence of moDCs is put into questions. The functions once attributed to moDCs are now proposed to be exerted by type 2B conventional DCs (cDC2Bs) or type 3 DCs (DC3s) that have been characterized recently, harbor strongly overlapping phenotypes with moDCs and a similar mixed expression of genes from both the cDC and Mo/MF lineages, but a different ontogeny. To clarify these issues, we generated a transcriptomic atlas of *in vitro*-derived mononuclear phagocytes (MNPs). We show that moDCs are the only DC type arising in BM cultures in the presence of GM-CSF and IL-4. They express a unique transcriptomic and phenotypic signature, different from those of cDC2Bs and DC3s that are produced in FLT3-L cultures and appear to correspond to the same DC type differentiating though two different but converging ontogenetic paths. By harnessing the phenotypic and transcriptomic signatures identified from the analysis of our in vitro MNP atlas, we are able to identify in vivo a population of putative DCs differentiated from monocytes derived from the common monocyte-granulocyte progenitor (GMP), in a respiratory allergy model, specifically in the lungs of mice sensitized to house dust mite (HDM). Using the model of thioglycolate-induced peritonitis, we also characterized the CD226^+^ Mo-derived cells that have been reported to be moDCs or small cavity MFs (SCM) depending on studies. We showed that these cells are mainly differentiated from monocytes derived from the MDP, and split into two populations, one resembling MFs (SCMs) and the other one closer to lung moDCs but expressing distinct genes suggesting a different functional potential. Collectively, we show that the generation and characterization of specific MNP types in vitro can be harnessed to rigorously define their *in vivo* counterparts based on phenotypic analysis and gene module assessment, as we demonstrate for moDCs.

## Introduction

Dendritic cells (DCs) are a population of professional antigen-presenting cells (APCs) with a unique efficacy to activate naïve T cells upon encounter with their cognate antigen. Hence, DCs are key for orchestrating adaptive immunity by inducing the initial activation of T cells and instructing their functional polarization. The ability of DCs to generate a variety of immune responses, each adapted to a specific type of threat, stems from both their heterogeneity, with distinct DC types excelling in different functions, and their plasticity, with each DC type adapting its response to the pathophysiological context^1^. Certain types of DCs have been well characterized, which has led to a consensus on how to define them based on their ontogeny, molecular make up and functions, as it is the case for type 1 conventional DCs (cDC1s)^2^. In contrast, rigorous characterization of the MHC-II⁺SIRPα⁺CD11b⁺ APC compartment (CD11b^+^ APCs) has proven extremely challenging, due to its high heterogeneity since it encompasses cell types of distinct ontogeny but that are difficult to discriminate based on their phenotype both in mice and humans. Indeed, CD11b^+^ APCs may encompass at least 6 different lineages, which precise ontogenetic and functional relationships are not entirely clear: type 2A cDCs (cDC2As), as well as type 2B cDCs (cDC2Bs), type 3 DCs (DC3s), inflammatory DC2s (inf-DC2s), monocyte(Mo)-derived DCs (moDCs) and macrophages (MFs)^3,4^. This complexity of the CD11b^+^ APC population may contribute to its functional plasticity, with contradictory reports on its role in mediating immune activation versus immune suppression and in polarizing T cell responses towards different functions^4–6^. Existing fate mapping models fail rigorously resolving the ontogenic relationships between the subsets of CD11b^+^ APCs. A universal method for distinguishing them robustly *in vivo* is lacking. Hence, to better understand how CD11b^+^ APCs shape immune responses, it is key to be able to rigorously discriminate from one another the distinct cell types constituting this population and to determine whether a division of labor exists between them depending on the pathophysiological context.

Classical Mos comprise 2 main populations, the main one derives from *Ms4a3*^Cre^-labeled granulocyte-Mo progenitors (GMPs) and the other from the small fraction of Mo-DC progenitors (MDPs) labeled by *Clec9a*^iCre7–10^. It is not clear whether moDCs can differentiate both from MDP- and GMP-derived Mos^7^. cDC2Bs are proposed to derive from common DC progenitors (CDPs), labelled by *Clec9a*^iCre^, akin to cDC1s, and DC3s from a Ly6C^+^ MDP fraction labeled by *Ms4a3*^Cre^ akin to what is expected for the moDCs differentiated from GMP-derived Mos. Hence, existing fate mapping models do not resolve the ontogenic relationships between CD11b^+^ APC subsets^5,8,11^ and a universal and robust method for distinguishing them in vivo is lacking, which has hindered deconvoluting CD11b^+^ APC subset functions^11^.

Human moDCs were identified 30 years ago, through *in vitro* differentiation of human circulating Mos with the cytokines GM-CSF and IL-4^12^, the later promoting DC feature acquisition^13^. The culture of total murine bone marrow (BM) with GM-CSF alone or in combination with IL-4 was developed in parallel^14^ and has been traditionally considered as also giving rise to moDCs^14–17^. However, this interpretation has been disputed with a report that the DCs developing in murine GM-CSF BM cultures correspond to cDC2s derived from the CDP, whereas Mos would only generate MFs in these cultures^18^. Yet, cDCs rely on Flt3L for their generation/survival. These issues have initiated a controversy about the very existence of moDCs^15,17,19^, which has been further fueled by the characterization of cDC2B and the discovery of the DC3s and of inf-cDC2s. The DC3s have been described as arising directly from the Ly6C^+^ MDP fraction, without transitioning through a CDP stage, already at steady state^20^. Inf-cDC2s arise from classical cDC2s in inflamed tissues, in particular under conditions of high type I interferon responses such as in the lung during Influenza infection^21^. Similarly to moDCs, cDC2Bs, DC3s and inf-cDC2s express mixed markers of cDCs and Mos/MFs, in both mice and humans^20,22–25^. Therefore, it has been questioned whether moDCs develop at all *in vitro* and *in vivo* ^18,19,26^. The functions that used to be thought to be exerted by moDCs have now been proposed to be reassigned to other cell types, in particular based on the assumption that all previous studies that claimed to have identified and studied moDCs *in vivo* were actually studying misclassified DC3s or inf-DC2s ^20,21,23^. Rigorously demonstrating monocytic origin of DCs *in vivo* requires adoptive transfer^7,11^, which is often infeasible or altering the processes studied. Thus, whether *in vitro*-generated moDCs are homologous between humans and mice, are MFs rather than DCs, and have in vivo counterparts, is debated^15,18,19^.

Three key facts must be acknowledged that support moDC existence and their belonging to the DC family. First, human *in vitro*-derived moDCs exhibit most of the hallmark DC functions; actually, they have been have been used for decades as the archetypical model to dissect the molecular mechanisms controlling antigen processing and presentation, T cell priming and their functional polarization^12,13,15,27–31^.

Second, at least two studies have demonstrated that purified murine Mos can give rise to moDCs *in vitro* in the presence of GM-CSF^32,33^, and one of these studies demonstrated that, upon IL-4 instruction, these murine moDCs excel at antigen cross-presentation for the activation of naïve CD8^+^ T cells, by using molecular mechanisms different from those acting in cDC1s^33^. Hence, similarly to what has been shown for humans, purified mouse Mos can give rise *in vitro* to moDCs endowed with hallmark phenotypic and functional characteristics of *bona fide* DCs.

Third, accumulating evidences show that *bona fide* DCs can develop from different hematopoietic lineages. The vast majority of cDC1s and cDC2Bs derive from the CDP in most pathophysiological conditions^8,34^. However, cDC1s can also develop from lymphoid progenitors in mice where the CDP is genetically depleted^35^, and a significant fraction of cDC2As develop from a dedicated subset of common lymphoid progenitors (CLP) in a *Notch2*-dependent fashion^8,36–39^. Moreover, the newly discovered Rorγt^+^ tolerogenic DCs also appear to develop from a dedicated CLP subset that appears to be shared with the lineages of pDCs and of lymphoid tissue inducer cells^40^. Thus, Langerhans cells, some cDC2As and Rorgt^+^ DCs exert hallmark DC functions despite not arising from the CDP^40–42^. Thus, functions should be used as the main criteria for defining cell types, including DCs, or, as a proxy, the molecular networks instructing them, as can be assessed by gene and protein expression profiling^1,29,43–45^, rather than only ontogeny as previously proposed^26^.

Human *in vitro*-derived moDCs have been used for decades, and are still being used, in clinical trials, for adoptive cellular immunotherapies against not only cancer^15,30^ but also difficult-to-treat viral infections^46–52^, and to promote immune tolerance in solid organ grafts^53–57^. Yet, the clinical benefits are not as high as hoped^30^. Several issues have been proposed to limit treatment efficacy, including the susceptibility of moDCs to functional reprogramming in the tissue microenvironment^58–62^ and their limited ability for migration into the lymph nodes^30,63^. Indeed, *in vitro*-generated moDCs differ substantially from blood or lymphoid organ DCs, based both on their whole gene expression profiling and on their responses to adjuvants^43,64,65^. In cancer, moDC infiltration, as assessed using transcriptomic signatures to deconvolute tissue composition, linked with poor or no prognosis, unlike the favorable association for cDC^66–73^. These observations support the hypothesis that, in tumors, moDCs may exert immunosuppressive functions, possibly overlapping with the monocyte-derived fraction of myeloid-derived suppressor cells (MDSCs), whereas cDC1s and eventually cDC2s are generally protective. However, historical moDC signatures overlap with those of MFs. Hence, a rigorous dissection of the respective identity and functions of moDCs, MDSCs, MFs and cDC2s *in vivo* is warranted. This is key to harness preclinical models and refine the analysis of patient data to better understand the mechanisms limiting the efficacy of moDCs in adoptive cell transfer protocols and how to overcome them.

To resolve these issues, we established a molecular atlas of *in vitro*-derived mononuclear phagocytes (MNPs), through gene expression and phenotypic profiling, by combining fate-mapping and reporter mice with *in vitro* differentiation systems and transcriptomic profiling. We have compared cells generated in total BM cultures versus those arising from purified Mos, under differentiation conditions reported to yield moDCs^32,33^ (GM-CSF and IL-4) versus cDCs (FLT3-L)^74^. Since Zbtb46 has been reported as a specific marker of the DC lineage within hematopoietic cells^75,76^ and fate mapping for Ms4a3 expression used to track monocyte progeny^10^, we have harnessed the *Ms4a3^Cre^*;Ai14;*Zbtb46*^GFP^ (MAZ) mouse strain to unequivocally identify DCs as CD45^+^ *Zbtb46-*GFP^+^ cells and separate as Ai14^+^ the cells derived either from the Ly6C^+^ MDPs (DC3s) or the Mos (moDCs). We showed that moDCs, defined as Zbtb46^+^ Mo-derived cells, are the only DC type present in GM-CSF and IL-4 cultures, whereas DC3s develop in FLT3-L BM cultures. We identified molecular fingerprints discriminating unambiguously moDCs from all other MNPs including DC3s and cDC2Bs, across all of the stimulation conditions used. We observed that the gene expression profiles of DC3s and cDC2Bs strongly resemble each other irrespective of stimulations, supporting the hypothesis that they correspond to the same DC type differentiating though two different but converging ontogenic paths. We identified different flavors of moDCs *in vivo* in a mouse model of respiratory allergy, upon scRNA-seq profiling the CD11b^+^ CD11c^+^ APCs isolated from the lung parenchyma and mediastinal lymph nodes of MAZ mice sensitized to house dust mite (HDM)32, and harnessing the gene modules derived from our atlas of *in vitro*-derived murine MNPs for cell cluster annotations. We showed that GMP-versus MDP-derived Mos can give rise to both distinct and convergent populations of mononuclear phagocytes in the peritoneum upon local induction of a sterile inflammation in response to thioglycolate administration, as assessed upon adoptive transfer experiments in DC45.1 congenic recipient mice of Mo subsets purified from MAZ mice. These findings will enable identifying moDCs across diseases and species, and engineering mutant mice to probe their physiological role and therapeutic potential.

## Results

### Total murine BM cultures in GM-CSF and IL-4 generate primarily moDC, and some macrophages, whereas total BM cultures in FLT3-L cultures generate cDC1s, cDC2s and DC3s

GM-CSF murine BM cultures have been reported to be heterogeneous, encompassing two main populations differing by their level of MHC-II expression. The MHC-II^low-int^ fraction was found to correspond to Mo-derived MFs, expressing a MF transcriptomic signature and harboring meagre antigen presentation properties. The MHC-II^hi^ fraction was claimed to correspond mainly to cDCs that are able to efficiently process and present antigens for cognate activation and expansion of T cells^18^. Yet, only a minor fraction of these MHC-II^hi^ cells is fate-mapped by *Clec9a^Cre^* and hence traced as being derived from the CDP. In addition, their precise nature remained unknown regarding whether they correspond to cDC2As, cDC2Bs or DC3s. Thus, considering the heterogeneity of total BM cell suspensions, methods are required to specifically track the fate of Mos or Ly6C^+^ MDPs in these cultures and to differentiate their progeny from that of the CDP. Moreover, to properly mirror the conditions the most frequently used for the *in vitro* differentiation of human moDCs, murine BM cultures should not be performed with GM-CSF only, but with IL-4 as well. This is all the more relevant as IL-4 has been demonstrated to be essential for the complete and functional differentiation of both human and mouse moDCs^13,33^. Thus, to ensure rigorous identification of murine *in vitro*-derived moDCs and their discrimination from CDP-derived DCs, we harnessed the recently developed mouse strain *Ms4a3^Cre^*;Ai14 that specifically fate-maps the GMP-derived cells^10^. We crossed it to *Zbtb46^GFP^* mice, whose reporter gene is specifically expressed in DCs within the hematopoietic lineage^75,76^, to generate MAZ mice. As a result, should moDCs develop in vitro, we would be able to identify them as co-expressing Ai14 and GFP (**Figure 1A**) (**Supplementary Figures 1-2**).

**Figure 1.**
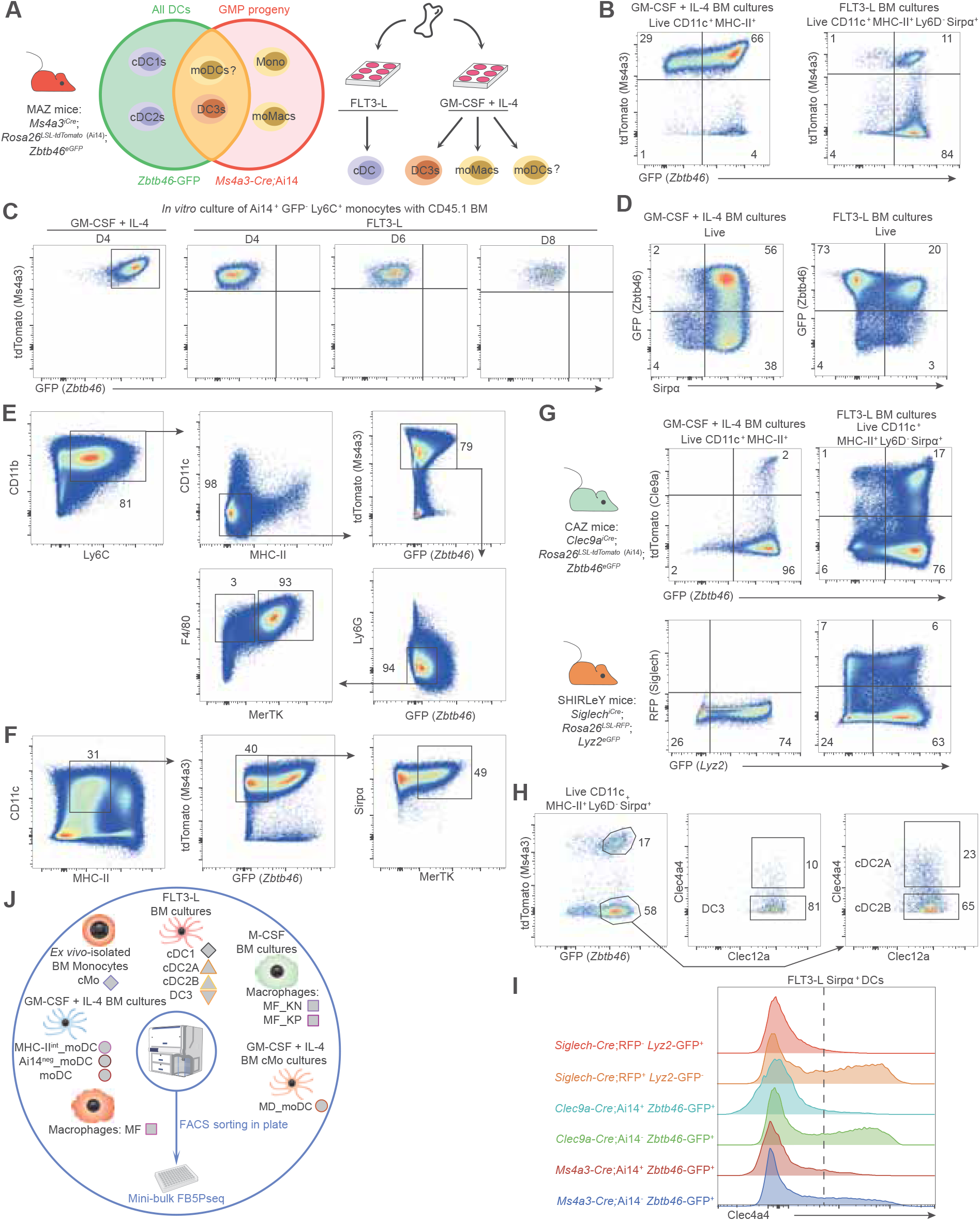
Combining *Ms4a3-Cre*;Ai14;*Zbtb46-*GFP mice with various *in vitro* differentiation systems allows to rigorously identify and compare moDCs, monocyte-derived macrophages, DC3s, cDC2Bs, cDC2As and cDC1s. **A)** Graphical scheme depicting how *Ms4a3-Cre*;Ai14;*Zbtb46-*GFP (MAZ) mice enable to identify all DCs as *Zbtb46*GFP^+^ cells, and GMP- or Ly6C^+^ MDP-derived cells as *Ms4a3-*Ai14^+^, such that Ai14^+^ GFP^+^ cells should encompass both DC3s and moDCs. Thus, harnessing MAZ mice in combination with cultures of total BM, or of purified BM monocytes, supplemented with either FLT3-L or GM-CSF and IL-4, should enable to generate and rigorously identify moDCs, monocyte-derived macrophages (MFs), CDP-derived cDC2Bs and cDC2As, as well as DC3s. **B)** Relative abundance of cells expressing either Ai14 or GFP, or co-expressing both, in total MAZ BM cultures differentiated upon exposure to GM-CSF and IL-4 versus to FLT3-L. Total BM cells from MAZ mice were cultured for 8 days in the presence of GM-CSF and IL-4 or of FLT3-L. Plots displaying the relative expression of tdTomato (Ai14) versus Zbtb46 were made by pre-gating on Live, CD11c^+^ MHC-II^+^ cells for GM-CSF and IL-4 cultures, and on Live, CD11c^+^ MHC-II^+^ Ly6D^-^ Sirpα^+^ DCs for FLT3-L cultures. **C)** *In vitro* culture of *ex vivo*-isolated BM Ai14^+^ GFP^-^ Ly6C^+^ monocytes from CD45.2^+^ mice cultured with supportive BM from CD45.1 congenic mice, in the presence of either GM-CSF and IL-4 or FL3T-L. Monocytes were cultured with GM-CSF and IL-4 for 4 days, whereas they were cultured in the presence of FLT3-L for different durations, namely for 4, 6 and 10 days. For cultures supplemented with GM-CSF and IL-4, the progeny of the purified monocytes was pre-gated as Live, CD45.1^-^, Ai14^+^, CD11c^+^ and MHC-II^+^. For FLT3-L cultures, they were pre-gated as Live, CD45.1^-^ Ai14^+^. The plots display the tdTomato versus GFP signal at each timepoint. **D)** Sirpα and *Zbtb46-*GFP expression on all live cells in day 8 cultures of total BM from MAZ mice with either GM-CSF and IL-4 or FLT3-L. **E)** Gating strategy to identify F4/80^+^ MerTK^+^ and F4/80^+^ MerTK^-^ macrophages differentiated in 6-day cultures of total BM from MAZ mice supplemented with M-CSF. **F)** Gating strategy to identify MertK^+^ macrophages differentiated in 8-day cultures of total BM from MAZ mice with GM-CSF and IL-4. **G)** Relative abundance and Ai14-, RFP- and GFP-expressing cells differentiated in cultures of total BM from *Clec9a*-Cre;*Ai14*;*Zbtb46*-GFP (CAZ) mice, or from *Siglech*-iCre;*Rosa26-LSL-RFP*;*Lyz2*-GFP (SHIRLeY) mice, in the presence of GM-CSF and IL-4 versus FLT3-L. Total murine BM cells were cultured for 6 days in the presence of GM-CSF and IL-4, or for 8 days in the presence of FLT3-L. Plots displaying the relative expression of red fluorescent proteins (Ai14 or RFP) versus GFP (reporting *Zbtb46* versus *Lyz2* expression) were made by pre-gating on Live, CD11c^+^ MHC-II^+^ cells for GM-CSF and IL-4 cultures, and, on Live, CD11c^+^ MHC-II^+^ Ly6D^-^ Sirpα^+^ DCs for FLT3-L cultures. **H)** Gating strategy to discriminate cDC2As, cDC2Bs and DC3s from one another within the population of Sirpα^+^ DCs arising in the cultures of total BM from MAZ mice with FLT3-L. Total BM cells from MAZ mice were cultured for 8 days in the presence of FLT3-L. Cells were pre-gated as Live CD11c^+^ MHC-II^+^ XCR1^-^ Sirpα^+^. Histograms showing CLEC4A4 expression levels within fate-mapped and non-fate-mapped Live CD11c^+^ MHC-II^+^ Ly6D^-^ XCR1^-^ Sirpα^+^ DCs in day-8_FLT3-L cultures of total BM from *Ms4a3-Cre*;Ai14;*Zbtb46-*GFP mice, or *Clec9a-Cre*;Ai14;*Zbtb46-*GFP animals or *Siglech-iCre*;RFP*;Lyz2-*GFP mice. **I)** Graphical scheme depicting the strategy used to develop an *in vitro* atlas for moDCs, MFs, DC3s and all cDC types (cDC2As, cDC2Bs and cDC1s). The various *in vitro*-generated cell populations have been sorted to high purity in minute quantities in individual wells of 96-well plates (100 cells/well) and profiled by medium throughput, highly sensitive, RNA-sequencing (FB5P-seq flash minibulk protocol). In addition, ex vivo isolated *Ms4a3*;Ai14^+^ Ly6C^+^ BM monocytes were included in the analysis, as well as moDCs differentiated from these purified monocytes upon culture with GM-CSF and IL-4 (MD-moDCs).

Total BM cells from MAZ mice were thus cultured side-by-side in the presence of either FLT3-L or GM-CSF and IL-4 (**Figure 1A**). In GM-CSF and IL-4 total BM cultures, the majority of CD11c^+^ MHC-II^+^ cells was fate-mapped by *Ms4a3* and displayed high levels of *Zbtb46*^GFP^ (**Figure 1B**), thus suggesting that they correspond to moDCs, and not to cDC2s. In FLT3-L total BM cultures, a minor population of doubly labelled cells also developed, although most of the CD11c^+^ MHC-II^+^ Ly6D^-^ Sirpα^+^ cells were single positive for Zbtb46 (**Figure 1B**) (**Supplementary Figure 2**).

Since DC3s are partially fate-mapped by *Ms4a3*^20^, we could not exclude the fact that they may develop in the presence of GM-CSF and IL-4. To resolve this uncertainty, we cultured, with either GM-CSF and IL-4 or FLT3-L, BM Ai14^+^ Ly6C^+^ Mos purified from CD45.2^+^ MAZ mice, together with supportive total BM cells from WT CD45.1 congenic animals (**Figure 1C**). MDPs and pre-DCs were successfully excluded from the sorted BM Mos, by excluding cells expressing *Zbtb46*^GFP^ or c-Kit. Consistent with previously reported results^32,33^, purified BM Mos did differentiate into moDCs in the presence of GM-CSF and IL-4 (**Figure 1C**), but not in the FLT3-L condition (**Figure 1C**). Whereas all Zbtb46^+^ cells in the GM-CSF and IL-4 BM cultures were Sirpα^+^, the majority of those in the Flt3-L cultures were Sirpα^-^ (**Figure 1C**). Altogether, these results confirm that the Ai14^+^ GFP^+^ cells arising in the GM-CSF and IL-4 total BM cultures must be *bona fide* moDCs, whereas the Ai14^+^ GFP^+^ cells differentiating in the FLT3-L total BM cultures should correspond to DC3s and the majority of the cells differentiating in these cultures are cDC1s.

To compare the cellular output of GM-CSF and IL-4 total BM to a reference MF population, we differentiated MFs from total BM cultures supplemented with M-CSF. These cells were negative for *Zbtb46*^GFP^, CD11c and MHC-II, while co-expressing F4/80 and MerTK (**Figure 1E**). We also identified MFs from the GM-CSF+IL-4 cultures as CD11c^+^ MHC-II^-/low^ Ai14^+^ GFP^-^ Sirpα^+^ MerTK^+^ cells (**Figure 1F**). To confirm that the DCs developing in GM-CSF and IL-4 BM cultures were not cDCs, we assessed their level of fate mapping in *Clec9a*^iCre^;Ai14;*Zbtb46*^GFP^ (CAZ) mice as compared to the DCs developing in FLT3-L cultures (**Figure 1G**). The fraction of Clec9a-fate mapped cells was extremely low within the CD11c^+^ MHC-II^+^ *Zbtb46*-GFP^+^ cells of the GM-CSF+IL-4 cultures (∼2%), around ten-times lower than what was observed in FLT3-L cultures (∼22%). cDC2As can be discriminated from cDC2Bs via Esam in the spleen. However, the expression of Esam is unreliable as it does not strictly reflect lineage identity but also varies with differentiation stages and it is not expressed in peripheral tissues^8,77^. cDC2As specifically express Clec4a4^37,78–80^, but its use remains limited in the field as many studies rely on ESAM expression or fate-mapper and reporter mouse models^8,36–38^. In addition, it remains elusive whether cDC2As can develop in FLT3-L total BM marrow cultures in the absence of OP9-DL1 cells that are necessary to induce *Notch2* signaling^25,81^. To screen for the development and phenotypic characterization of cDC2As in total BM cultures, we took advantage of our *Siglech^Cre^;*RFP;*Lyz2*^GFP^ (SHIRLeY) mouse model (**Supplementary Figure 1**) that allows to discriminate cDC2As from cDC2Bs based on their reciprocal expression of RFP versus GFP^8^. In FLT3-L total BM cultures from SHIRLeY mice, Clec4a4 expression was observed selectively on the cDC2As, defined as *Siglech*;RFP^+^ *Lyz2*^GFP-^ cells, and not on the cDC2Bs/DC3s defined reciprocally as *Siglech*;RFP^-^ *Lyz2*^GFP+^ cells (**Figure 1G-H**). Hence, in the FLT3-L total BM cultures from MAZ mice, within the mixed cDC2 population defined as *Ms4a3*^Cre^;Ai14^-^ *Zbtb46*^GFP+^, we could enrich cDC2As as Clec4a4^+^ and cDC2Bs as Clec4a4^-^ (**Figure 1H-I**). Yet, the differentiation of cDC2As appeared to be partly blunted in FLT3-L total BM cultures, since a *Siglech*;RFP^+^ *Lyz2*^GFP+^ fraction arose that is not naturally present in the spleen at steady state^8^, and which may correspond to a fraction of CX3CR1-expressing pre-cDC2As that has been reported by other groups^36–38^. In any case, these results suggest that cDC2As develop in FLT3-L total BM cultures in the absence of *Notch2* signaling and can be specifically identified by their Clec4a4 expression.

### Murine FLT3-L BM cultures faithfully generate cDC2 subsets, with cDC2As clearly differing from the transcriptionally identical cDC2Bs and DC3s

To validate our findings, we used an orthogonal approach for a deeper characterization of the *in vitro* generated DC types and the alignment with their *ex vivo* counterpart. Specifically, we profiled their gene expression, by mini-bulk RNA sequencing, based on an adaptation of SmartSeq2, termed FACS-Based 5-Prime End single-cell RNA sequencing (scRNA-seq) FLASH (FB5P-seq FLASH)^82^ (**Figure 1J**). We sorted different populations of MNPs generated through our *in vitro* culture pipeline from BM cells of MAZ mice: two MF populations generated in M-CSF cultures, *Ms4a3*-Ai14^+^ *Zbtb46*^GFP-^ MerTK^hi^ (KP_MFs) and *Ms4a3*-Ai14^+^ *Zbtb46*^GFP-^ MerTK^lo^ (KN_MFs) (**Figure 1E**); four DC populations arising in FLT3-L cultures, *Zbtb46*^GFP+^ cDC1s, *Ms4a3*-Ai14^-^ *Zbtb46*^GFP+^ Clec4a4^+^ Clec12a^+^ cDC2As, *Ms4a3*-Ai14^-^ *Zbtb46*^GFP+^ Clec4a4^-^ Clec12a^+^ cDC2Bs and *Ms4a3*-Ai14^+^ *Zbtb46*^GFP+^ Clec4a4^-^ Clec12a^+^ DC3s (**Figure 1H**); 4 populations developing in BM cultures in GM-CSF and IL-4, *Ms4a3*-Ai14^+^ *Zbtb46*^GFP+^ moDCs, *Ms4a3*-Ai14^-^ GFP^+^ moDCs (**Figure 1B)**, *Ms4a3*-Ai14^+^ *Zbtb46*^GFP+^ MHC-II^int^ pre-moDCs and *Ms4a3*-Ai14^+^ *Zbtb46*^GFP-^ MerTK^+^ MFs (MFs) (**Figure 1F**); *Ms4a3*-Ai14^+^ *Zbtb46*^GFP-^ Ly6C^+^ Mos sorted from the BM of MAZ mice (**Supplementary Figures 2-3**); and Ai14^+^ GFP^+^ moDCs generated from Ai14^+^ GFP^-^ Ly6C^+^ Mos sorted from the BM of MAZ mice and cultured with GM-CSF and IL-4, in the presence of supportive BM cells from congenic CD45.1^+^ C57BL/6 mice (**Figure 1C**). We computed all the gene expression data and clustered the different samples via uniform manifold approximation mapping (UMAP) (**Figure 2A**).

**Figure 2.**
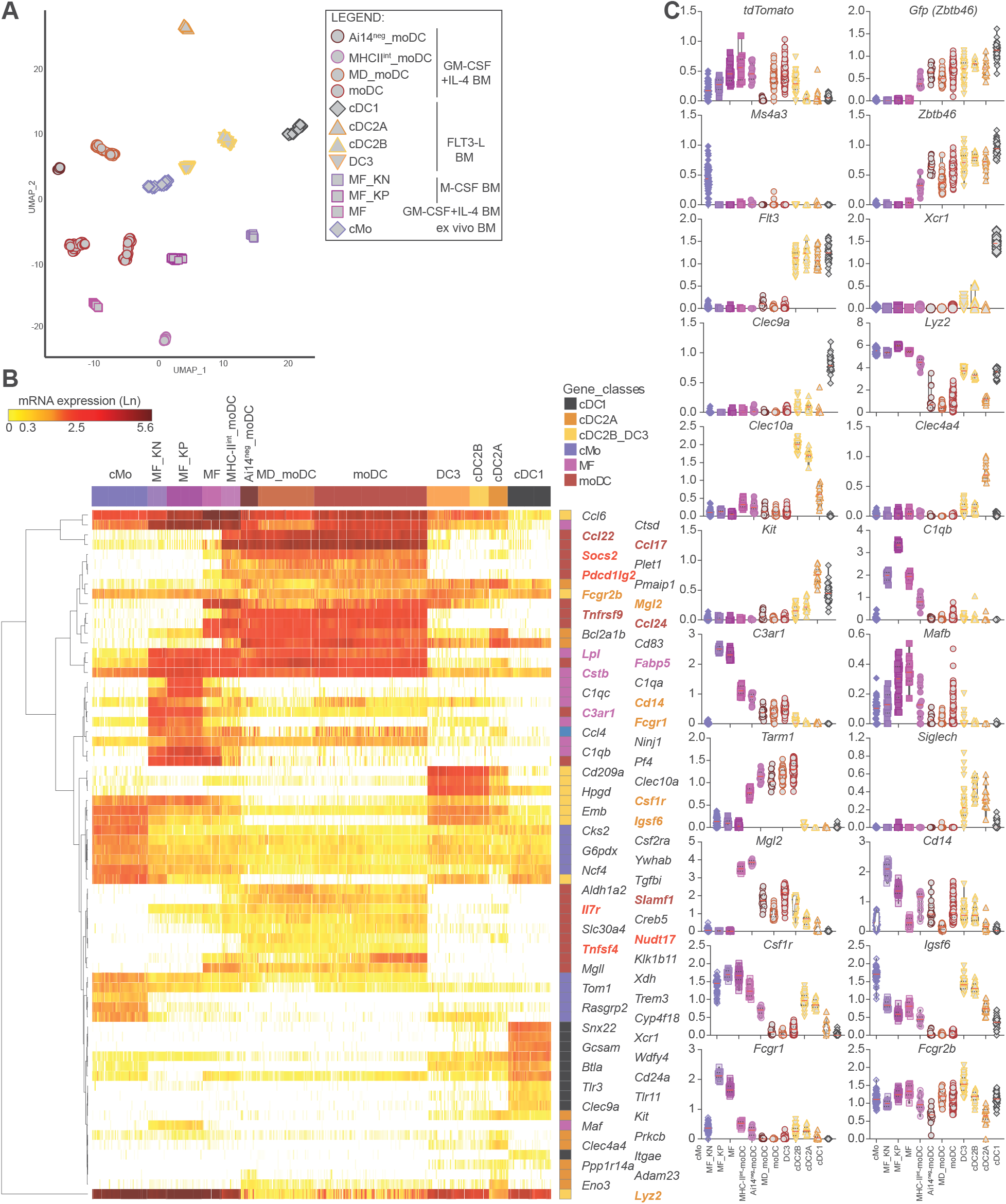
moDCs share genes with monocytes and macrophages and with cDCs, while displaying a unique transcriptome. **A)** UMAP representation of all the cells comprising the in vitro DC and macrophage atlas. **B)** Violin plots showing normalized RNA expression levels of selected genes across the various populations (in Ln values). **C)** Heatmap of selected genes that had been previously reported to be specifically expressed by splenic cDC1s, cDC2As, cDC2Bs/DC3s and macrophages when compared to one another in single cell RNA sequencing experiments^37^, or by in vitro-derived moDCs when compared to BM monocytes and to spleen cDC1s and cDC2s as assessed by microarrays^33^.

moDCs derived from total BM cultures and moDCs differentiated from FACS-sorted *Ms4a3*-Ai14^+^ Ly6C^+^ Mos (MD_moDC) clustered close to each other (**Figure 2A**) and shared a core transcriptional signature while also expressing some genes shared with cDCs and other genes shared with Mos (**Figure 2B-C**). cDC2Bs and DC3s clustered together and away from from cDC2As and moDCs.

As expected, sorted DC populations expressed the t*dTomato* and *Gfp* transcripts in accordance to their protein expression of *Ms4a3*-Ai14 and *Zbtb46*^GFP^ (**Figure 2B**). Each of the cell subsets analyzed expressed the expected canonical marker genes, such as *Ms4a3* for Mos; *Zbtb46* for all DC types, *Flt3* for cDCs and DC3s, *Xcr1* and *Clec9a* for cDC1s; *Lyz2* and *Clec10a* for cDC2Bs and DC3s; *Clec4a4* and *Kit* for cDC2As; *C1qb* for MFs. moDCs and MFs shared the expression of *C3ar1* and *Mafb. Tarm1* was confirmed to be rather selectively expressed by moDCs (**Figure 2B**), as previously reported for both mouse and human cells^83^. *Siglech* was expressed at similar levels in cDC2Bs and cDC2As, above the levels observed in DC3s and other cell types, suggesting that a sizeable fraction of cDC2As fail to terminally differentiate and acquire Clec4a4 expression, leading to their contamination of the cDC2B population. This can indeed be observed in the FLT3-L BM cultures from SHIRLeY mice where a significant fraction of the *Siglech*^iCre^;RFP^+^ *Lyz2*^GFP-^ cDC2As does not express Clec4a4 (**Figure 1H**).

To further assess the faithfulness of our *in vitro*-derived cells as models of *ex vivo* isolated DCs, we statistically tested their relative expression of canonical gene signatures previously reported to characterize splenic Mos, MFs and DC types, using the BubbleGUM algorithm^84^. BubbleGUM enabled us to automatically perform all pairwise Gene Set Enrichment Analysis (GSEA) between the cell types studied, with the implementation of the required multiple testing correction, for the reference transcriptomic signatures retrieved from the only scRNA-seq analysis that has compared side-by-side murine cDC2As, cDC2Bs and DC3s^37^ to the best of our knowledge. Our *in vitro*-derived cDC1s, cDC2As and MF_KP were consistently and significantly enriched for the transcriptomic signature of their *ex vivo* counterparts across all of the pairwise comparisons tested, as highlighted by the green squares on the corresponding BubbleMaps (**Figure 3**). A similar enrichment was observed in the *in vitro*-derived DC3s for the signature shared by *ex vivo* isolated DC3s and cDC2Bs as compared to cDC2As and cDC1s, and for the signature specific to *ex vivo* isolated cDC2Bs as compared to DC3s and cDC2As although it did not reach statistical significance for the comparisons with DC3s or cDC1s. Many of the genes identified as more highly expressed in DC3s as compared to cDC2Bs are expressed in MFs, and, consistently, among the cell types derived *in vitro*, the corresponding signature was higher in DC3s than in cDC2As, cDC2Bs and cDC1s, but lower than in MFs (**Figure 3**). For example, the expression of *Csf1r*, *Mgl2*, *Fcgr2b* and *Lyz2* was shared between *in vitro*-derived cDC2Bs/DC3s, MFs and moDCs (**Figure 2B-C**), emphasizing the difficulty to rigorously discriminate these cell types from one another when assessing their expression of only a few cell surface receptors or genes. The similarities between the BubbleMAPs for the comparisons of cDC2Bs and DC3s to all of the other MNP types examined confirms their strong transcriptomic relatedness and supports the hypothesis that they correspond to the same cell type that could differentiate along two different but converging ontogenetic paths, as already reported for example for cDC2As^5^. The expression pattern of some of the genes contributing the most to the GSEA results are illustrated as a heatmap (**Figure 2B**). Overall, these analyses confirm the identities of the *in vitro* derived cell types and their equivalency with the MNPs that can be directly isolated *ex vivo* from the murine spleen, hence demonstrating that the murine FLT3-L BM cultures we used faithfully generate the different types of DC2s, with cDC2As clearly differing from the transcriptionally similar cDC2Bs and DC3s. Hence, our atlas should enable for the first time to rigorously compare side-by-side murine moDCs and cDC2Bs/DC3s to identify their specific transcriptomic and phenotypic identities.

**Figure 3.**
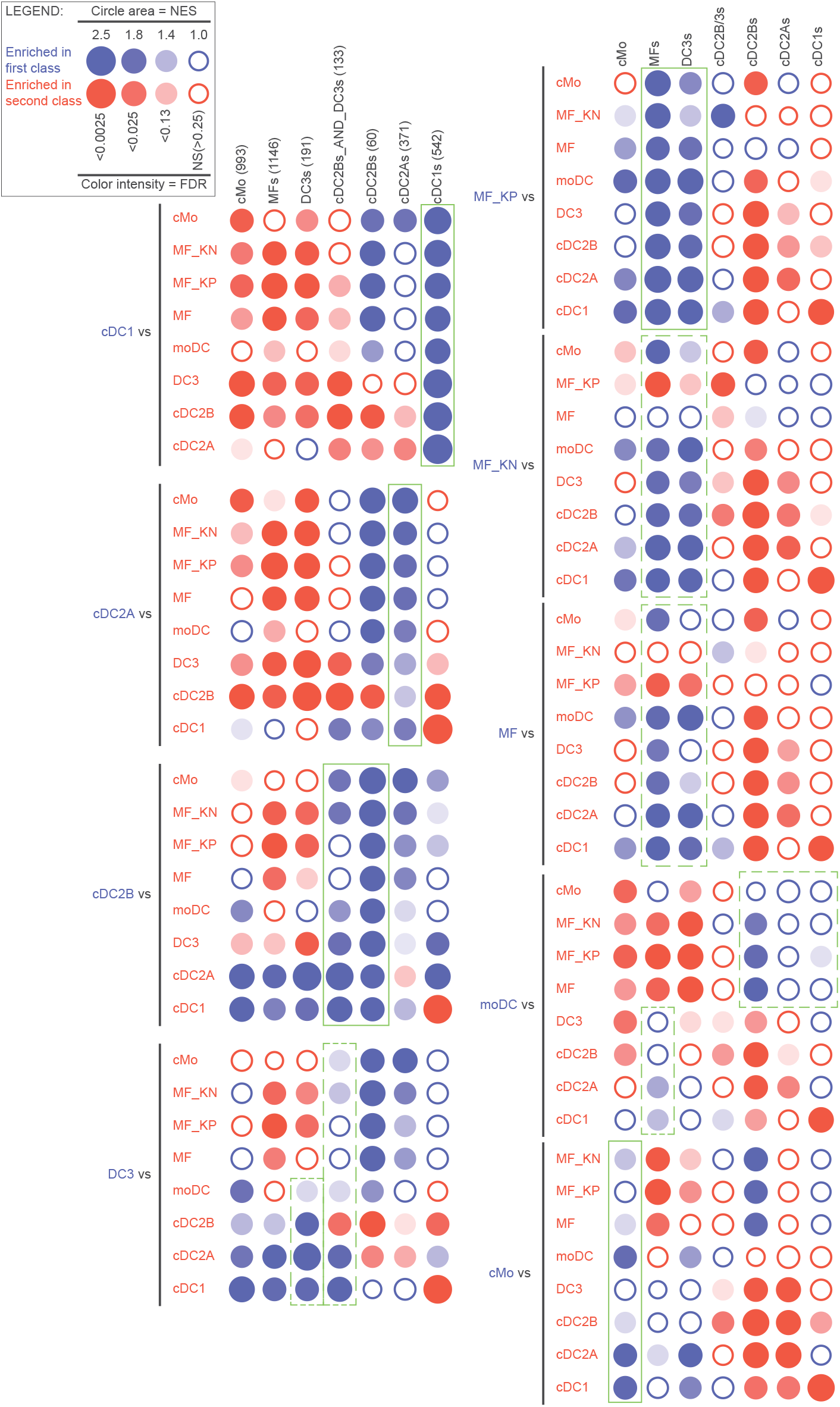
*In vitro*-generated cells are enriched for the transcriptomic signatures of their *ex vivo*-isolated counterparts. moDCs are characterized by a mixed expression of macrophages and DC genes. High throughput GSEA was performed on pairwise comparisons between in vitro-derived cells (rows), for enrichment of the transcriptomic signatures defined by single cell RNA sequencing analysis of murine splenic MHC-II-positive cells^37^ (columns), by using the BubbleGUM algorithm. Vertical green squares highlight consistent enrichment across all pairwise comparisons of the signature for an *ex vivo*-isolated cell type in its *in vitro*-generated counterpart (e.g. upper left, BubbleMAP, ex vivo cDC1 signature enrichment in *in vitro*-generated cDC1), even though these enrichments do not always reach the FDR threshold for statistical significance (empty bubbles, e.g. bottom left BubbleMAP, *ex vivo* splenic cDC2B signature enrichment in in vitro-derived cDC2Bs as compared to MF). Vertical dotted green squares highlight relative enrichments of interest, e.g. enrichment of the transcriptomic signature for *ex vivo*-isolated macrophages in *in vitro*-generated moDCs when compared to other *in vitro*-generated DCs, and enrichment of the transcriptomic signatures of *ex vivo*-isolated DCs in *in vitro*-generated moDCs when compared to monocytes or macrophages.

### Murine moDCs harbor a dual identity, being enriched for DC gene signatures when compared to MFs but for MF gene signatures when compared to DCs

Our GSEA analyses showed that *in vitro*-derived murine moDCs were enriched for DC gene signatures when compared to MFs, but for MF gene signatures when compared to DCs, as indicated by the dotted green rectangles on the corresponding BubbleMap (**Figure 3**). For example, i*n vitro*-derived moDCs shared with MFs the expression of certain of their canonical signature genes including *C3ar1*, *Cstb*, *Fabp5* and *Lpl*. To further characterize the molecular fingerprints of our *in vitro*-derived murine moDCs, we examined their expression of genes that were independently identified as specific to murine moDCs differentiated *in vitro* from FACS-sorted Mos cultured with GM-CSF and IL-4, when compared to BM Mos and to spleen cDC1s and cDC2s, as assessed by microarrays^33^. Many of these genes were also uniquely expressed by moDCs in our dataset when compared to cDCs, Mos and MFs, including *Ccl17*, *Ccl22*, *Ccl24*, *Tnfrsf9* and *Slamf1* (**Figure 2C**). The induction of several of these genes is known to occur during DC maturation, including for *Nudt17*, *Tnfsf4*, *Il7r*, *Pdcd1lg2* and *Socs2*. Hence, we wondered to which extent the genes identified as specific to moDCs could in part be linked to the specific activation state of these cells, including their response to the GM-CSF and IL-4 cytokines, rather than being a hallmark of their ontogenetic identity.

### Exposing the various DC types to different stimulation conditions allows to separate cell-type-specific core invariant gene modules from modules associated to activation/maturation states

To evaluate whether the moDCs generated *in vitro* in the presence of the proinflammatory cytokines GM-CSF and IL-4 harbored a more mature state than the DCs developing under FLT3-L instruction, we examined whether they expressed differentially gene modules linked to DC maturation^2^.To this aim, we performed pairwise GSEA using BubbleGUM between moDC and each of the other 3 *in vitro*-derived DC types. In each of these comparison, the moDCs were enriched for the expression of the gene signatures induced in mature DCs (**Supplementary file 1**), including a module previously reported to be selectively expressed by the “mature DCs enriched in immunoregulatory molecules” (mregDCs)^85^ (**Figure 4A**). Out of the 625 genes constituting the mregDC module, 253 were always enriched in moDCs when compared to each of the other 3 types of DCs generated *in vitro* in FLT3L BM cultures. The genes contributing the most to this enrichment included genes known to control the quality of the maturation of the DCs and the functional outcome of their interactions with T cells, such as *Ccr7*, *Cd40*, *Cd80*, *Cd83*, *Cd274*, *Pdcd1lg2*, *Cd200*, *Tnfrsf9*, *Icam1*, *Ccl5*, *Ccl22*, *Il15*, *Il7r*, *Stat4*, *Nfkbib*, *Traf1*, *Socs2*, *Etv3* and *Mir155hg* (**Supplementary file 1**). In contrast, the DCs generated in FLT3L BM cultures were enriched in genes characteristic of immature DCs, including cell cycle genes (**Supplementary file 1**). Thus, in our experimental conditions, the DCs generated in total BM FLT3-L cultures were in an immature state, whereas the moDCs generated in the total BM cultures with GM-CSF and IL-4 were highly activated, in a terminally mature “mregDC” cell state. This observation thus called for a strategy enabling to refine the identification of a core transcriptomic gene signature that would enable a robust discrimination of moDCs from other DC types irrespective of their activation state. Hence, to compare moDCs to other DC types in a similar maturation state, and also to identify genes that would be similarly regulated by IL-4 or GM-CSF across multiple DC types, we completed our atlas by profiling the gene expression of *in vitro* generated DC types exposed to a few selected maturation stimuli, IFN-γ and polyI:C for all DC types, and IL-4, GM-CSF and their combination for the FLT3-L-derived DCs (**Figure 4B**). This strategy should help overcome one of the main confounding effects currently in the field that is the induction of shared cell surface molecules on moDCs and cDC2Bs/DC3 under various inflammatory conditions^21,59,86^.

**Figure 4.**
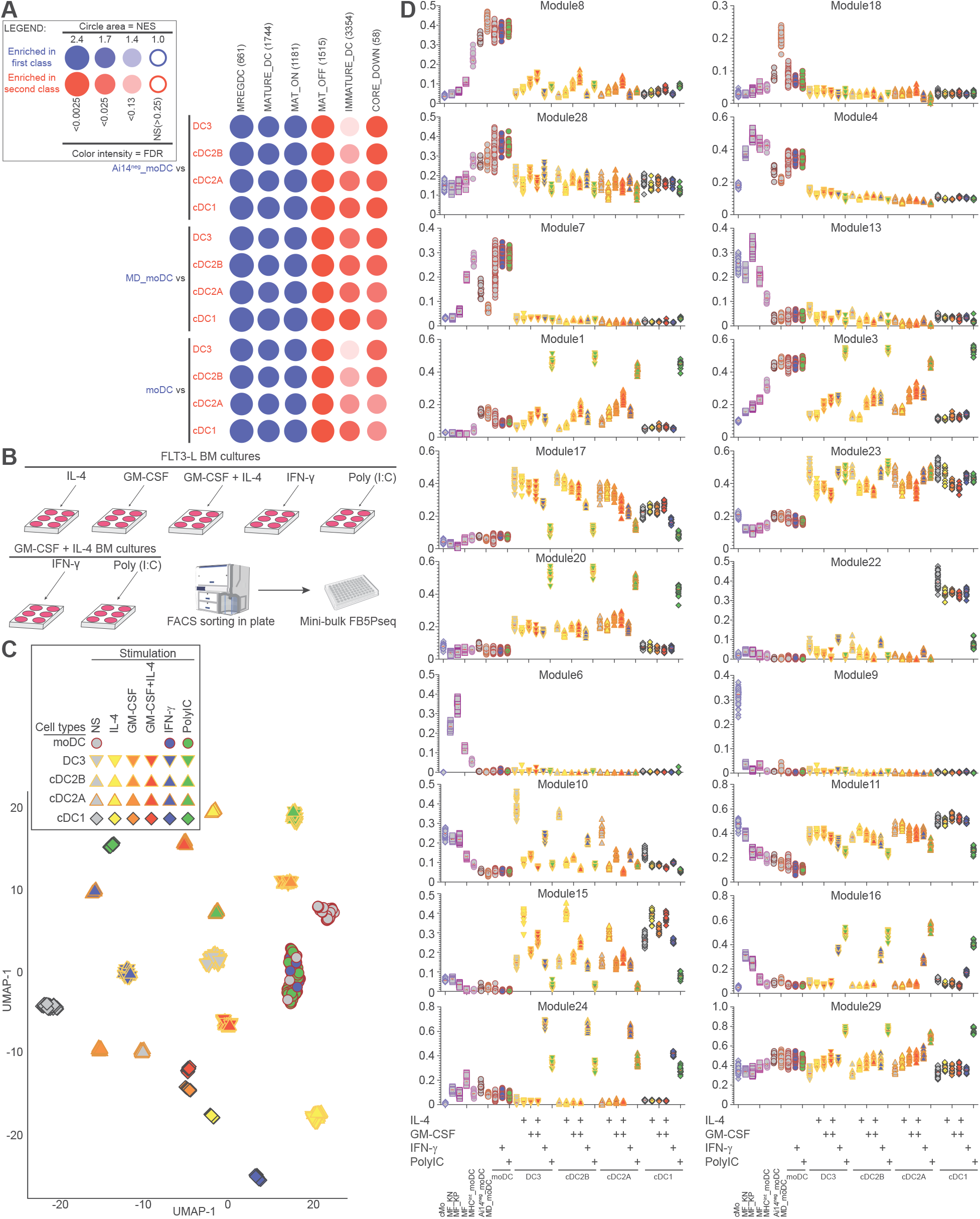
The refined *in vitro* atlas, including DCs matured under various stimulation conditions, enables to refine the transcriptomic identity of moDCs. **A)** GSEA enrichment plots from BubbleGUM showing that the unstimulated (NS) moDCs generated in total BM cultures with GM-CSF and IL-4 are significantly enriched for the “mregDC” gene module characteristic of terminally mature DCs, as compared to the unstimulated DC types generated in FLT3-L total BM cultures. **B)** Scheme depicting the strategy employed to stimulate overnight FLT3-L-derived cDC1s, cDC2As, cDC2Bs and DC3s at day 9 of culture with either IL-4, GM-CSF, GM-CSF and IL-4, IFN-γ, or Poly (I:C). moDCs from GM-CSF and IL-4 cultures were stimulated overnight at day 5 with either IFN-γ or Poly (I:C). Afterwards, each cell population was sorted in plates to perform FB5P-seq FLASH. **C)** UMAP representation of the various DC populations under stimulation conditions. **D-G)** Violin plots showing the normalized expression across cell types of several modules of genes co-expressed across the entire atlas as identified with the SciGeneX algorithm, including moDC-specific gene modules (**D**), modules expressed in moDCs over monocytes, macrophages and NS FLT3-L DCs but induced in the later under certain maturation conditions (**E**), modules selectively expressed in both moDCs and monocytes or macrophages (**F**), or gene modules selectively expressed in FLT3-L DCs eventually depending on their stimulation conditions (**G**).

The samples of the resulting expanded atlas clustered in the UMAP space in a manner dependent on both the cell type and stimulation condition (**Figure 4C**). The various FLT3-L-derived DCs clustered separately for each type of stimulation, with the exception of cDC2Bs and DC3s, that co-clustered in every stimulation condition. This observation further supports that cDC2Bs and DC3s may correspond to the same DC type, although bearing a different developmental origin. Contrary to FLT3-L-derived DCs, stimulated moDCs clustered together with their unstimulated counterparts, suggesting that the combination of GM-CSF and IL-4 may induce by itself an irreversible program of terminal maturation on moDCs that is refractory to most of the changes induced by poly (I:C) or IFN-γ in the other DC types. Following integration of two mini-bulk datasets, we utilized the algorithm termed Single-cell informative Gene eXplorer (SciGeneX)^87^ in order to identify the major core modules of co-expressed genes associated to the overall biological variability in this dataset. This unbiased, global approach identified several moDC-specific modules which encompassed genes related to the regulation of T cell activation (Module 7) and to the differentiation and activation of myeloid cells (Module 17) (**Figure 4D, Supplementary file 2**). Two of the modules encompassing the highest number of coregulated genes were associated to DC maturation states shared between moDCs and FLT3L-derived DCs stimulated with polyI:C or GM-CSF and IL-4. These modules included genes related to lymphocyte differentiation (Module 1) or induced in response to LPS (Module 3) (**Figure 4D, Supplementary file 2**). Furthermore, we identified conserved myeloid modules shared between moDCs, Mos and MFs, including for instance genes associated to TNF production and TNFR1 signaling (Module 4) or linked to response against bacteria and leukocyte migration (Module 8) (**Figure 4D, Supplementary file 2**). Finally, we identified FLT3-L-derived DC-specific modules comprising genes linked to cellular biosynthesis and cytoplasmic translation machinery (Modules 18 and 23) (**Figure 4D, Supplementary file 2**). We then computed some of the most representative genes from the modules linked to cell type identities for illustration as a heatmap (**Figure 5A**). As compared to all other examined combination of cell types and activation states, moDCs uniquely expressed many genes to high levels, including *Slamf1*, *Dcstamp*, *Slc7a11*, *Creb5*, *Nos2* and *Tarm1*. They shared with mature FLT3-L DCs the expression of activation genes involved in the cross-talk with T cells, including *Cd40*, *Fscn1*, *Etv3* and *Relb, and with* MFs genes involved in innate immune recognition of danger signals or in lipid metabolism, including *C3ar1, Chil3, Trem2*, *Cd36,* and *Lpl*. FLT3-L-derived DCs but not moDCs specifically expressed canonical cDC genes, corresponding to module 23 (**Figure 4G**) and including *Flt3*, *Btla* and *Dpp4* (**Figure 5A, Supplementary file 2**). We also identified genes specifically expressed in MFs, corresponding to module 6 and including *C1q*a, *C1qb*, *C1qc*, *Siglec1*, *C5ar1* or *Ccl7*, or in Mos, corresponding to module 11 and including *Elane*, *Hp*, *Fn1*, *Mpo*, *S100a8* and *S100a9* (**Figure 5A, Supplementary file 2**). Hence, these results confirmed the dual/Janus nature of moDCs, which share one part of their molecular identity specifically with other DCs, linked to the machinery necessary for professional antigen processing and presentation and the downstream activation and functional polarization of T cells, and another part of their molecular identity with MFs linked to antimicrobial, inflammatory and metabolic activities. They also enabled the identification of a list of genes selectively expressed in moDCs as compared to Mos, MFs or cDC2s/DC3s, and reciprocally, list of genes expressed in these MNP types but not in moDCs. Hence, the combination of these gene signatures should facilitate robust identification of moDCs *in vivo* upon bioinformatics analyses of scRNA-seq data, including their rigorous discrimination from activated Mos, MFs as well as cDC2Bs/DC3s.

**Figure 5.**
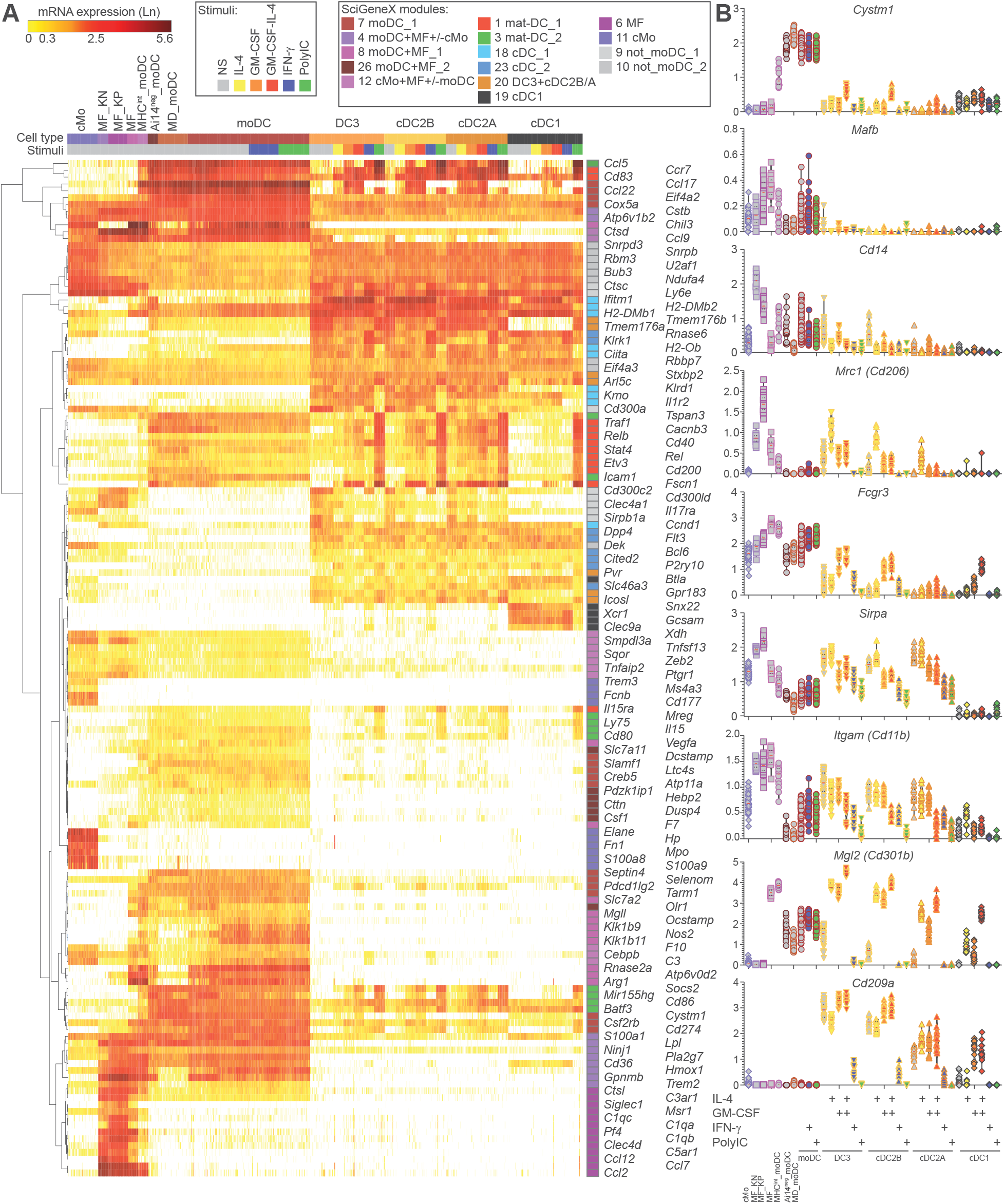
Stimulated FLT3-L-derived DCs and moDCs share the expression of common activation genes, while also expressing unique gene modules. **A)** Heatmap of selected genes representative of some of the identified core differentially expressed genes obtained from the two merged mini-bulk experiments, one with various unstimulated cell types and another with moDCs and FLT3-L-derived DCs under stimulation conditions. **B)** Violin plots showing the expression of selected genes across cell types and stimulation conditions.

We then proceeded to identify genes whose expression could be altered in the various stimulation conditions. We found that some genes considered specific of the Mo/MF lineage were upregulated by cDC2Bs/DC3s and cDC1s upon stimulation (**Figure 5B**). In particular, cDC2Bs/DC3s upregulated *Fcgr3* in response to IL-4 alone or GM-CSF and IL-4, while cDC1s upregulated this gene in response to GM-CSF and IL-4. Similarly, *Mgl2* was induced in all Sirpα^+^ FLT3-L-derived DCs in response to GM-CSF, IL-4, or GM-CSF and IL-4, while it was induced in cDC1s only by the combination of GM-CSF and IL-4. *Mrc1* was mainly expressed by MFs, and showed low expression levels in non-stimulated FLT3-L-derived DCs and moDCs. However, the addition of GM-CSF to FLT3-L cultures resulted in *Mrc1* upregulation in cDC2Bs/DC3s. Other canonical DC genes of Sirpα^+^ FLT3-L-derived DCs were downregulated in some of the stimulation conditions. Sirpα^+^ FLT3-L-derived DCs tended to express *Itgam* and *CD209a*. Nevertheless, stimulation with IFN-γ or Poly (I:C) led to a strong decrease of these genes, suggesting that the phenotype of activated DCs may differ depending on the maturation stimuli. We further observed that unstimulated DC3s and cDC2Bs expressed some levels of *Cd14* in the non-stimulated condition, while it was downregulated upon stimulation. *Cd14* was also expressed by both *in vitro*-derived moDCs and MFs. In addition, we found that *Mafb* expression was restricted to moDCs, Mos and MFs but not FLT3-L-derived DCs thus making it a valuable candidate for targeting cells of the Mo lineage.

Afterwards, we identified the most representative genes induced across the various DC types that were unique to each maturation condition. moDCs did not undergo notable gene expression changes in response to Poly (I:C) or IFN-γ (**Figure 6A**). However, FLT3-L DCs underwent gene expression changes that were shared across the various stimulation conditions, but also displayed specific gene patterns uniquely induced by each maturation cue (**Figure 6A**). For instance, stimulation with Poly (I:C) uniquely induced the upregulation of *Il1b, IL15ra, and Il2ra* across cDC1s, cDC2As, cDC2Bs and DC3s, suggesting that these genes are activated via common pathway shared across all cDCs (**Figure 6B**). Conversely, *Clxcl9* was uniquely induced across all cDCs by exposure to IFN-γ, while both poly (I:C) and IFN-γ stimulation led to an increase in *Il12b* and *Cxcl10* expression by cDC2s and DC3s. This highlights that, *in vivo*, unique microenvironments, enriched in specific maturation cues, can radically shape DC function by steering specific gene expression programs and that this is for the most part not dependent on the specific DC type. However, we still observed some differences that were subset specific, as cDC1s failed to upregulate *Il12b* or *Cxcl10* in response to IFN-γ as compared to cDC2s and DC3s, and expressed higher levels *Il27* (**Figure 6B**).

**Figure 6.**
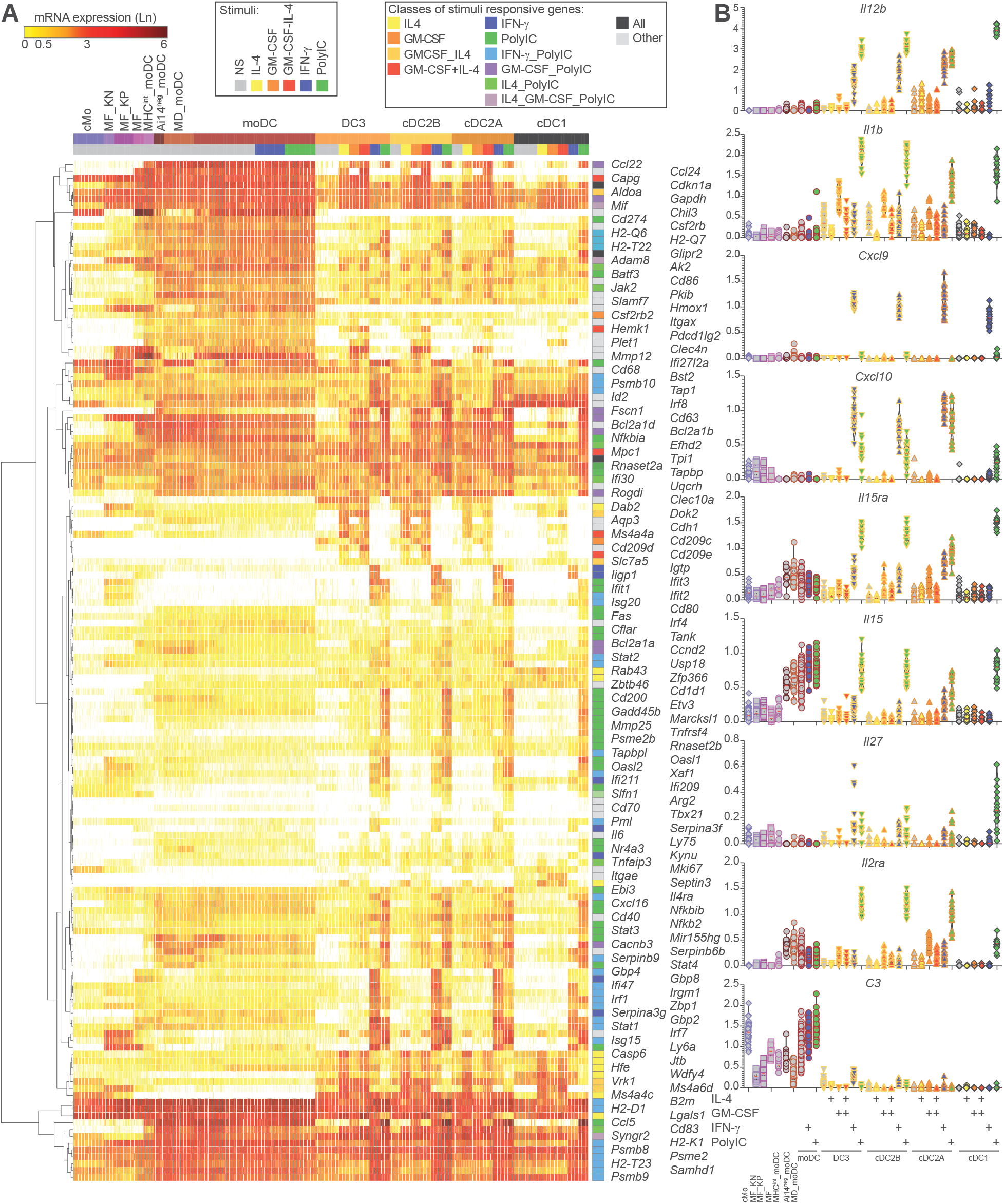
FLT3-L-derived DCs undergo common gene expression changes in response to various maturation stimuli, induced uniquely by each maturation cue. **A)** Heatmap of selected genes representative of some of the identified genes uniquely induced upon DC maturation by each specific stimulus, obtained from the two merged mini-bulk experiments, one with various unstimulated cell types and another with moDCs and FLT3-L-derived DCs under stimulation conditions. **B)** Violin plots showing the expression of selected genes uniquely induced by some stimulation conditions.

### Murine moDCs share specific gene expression patterns with human moDCs

We have generated murine *in vitro*-derived moDCs with high confidence and demonstrated that these cells are different from *in vitro*-derived cDCs/DC3s and MFs. We wondered how this *bona fide* population of moDCs, differentiated in the presence of GM-CSF and IL-4, would compare to human moDCs derived from purified Mos, also exposed to GM-CSF and IL-4. We gathered a sub-dataset of the microarray compendium developed by the teams of David A. Hume and Tom C. Freeman including blood and tonsil cDC2s, alveolar MFs, classical Mos and *in vitro*-derived human moDCs and MFs^88^ and compared it to the murine MNP types identified in our mini-bulk *in vitro* atlas (**Figure 7**). Murine and human *in vitro*-derived moDCs shared some core signature genes including *Slamf1*, *Sdc1*, *Dcstamp* or *Arg1*, while sharing some common genes with human and mouse cDC2s including *Batf3*, *Etv3*, *Ly75*, *Irf4* or *Zbtb46*. Interestingly, human *in vitro*-derived moDCs expressed higher levels of some genes enriched in *in vitro*-derived MFs compared to murine moDCs such as *Msr1*, *C1qb*, *Spp1*, *Trem2*, *Gpnmb* suggesting a stronger Mo/MF imprinting in human-derived moDCs. Collectively, we have identified gene sets with a conserved expression pattern in human and mouse moDCs as compared to other MNPs, thus supporting the biological relevance of studying the identity and function of murine moDCs to understand the biology of their human counterparts.

**Figure 7.**
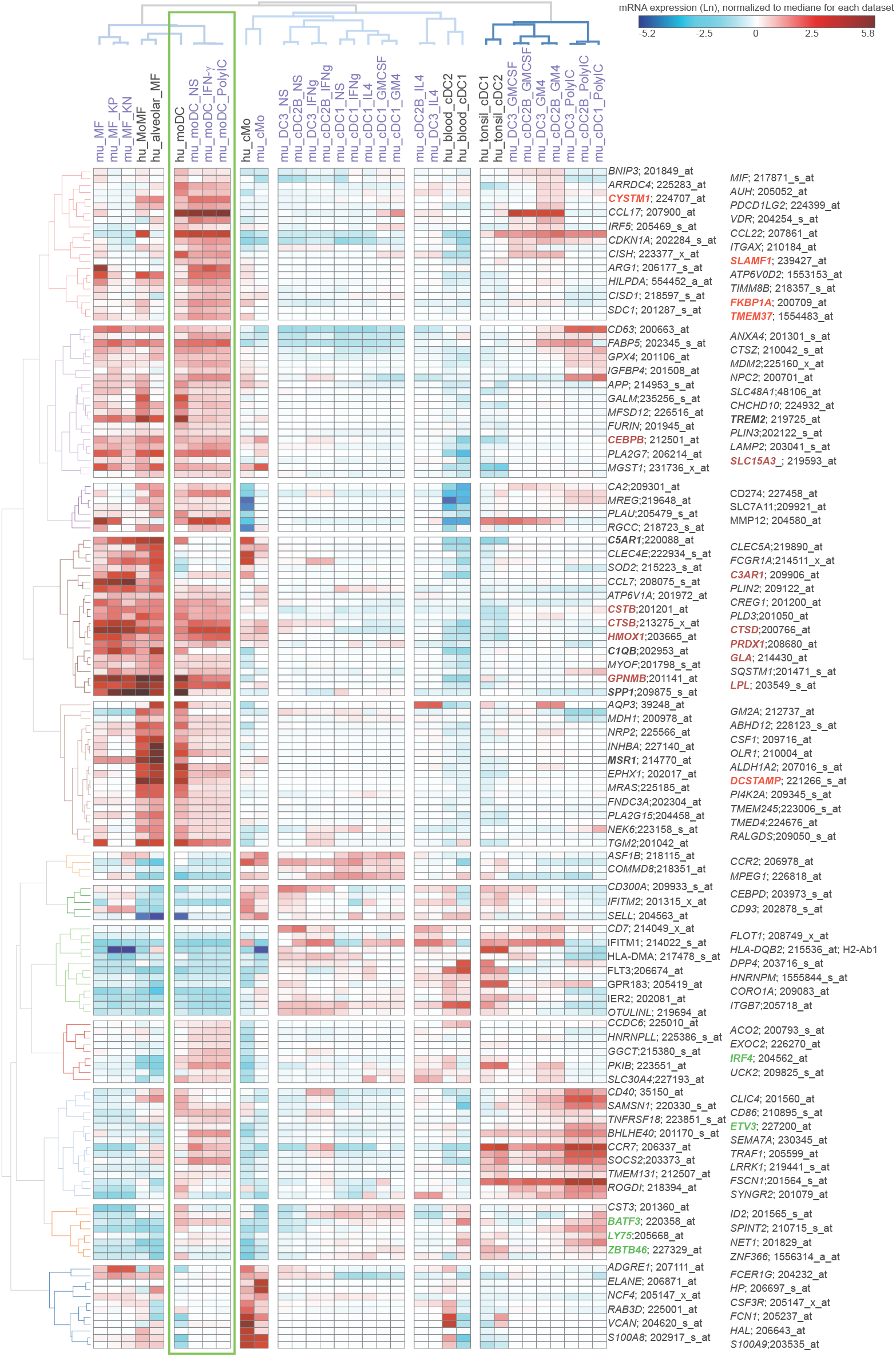
Murine and human *in vitro*-derived moDCs share the expression of conserved genes unique to the moDC lineage. A sub-dataset of the microarray compendium, developed by the teams of David A. Hume and Tom C. Freeman including blood and tonsil cDC2s, alveolar macrophages, classical monocytes and *in vitro*-derived human moDCs and macrophages, was integrated and compared to the transcriptomic data of murine DC and macrophages populations from the mini-bulk *in vitro* atlas.

### *In vitro*-derived moDCs elicit responses to TLR stimulation distinct from that of cDC2s and DC3s

We have shown that FLT3-L BM cultures enable to generate cDCs in large numbers, and that moDCs are the only DC type present in GM-CSF and IL-4 BM cultures. Thus, we wondered whether differences between cellular lineages and culture types would imprint the distinct DC types to respond differently to several TLR ligands, as we previously reported for human moDCs and cDC1s^64^. To test this, we stimulated overnight the *in vitro*-derived DC types, with poly (I:C), LPS and R848 (**Figure 8A**). moDCs showed an enhanced response to LPS stimulation and where by far the largest producers of IL-6 and TNF, whereas the other cDC types responded mildly (**Figure 8B**). In response to poly (I:C) stimulation, cDC1s were the only producers of cytokines, secreting in particular relatively high levels of IFN-β and also producing IL-6 and TNF, with the limitation of the cytokine panel measured. In response to Poly (I:C) stimulation, although all FLT3-L DC types expressed the *Il12b* gene encoding the IL-12p40 chain, none secreted substantial amounts of the bioactive IL-12p70 heterodimer, suggesting that additional signals are required for its production. Sirpα^+^ cDCs were the most responsive to R848, with DC3s producing the highest levels of IL-6 that was further amplified upon addition of IFN-γ (**Figure 8C**). cDC2Bs followed a similar pattern as DC3s, by producing IL-6 in response to R848, although this response was not increased upon addition of IFN-γ. cDC2As were the strongest producers of IL-10 upon R848 stimulation, and this response was diminished by the addition of IFN-γ (**Figure 8D**). These results support a division of labor between distinct DC types for preferential sensing of specific danger signals, e.g. LPS for moDCs, poly(I:C) for cDC1s and R848 for cDC2s/DC3s, and/or the production of different combinations of cytokines in response to the same stimulus sensed by all DCs, as it is the case for poly (I:C) and LPS, which induce TNF and IL-6 in all DC types but at various levels and together with different other cytokines. Although DC3s and cDC2Bs show similar responses to TLR ligands, DC3s seem to react more actively. Whether this is due to their different ontogeny will have to be evaluated in further studies. As reported by others^24^, cDC2As seem to be more pro-tolerogenic when compared to cDC2Bs and DC3s, as evidenced by their higher production of IL-10. We observed similar patterns of IL-6 and TNF production by *in vitro*-derived cDC2As and cDC2Bs, contrasting with the differences that have been reported between splenic cDC2As and cDC2Bs whereby the later were more potent producers of these proinflammatory cytokines^24^. This apparent discrepancy suggests that *in vitro*-derived cDC2Bs may not be terminally differentiated and have not acquired the full-blown functions of their *in vivo* counterparts.

**Figure 8.**
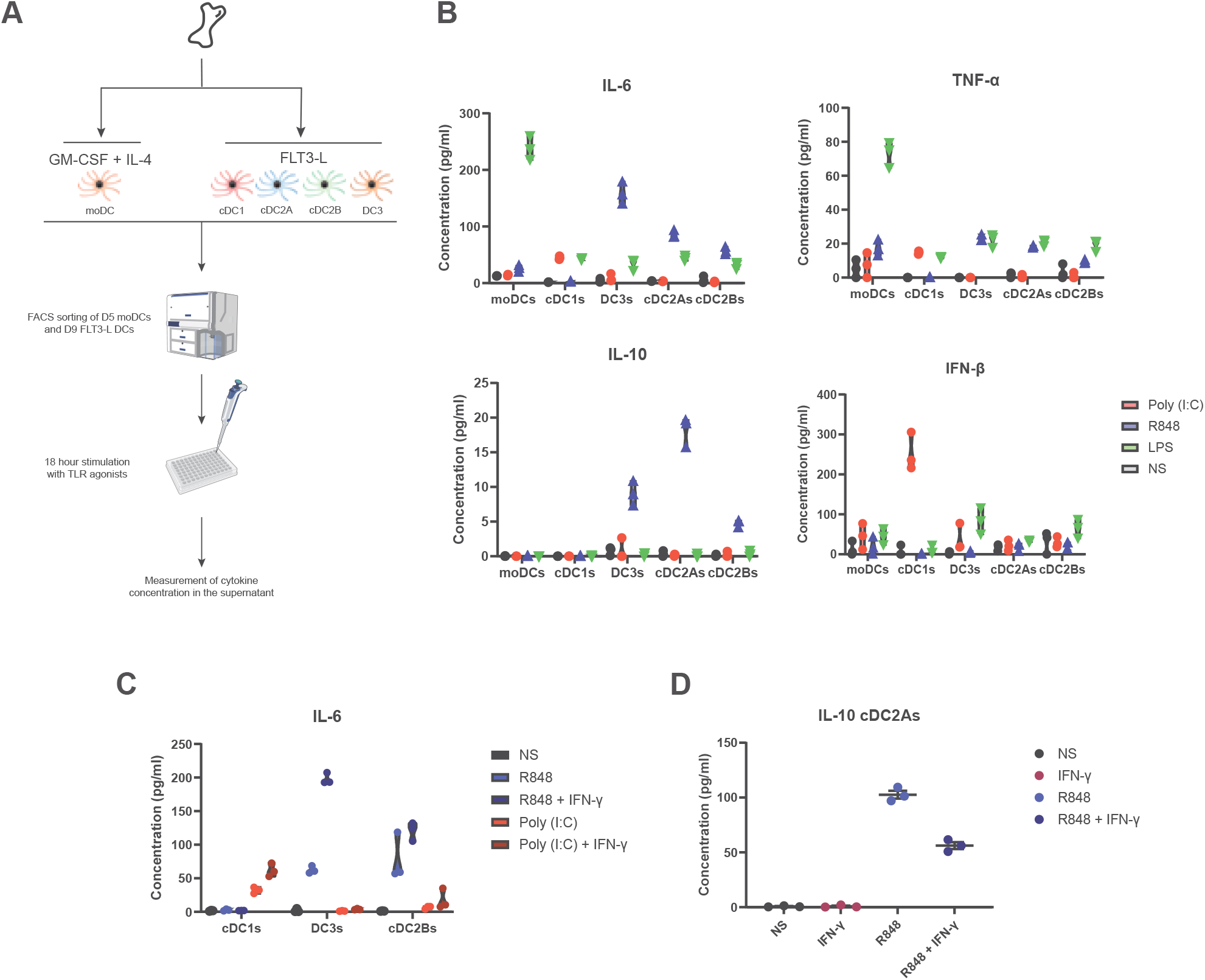
In vitro derived DC types respond differently to TLR stimulation. **A)** Graphical scheme depicting the workflow to measure cytokine production in response to TLR ligands by the various DC types. moDCs, cDC1s, cDC2As, cDC2Bs, and DC3s were produced using GM-CSF and IL-4 or FLT3-L BM cultures respectively. Day-5 moDCs and day-9 FLT3-L DCs were FACS sorted and 20000 (B) or 15000 (C) cells were plated in triplicate for each DC type and condition. DCs were stimulated overnight and the supernatant was collected 18 hours after stimulation for the titration of a restricted panel of cytokines by bead arrays. **B)** Graphs showing the concentration levels of IL-6, TNF-α, IL-10 and IFN-β in the supernatants of DC types purified from total BM cultures and stimulated with Poly (I:C), R848 or LPS. **C)** Graph showing the concentration of IL-6 secreted by cDC1s, DC3s or cDC2Bs when Poly (I:C) or R848 were combined with IFN-γ. **D)** Graph showing the concentration of IL-10 secreted by cDC2As in response to R848 alone or in combination with IFN-γ.

### Phenotypic characterization of MNP types *in vivo* in the lung, MedLN and peritoneum of MAZ mice

moDCs have been reported to develop in the murine model of allergy to house dust mite (HDM) induced upon sensitization in the lungs via repetitive intranasal instillation of the allergen^89,90^ and in the peritoneum upon local sterile inflammation induced by thioglycolate administration^91^. However, their phenotype largely overlaps with those of cDCs and MFs, to the point that it remains elusive whether moDCs actually develop in these settings or whether the cells that had been identified as moDCs corresponded to mis-identified activated Mos, MFs, cDC2Bs/DC3s or inf-cDC2s^20,21^. These two murine models of sterile tissue inflammation differ substantially. The lung microenvironment harbors an abundance of GM-CSF and IL-4/IL-13, further increased upon type 2 immunity induction^90,92,93^, that could potentially foster the development of moDCs closely resembling their *in vitro* counterparts. In contrast, thioglycolate-induced peritonitis is a model independent of GM-CSF, whereby the molecular makeup of moDCs could drastically differ from those found in HDM. To gain further rationale on moDC development in the two models, we adopted a multimodal approach. The major bottlenecks hampering the study of moDCs are the lack of reliable markers for their identification and the contamination with pre-DCs when performing adoptive transfer of Mos. With our in vitro atlas, we overcame both issues, since (i) we uncovered a series of differentially expressed genes unique to *in vitro*-derived moDCs, and (ii) we validated our sorting strategy to isolate pure GMP-derived cMos as *Ms4a3*-Ai14^+^ Ly6C^+^ *Zbtb46*^GFP-^ c-Kit^-^ cells as we did not observe any traces of genes associated to cDCs. For the peritonitis model, we performed adoptive transfer of Mos as this strategy yields optimal numbers of daughter cells^91^, thus enabling their in-depth characterization. For the HDM model, due to the technical difficulties of performing successful adoptive transfer of Mos to the allergic lungs, we opted to rely on cell surface markers to delineate putative populations of moDCs as compared to cDCs.

From the series of differentially expressed genes identified in our *in vitro* atlas, we selected candidates encoding cell surface molecules that may be useful to phenotypically distinguish moDCs from FLT3-L-derived cDC2s and DC3s (**Figure 9A**), in order to enable their *ex vivo* identification, characterization and purification by flow cytometry. The differential expression of the corresponding cell surface markers between *in vitro*-derived moDCs and cDC2s was validated by flow cytometry. Specifically, CD36 and BTLA harbored reciprocal expression between *in vitro*-derived moDCs (CD36^+^ BTLA^-^) and DC3s (CD36^-^ BTLA^+^) (**Figure 9B**). Thus, we hypothesized that this phenotypic key could facilitate the *in vivo* identification of moDCs and their discrimination from cDC2Bs/DC3s. In the lungs of MAZ mice treated with HDM, we first assessed whether there was an increase in DCs co-expressing *Ms4a3*-Ai14 and *Zbtb46*^GFP^, and how this compared with the dynamics of the non-cDC compartment identified as *Clec9a*-Ai14^-^ *Zbtb46*^GFP+^ cells in *Clec9a*^iCre^;Ai14;*Zbtb46*^GFP^ (CAZ) mice. Both *Ms4a3*-Ai14^+^ *Zbtb46*^GFP+^ DCs in MAZ mice and *Clec9a*-Ai14^-^ *Zbtb46*^GFP+^ DCs in CAZ mice increased in absolute numbers and relative frequency in the lungs upon HDM sensitization, thus suggesting that *in vivo* moDCs could develop in the lung in this pathophysiological context (**Figure 9C**). To further explore this scenario, we examined the two different populations for the expression of CD36 and BTLA. In cell suspensions from lungs of HDM-exposed MAZ and CAZ mice, CD36 and BTLA expression segregated two populations putatively corresponding to CD36^+^ BTLA^-^ moDCs versus CD36^-^ BTLA^+^ DC3s (**Figure 9D**) (**Supplementary Figures 4-5**). To further test this hypothesis, we profiled the gene expression programs of the various DC and Mo-derived populations present in the lungs and its draining mediastinal lymph nodes (MedLNs) from HDM-treated MAZ mice, by scRNA-seq using the FB5P-seq FLASH protocol. We sorted the following cell populations based on the indicated phenotypes (**Supplementary Figures 6-7**): (a) from the lungs; cDC1s (*Ms4a3-*Ai14^-^ *Zbtb46*^GFP+^ CD11c^+^ MHC-II^+^ Sirpα^-^ XCR1^+^), cDC2s (*Ms4a3-*Ai14^-^ *Zbtb46*^GFP+^ CD11c^+^ MHC-II^+^ XCR1^-^ Sirpα^+^ CD36^-^), DC3s (*Ms4a3-*Ai14^+^ *Zbtb46*^GFP+^ CD11c^+^ MHC-II^+^ XCR1^-^ Sirpα^+^ CD36^-^), CD11c^high^ moDCs (*Ms4a3-*Ai14^+^ *Zbtb46*^GFP+^ CD11c^high^ MHC-II^+^ XCR1^-^ Sirpα^+^ CD36^+^), CD11c^int^ moDCs (*Ms4a3-*Ai14^+^ *Zbtb46*^GFP+^ CD11c^int^ MHC-II^+^ XCR1^-^ Sirpα^+^ CD36^+^) and GMP-derived MFs (*Ms4a3-*Ai14^+^ *Zbtb46*^GFP-^ F4/80^+^ MHC-II^+^), and (b) from the MedLNs; cDC2s (CD45^+^ *Ms4a3-*Ai14^neg^ *Zbtb46*^GFP+^ CD11c^+^ MHC-II^+^ XCR1^-^ Sirpα^+^), DC3s (CD45^+^ *Ms4a3-*Ai14^+^ *Zbtb46*^GFP+^ CD11c^+^ MHC-II^+^ XCR1^-^ Sirpα^+^), *Ms4a3*-Ai14^+^ GFP^int^ moDCs (CD45^+^ *Ms4a3-*Ai14^+^ *Zbtb46*^GFPint^ CD11c^+^ MHC-II^+^) and *Ms4a3*-Ai14^+^ *Zbtb46*^GFP-^ MFs (CD45^+^ *Ms4a3-*Ai14^+^ *Zbtb46*^GFP-^ F4/80^+^ MHC-II^+^).

**Figure 9.**
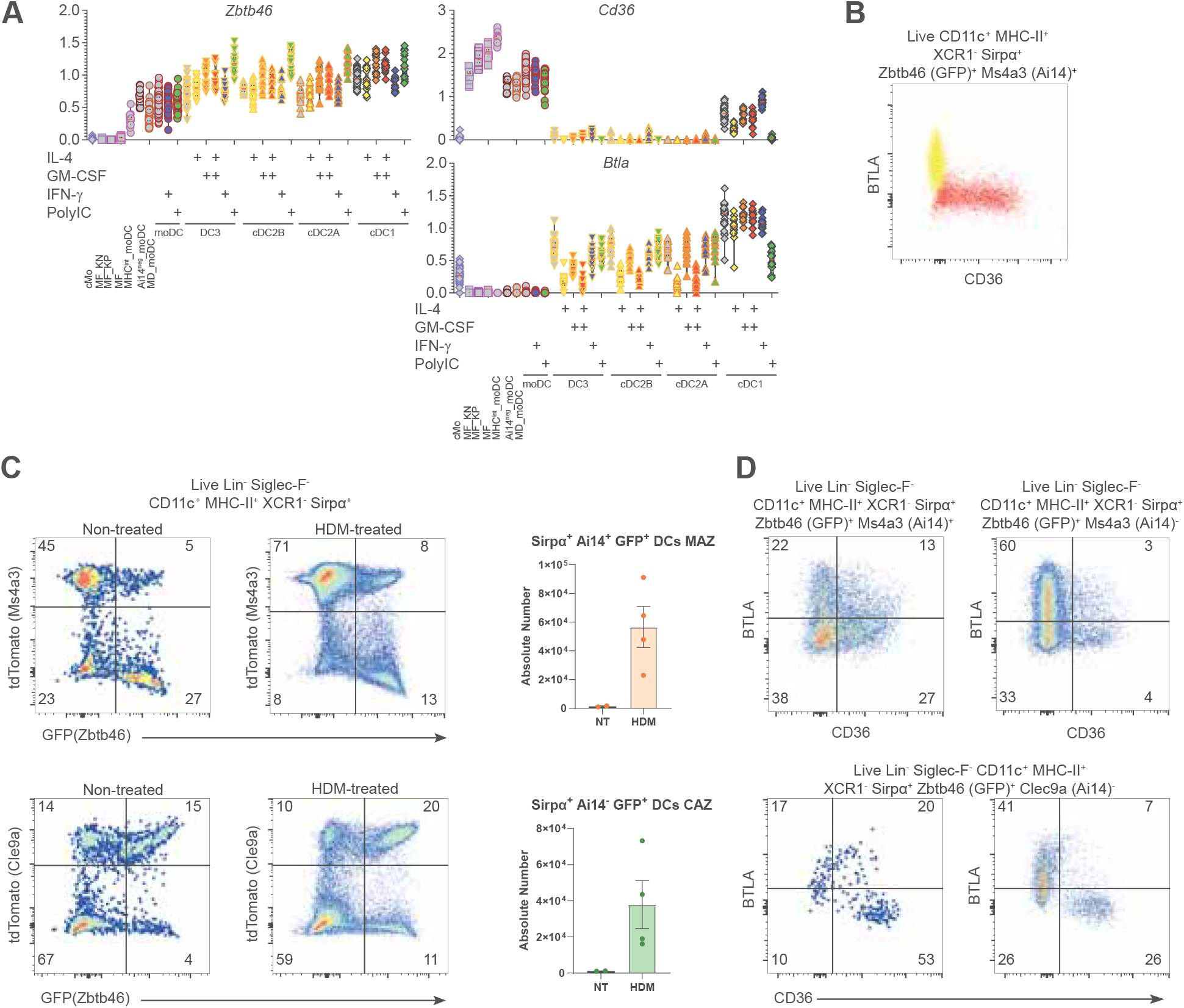
Sirpα^+^ CD36^+^ Ai14^+^ GFP^+^ arising in the lungs from HDM-treated mice could potentially correspond to *in vivo* moDCs. **A)** Violin plots showing the normalized expression of *Zbtb46*, *Cd36* and *Btla* from cells of the two mini-bulk datasets. **B)** Dot plot showing the cell surface marker expression of BTLA and CD36, comparing Ai14^+^ GFP^+^ DC3s arising in FLT3-L BM cultures to moDCs generated in GM-CSF and IL-4 BM cultures. **C)** Representative dot plots of individual mice and summary histograms showing the relative frequency and absolute numbers of *Ms4a3*;Ai14^+^ GFP^+^ and *Clec9a*;Ai14^-^ GFP^+^ cells arising in the lungs from mice exposed to HDM in comparison with their non-treated counterparts. **D)** Top: Dot plot showing cell surface expression of BTLA and CD36 within the Sirpα^+^ GFP^+^ DCs positive (left) or negative(right) for *Ms4a3*;Ai14 from mice treated with HDM, enriched upon magnetic depletion of CD3-, CD19-, Siglec--F and Ly6G-expressing cells. Bottom: Dot plots showing cell surface expression of BTLA and CD36 within Sirpα^+^ *Clec9a*;Ai14^-^ GFP^+^ DCs of untreated (left) versus HDM-instilled (right) mice.

The cellular composition of the peritoneum is heterogeneous. There is some unresolved overlap between cDC2s and serous cavity MFs, with the latter harboring varying degrees of DC features and F4/80 expression across different studies, with large cavity MFs (LCMs) expressing high levels of F4/80 and ICAM-2 but not CD226, conversely from small cavity MFs (SCMs) that also share some features with moDCs^94–99^. Peritoneal moDCs arising from Ly6C^+^ Mos have been reported to specifically upregulate CD226^91^ but whether they originate from GMP-derived or MDP-derived Mos and how they compare with SCMs remains to be established. Furthermore, it remains elusive whether DC3s arise in the peritoneum and whether these cells would resemble peritoneal moDCs, cDC2s or MFs. To clarify these issues, we harnessed MAZ and CAZ mice to compare the frequency and phenotype of the various MNP populations between the steady state versus inflamed peritoneum. To conduct a phenotypic assessment of moDC populations that could develop in peritonitis, we partially relied on adoptive transfer of Mos. For sorting the *Ms4a3*-Ai14^+^ Ly6C^+^ Mos, both for the experiments for the *in vitro* MNP atlas and for *in vivo* adoptive transfer experiments into the peritoneum of mice treated with thioglycolate, we strived to be as stringent as possible as minimal differences in Ly6C antibody signal alone can lead to contamination with DC precursors since their phenotype closely overlaps with that of cMos^10,20^ (**Supplementary Figure 8**). By excluding c-Kit and *Zbtb46*^GFP+^ cells, and gating on cells fate-mapped by *Ms4a3*, we managed to sort pure GMP-derived Mos, as confirmed in our mini-bulk experiments where no DC progeny was observed, contrary to conditions with less stringent methods for cMo purification.

To get a better picture of the cell composition, we excluded Ly6D-expressing cells, encompassing plasmacytoid DCs and B cells, and we split the remaining cells according to their expression of CD11c and MHC-II (**Figure 10A**) (**Supplementary Figure 9**). In this manner, we segregated the cells into CD11c^+^ MHC-II^+^ and CD11c^-^ fractions. Within Sirpα^+^ cells, we then evaluated the relative abundance of those fate-mapped by either *Clec9a* or *Ms4a3*, using CAZ and MAZ mice, and we assessed their expression of *Zbtb46*^GFP^. We observed that CD11c^+^ MHC-II^+^ Sirpa^+^ *Zbtb46*^GFP+^ DCs were mainly fate-mapped by *Clec9a* and not by *Ms4a3*. In contrast, *Clec9a* did not significantly label CD11c^-^ cells. Yet, there was a sizable population of CD11c^-^ Sirpa^+^ *Ms4a3*-Ai14^-^ *Zbtb46*^GFP+^ cells predominantly found at the steady state peritoneum that shrunk upon peritonitis induction. Upon thioglycolate administration, we observed an increase in *Ms4a3-*Ai14^+^ *Zbtb46*^GFP-^ and *Ms4a3-*Ai14^+^ *Zbtb46*^GFP+^ cells within the CD11c^-^ fraction, but only minor changes among CD11c^+^ MHC-II^+^ cells. To gain further insight into the identity of the *Ms4a3*-Ai14^+^ cells, we performed adoptive transfer of GMP-derived Mos, sorted from the BM of MAZ mice, into the peritoneum of congenic CD45.1^+^ WT C57BL/6 mice challenged with thioglycolate (**Figure 10B**) (**Supplementary Figures 9-10**). Immune cells were harvested by performing peritoneal wash 3 days later. Three different populations could be observed based on *Ms4a3*-Ai14 and *Zbtb46*^GFP^ expression, one comprising *Ms4a3*-Ai14^+^ *Zbtb46*^GFP+^ cells, and two fractions encompassing *Zbtb46*^GFP-^ cells with varying intensities of *Ms4a3*-Ai14 expression. We then carried a phenotypic analysis based on cell surface expression of canonical mO and MF markers. *Ms4a3*-Ai14^+^ cells could be split according to F4/40 expression. The F4/80^+^ fraction likely corresponded to peritoneal MFs, whereas the F4/80^-^ cells were negative for MHC-II and co-expressed Ly6C and Ly6G, suggesting that they were neutrophil-like Mos. When comparing *Ms4a3*-Ai14^+^ *Zbtb46*^GFP+^ cells and F4/80^+^ MFs, we did not observe major differences in terms of cells surface marker expression. Both fractions were positive for F4/80, showed high levels of MHC-II, but did not express CD11c or Ly6C, while showing variable expression of Ly6G. Peritoneal CD226^+^ MNPs have been reported to correspond to SCM or a canonical moDC population found at steady state and upon inflammation^91,100^. We therefore screened for CD226 expression within the progeny of the *Ms4a3-*Ai14^+^ Ly6C^+^ Mos adoptively transferred to thioglycolate-treated mice (**Figure 10C**) (**Supplementary Figure 10**). We were unable to find a sizeable portion of cells positive for CD226, suggesting that GMP-derived *Ms4a3-*Ai14^+^ Ly6C^+^ Mos do not give rise CD226^+^ peritoneal MNPs. Instead, we found that the progeny of *Ms4a3-*Ai14^+^ Ly6C^+^ Mos encompassed CD36^-^ and CD36-expressing cells. The MDP has been proposed to generate CD226^+^ Mfs in the peritoneum^9^. We therefore decided to sort *Ms4a3-*Ai14^-^ Ly6C^+^ Mos as *Zbtb46*-GFP^-^ Ly6G^-^ *Ms4a3;*Ai14^-^ Ly6C^+^ Ly6D^-^ B220^-^ c-Kit^-^ CD11b^+^ CD115^+^ cells, after performing Mo enrichment using the EasySep™ Mouse Monocyte Isolation Kit from StemCell technologies. Afterwards, we performed the adoptive transfer of these *Ms4a3-*Ai14^-^ Mos to congenic CD45.1^+^ WT C57BL/6 mice and we carried out a phenotypic characterization of the progeny (**Figure 10C**) (**Supplementary Figures 9-10**). The *Ms4a3-*Ai14^-^ Ly6C^+^ Mos developed into CD226^+^ cells while also generating a CD36^+^ fraction. These results thus support the hypothesis that MDP-derived but not GMP-derived Mos can generate CD226-expressing MNPs *in vivo* in the peritoneum and that these cells correspond to peritoneal moDCs or SCMs.

**Figure 10.**
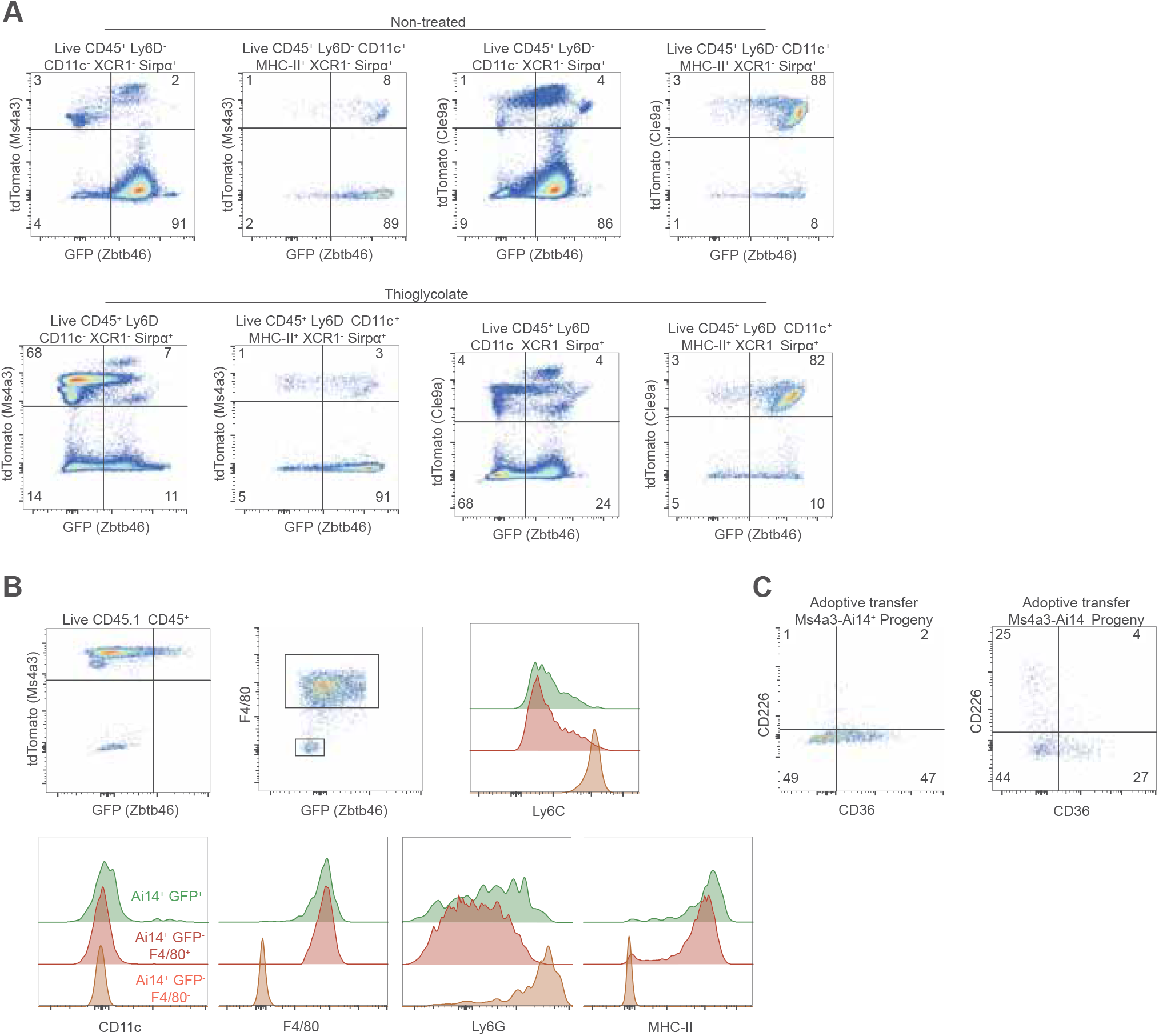
Ly6C^+^ *Ms4a3*;Ai14^+^ and Ly6C^+^ *Ms4a3*;Ai14^-^ monocytes give rise to convergent and unique monocyte-derived populations in the inflamed peritoneum. **A)** Relative abundance of fate mapping and reporter expression shown by *Ms4a3-Cre*;Ai14;*Zbtb46*GFP and *Clec9a-Cre*;Ai14;*Zbtb46*GFP mice within CD11c^-^ versus CD11c+ MHC-II^+^ cells at steady state or upon thioglycolate-induced peritonitis. **B)** Dot plots showing the three different populations that develop in the inflamed peritoneum upon Ai14^+^ Ly6C^+^ BM monocyte transfer to CD45.1 mice treated with thioglycolate. Histograms show the expression levels of different markers across the three different populations. **C)** Dot plots displaying the relative expression of CD226 versus CD36 in the progeny from adoptively transferred *Ms4a3*;Ai14^+^ Ly6C^+^ BM monocytes versus *Ms4a3*;Ai14^-^ Ly6C^+^ BM monocytes injected into CD45.1 mice. *Ms4a3*;Ai14^+^ and *Ms4a3*;Ai14^-^ Ly6C^+^ monocytes were adoptively transferred to different mice and never in competition. The cells were gated as Live CD45.1^-^ CD45.2^+^.

In the peritoneum, the different populations we have described have not been transcriptionally profiled at the single-cell level and their functions largely remain to be characterized. We thus proceeded to characterization of these populations by scRNA-seq using the FB5P-seq FLASH protocol, for the three different populations arising from GMP-derived *Ms4a3-*Ai14^+^ Ly6C^+^ Mos, *Ms4a3*-Ai14^+^ *Zbtb46*^GFP+^ cells, *Ms4a3*-Ai14^+^ *Zbtb46*^GFP-^ MFs and *Ms4a3*-Ai14^+^ *Zbtb46*^GFP-^ neutrophil-like Mos, as well as for the two populations, CD226^+^ versus CD226^-^, observed following the adoptive transfer of *Ms4a3-*Ai14^-^ Ly6C^+^ Mos (**Supplementary Figure 11**).

### The gene expression profiling of MNP types isolated *ex vivo* from the lung, MedLN and peritoneum of MAZ mice identifies two main moDC populations in the lung, with one only also detected in the MedLNs, and CD226^+^ MNPs in the peritoneum resembling either DCs or SCMs

To compare at the single cell level the distinct MNP populations sorted from the lung and MedLNs of HDM-treated mice, with those arising from the Mos adoptively transferred into the peritoneum of thioglycolate-treated mice, we integrated these two scRNA-seq datasets to generate a single UMAP leading to the identification of 27 cell clusters (C0 to C26) **(Figure 11A)**. For easing data illustration and interpretation, we chose a graphical key to represent the clusters **(Figure 11A)**, and their ordering on histograms **(Figure 11B)** or violin plots **(Figure 11C-E)**, according to their proximity as assessed by their relative location on the UMAP (**Figure 11A**) and the similarity in their composition in the different sorted cell populations (**Figure 11B**). On the UMAP, the cells remained largely grouped according to their anatomical compartment of origin, with only two clusters encompassing cells from both the lung and the peritoneum, C23 and C24 **(Figure 11B, top histograms)**. C23 comprised mainly peritoneal cells sorted as putative CD226^+^ moDCs, together with cells sorted from the lung parenchyma as moDCs or cDC2s (**Figure 11B, bottom histograms; Figure 11C**). C24 included cells sorted from the lung parenchyma mainly as putative CD11c^hi^ moDCs or from the peritoneum, sorted as GFP^hi^ moDCs or GFP^+^ MFs (**Figure 11B, bottom histograms; Figure 11C**). The cells from the MedLNs were always mixed in varying proportions with cells from the lung parenchyma. In contrast, a few clusters were almost exclusively composed of cells from the lung parenchyma, namely C1, C2 and C25 **(Figure 11B, top histograms)**, which were highly enriched in cells sorted as putative moDCs (**Figure 11B, bottom histograms; Figure 11C**), in addition to C13 that corresponded to the majority of the cDC1s, which had been exclusively sorted from the lung parenchyma.

**Figure 11.**
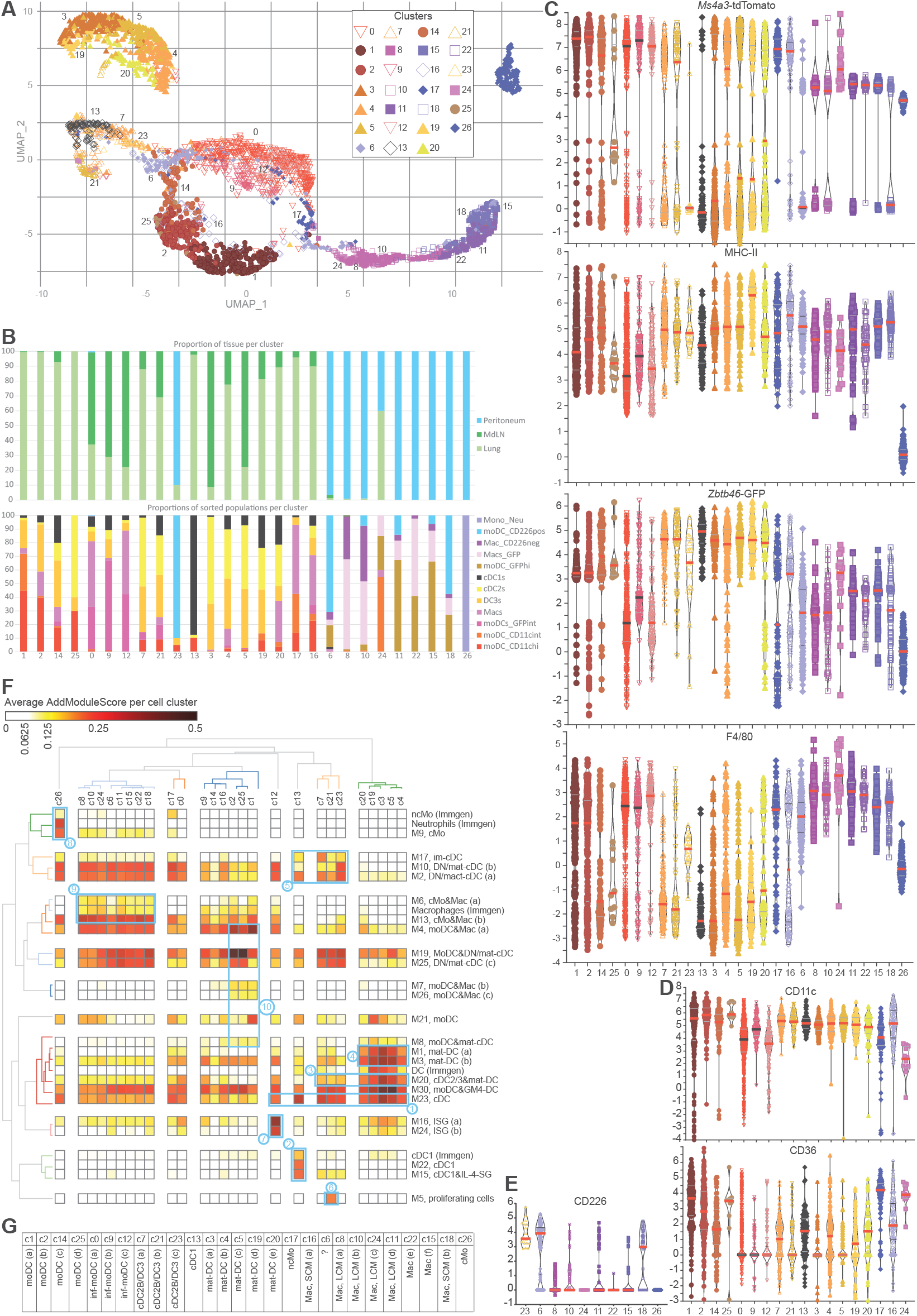
Lung MNPs from HDM-treated mice contain various moDC populations found in different activation states. **A)** UMAP representation depicting the integrated dataset comprising MNPs isolated from the lungs and MedLNs of MAZ mice treated with HDM and various monocyte-derived progenies from thioglycolate treated mice following the adoptive transfer of *Ms4a3*;Ai14+ and *Ms4a3*;Ai14-monocytes. **B)** Relative composition according to the tissue of origin of the cells comprising each cluster. **C)** Violin plots displaying the MFIs corresponding to lung, MedLN and peritoneal MNPs, corresponding to single cell surface marker instensity on the day of sorting. **D)** Violin plots displaying the MFIs corresponding to lung and MedLN MNPs, corresponding to the cell surface marker instensity on the day of sorting. **E)** Violin plot displaying the MFI of CD226 in the progeny of Ms4a3;Ai14-monocytes adoptively transferred to CD45.1 mice corresponding to single cell surface marker instensity on the day of sorting. **F)** Heatmap showing the gene module scores in each individual cluster utilizing specific geen signatures retrived from Immgen as well as the genes modules indentified by SciGeneX in the in vitro minibulk atlas. **G)** Proposed cellular identity for each cluster based on gene module annotation.

To help annotate the different clusters for cell types and activation states, we calculated enrichment scores of all individual cells for gene modules that had been generated previously and independently from other datasets, namely from our atlas of *in vitro*-generated MNPs as reported earlier in this study (nomenclature Mx), and from the reanalysis of the Immgen microarray database (Immgen). We then calculated the average score of each cluster for each gene module, and generated the corresponding heatmap with hierarchical clustering of clusters in columns and gene modules in rows **(Figure 11F)**. M23, corresponding to genes expressed to higher levels in cDCs in our *in vitro* atlas (**Figures 4-6**), was expressed to higher levels in clusters C13, C7, C21, C23, C3, C4, C5, C19, C20 and C12 **(Figure 11F, box n°1)**, suggesting that these clusters corresponded to cDCs. C13 was also highly enriched for cDC1 modules **(Figure 11F, box n°2)**, consistent with the fact that it mostly encompassed cells sorted as cDC1s **(Figure 11B)**. C7, C21, C23, C3, C4, C5, C19 and C20 corresponded to cDC2Bs/DC3s, since they were enriched for M20 that was selectively expressed in cDC2s/DC3s and mature DCs in our *in vitro* atlas **(Figure 11F, box n°3)**. C3, C4, C5, C19 and C20 corresponded more precisely to mature cDC2s/DC3s, since they were enriched for M1 and M3 **(box n°4)** whereas clusters C7, C21 and C23 corresponded to immature cDC2s/DC3s, since they were enriched in modules expressed specifically by immature DCs (M17) or down-regulated in mature DCs (M2, M10) **(box n°5)**. In addition, C21 corresponding to proliferating cells **(box n°6)**. Consistent with this annotation, the levels of *Zbtb46*^GFP^ expression were the highest in clusters with elevated scores for cDC modules (C7, C21, C23, C13, C3, C4, C5, C19 and C20), which also expressed relatively high levels of MHC-II and CD11c, but low levels of F4/80 and CD36 (except for cDC1s as expected) **(Figure 11C-D)**. M16 and M24, corresponding to ISGs, were expressed to the highest degree in cluster C12, and also to relatively high levels in C0 and C9, similar to those observed in the mature cDC clusters C3, C5 and C19, above the levels of all other clusters **(Figure 11F, box n°7)**. Clusters C0, C9 and C12 were constituted almost exclusively by cells fate-mapped by *Ms4a3*-Ai14 and expressing relatively high levels of F4/80 but low levels of *Zbtb46*^GFP^ **(Figure 11C)**, suggesting a monocytic origin. Yet, they did express neither CD36 **(Figure 11C)** nor high levels of the modules corresponding to Mos, MFs or moDCs **(Figure 11F)**. As we will see later upon mining of their specific gene expression profile, clusters C0, C9 and C12 appear to correspond to monocyte-derived inf-DCs. Of note, they encompassed cells from both the lung parenchyma and the MedLNs **(Figure 11C)**. M9, corresponding to genes specifically expressed in cMos in our *in vitro* atlas, was specifically enriched in C26, which also expressed selectively high levels of the Immgen gene modules for ncMos and neutrophils **(Figure 11F, box n°8)**. Hence, C26 corresponded to neutrophil-like Mos, accordingly to their sorting gate (**Figure 11B**). C17 corresponds to ncMos, since it harbors the highest enrichment for the corresponding Immgen module and this identity will be confirmed later upon deeper analysis of their specific gene expression profile. Clusters C8, C10, CC24, C6, C11, C15, C22 and C18 were enriched for the Immgen MF gene module as well as for the modules M6 and M3 corresponding to genes specifically expressed in MFs and cMos in our in vitro atlas **(Figure 11F, box n°9)**, but scored low for Mo modules (M9 and ncMo of Immgen). All these clusters expressed high levels of F4/80 but low levels of *Zbtb46*^GFP^ **(Figure 11C)**. Hence, they appear to correspond to MFs. All these clusters but C6 and C18 were almost exclusively made up by cells fate-mapped by *Ms4a3*-Ai14 (**Figure 11C**). C6 and C18 encompassed a majority of cells corresponding to the CD226^+^ progeny of the MDP-derived Mos adoptively transferred *in vivo* in the peritoneum **(Figure 11B and E)**. Thus, clusters C6 and C18 specifically correspond to peritoneal SCMs, whereas C8 and C10 correspond to LCMs, as will be confirmed later upon mining their specific gene expression profiles. Clusters C1, C2 and C25 are selectively enriched for the combination of our *in vitro* atlas modules expressed specifically by moDCs or shared between moDCs and MFs or mature DCs **(Figure 11F, box n°10)**. These clusters were enriched in cells sorted as moDCs **(Figure 11B)**, and were constituted almost exclusively by cells fate-mapped by *Ms4a3*-Ai14, specifically expressing CD36, harboring the highest levels of CD11c and intermediated levels of *Zbtb46*^GFP^ and F4/80 **(Figure 11C-D)**. Thus, C1, C2, C25 and by inference C14 represent putative moDCs. Interestingly, on the UMAP the C18 SCM cluster was close to the clusters of LCMs, whereas the C6 SCM cluster was closer to DCs. In summary, this cluster annotation identified two main moDC populations in the lung, with one only also detected in the MedLN, and CD226^+^ MNPs in the peritoneum resembling either DCs or SCMs **(Figure 11G)**.

### *Ex vivo*-isolated MNPs from the lungs, MedLNs and peritoneum of MAZ mice share the expression of core genes of cellular identity with the murine MNPs generated in the *in vitro* atlas and with their human counterparts

Following gene module analysis, we queried our merged dataset of lung, MedLNs and peritoneal MNPs to identify gene modules associated to the identity of cellular lineages, including transcriptomic fingerprints conserved between *in vitro*-derived murine moDCs, their putative *in vivo* equivalents and human *in vitro*-derived moDCs.

cDC1s could easily be identified due to their unique high expression of *Wdfy4*, *Itgae* (encoding CD103), *Clec9a*, *Xcr1* and *Naaa* (**Figure 12**, gene names highlighted in bold black font).

**Figure 12.**
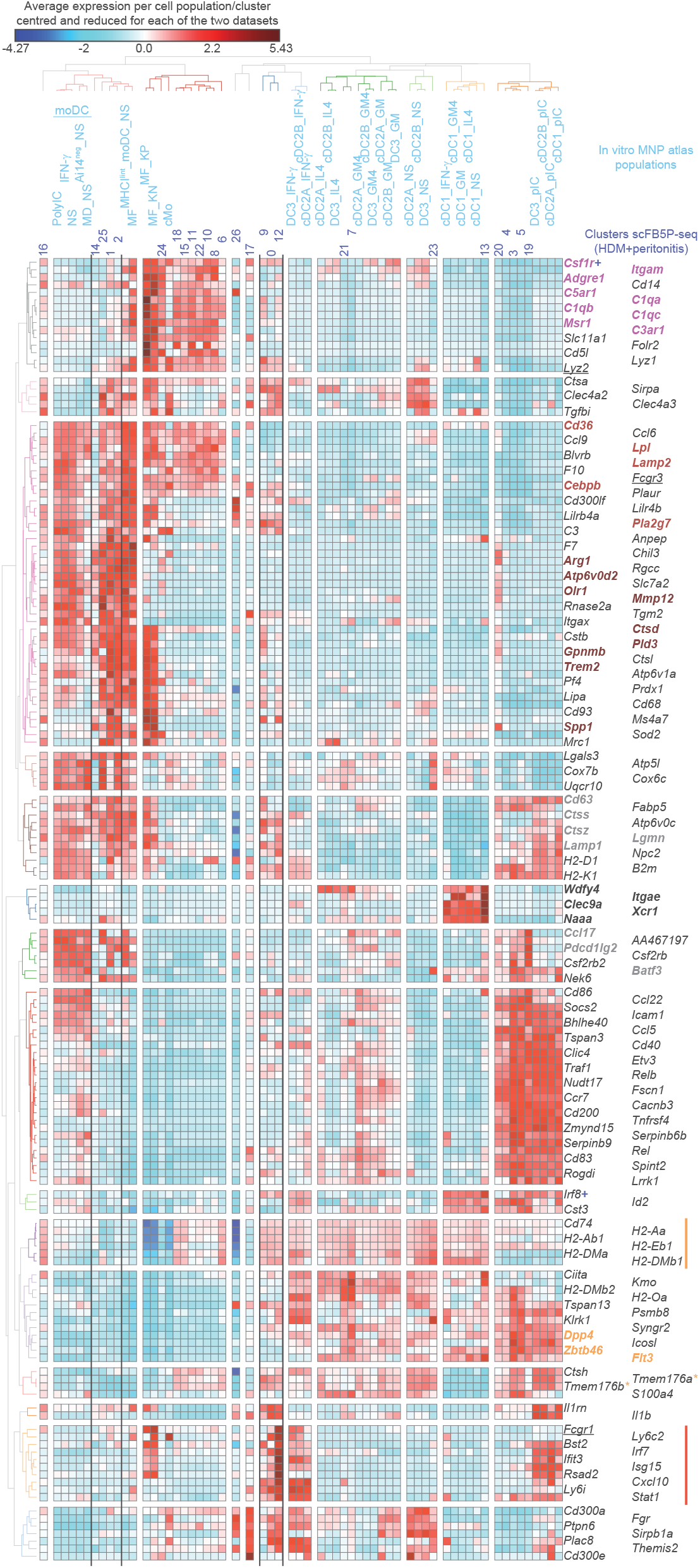
*Ex vivo*-isolated MNPs from the lungs, MedLNs and peritoneum of MAZ mice share the expression of core genes of cellular identity with the murine MNPs generated in the in vitro atlas and with their human counterparts. Heatmap showing the normalized average gene expression for each cluster of MNPs isolated from the lungs MedLNs and peritoneum of MAZ mice as compared to the various MNPs generated in our in vitro atlas. Selected genes are indicated with a specific color code: Genes enriched in macrophages (purple), genes usually employed to define cDC2Bs as compared to cDC2As (underlined gene names), genes associated to lipid handling and phacytosis shared between moDCs and macrophages (light brown) genes specifically expressed by moDCs (dark brown), genes specific to cDC1s (bold black), genes involved in antigen processing and presentation or in the regulation of T cell activation (grey), genes encoding MHC-II molecules (vertical orange bar), core cDC genes (orange), *Tmem176a* and *Tmem176b* (orange asterisks), interferon stimulated genes (red vertical line).

cDC2s shared with cDC1s a core cDC transcriptomic signature, including the genes *Flt3*, *Zbtb46*, *Dpp4* and *Syngr2* (**Figure 12**, bold orange names). Lung cDC2s have been reported to mainly comprise cDC2Bs/DC3s^8^. However, in the context of HDM sensitization, the genes known to be selectively expressed to higher levels in the cDC2B/DC3 lineage over the cDC2A one, including *Lyz2*, *Fcgr1* and *Fcgr3*^8,20,24,37^, were most prominently enriched in Mo-derived cells (**Figure 12**, underlined gene names).

All populations annotated as MFs via gene module analysis (C6, C8, C10, C11, C15, C18, C22 and C24) expressed high levels of canonical MF genes including *Adgre1* (encoding F4/80), *Itgam* (encoding CD11b), *Csf1r*, *C5ar1*, *C3ar1*, C1qa*, C1qb, C1qc, Msr1* (**Figure 12**, bold purple names). These genes were also selectively expressed to high levels in murine and human Mo-derived MFs, over the levels observed in murine and human *in vitro*-derived moDCs, and were not detected in cDCs **(Figures 11-12)**.

Lung putative moDCs, corresponding to clusters C1, C2 and to a lesser extent C14 and C25, selectively expressed genes that were also uniquely shared between murine and human *in vitro*-derived moDCs as compared to Mos, MFs and cDCs, including *Gpnmb*, *Arg1*, *Atp6v0d2*, *Olr1* and *Mmp12*, as well as genes associated to lipid metabolism including *Trem2*, *Spp1*, *Ctsd* or *Pld3* (**Figure 12**, bold brown names). In addition, *ex vivo*-isolated moDCs shared with MFs a high expression of other genes related to lipid metabolism or phagocytic activity, including *Lpl*, *Cd36*, *Lamp2, Cebpb* and *Pla2g7*, akin to what was observed with *in vitro*-derived MNPs (**Figures 2, 5 & 7**). These observations suggest that common lipid handling programs are selectively active in moDCs in a manner that is conserved across species.

The Mo-derived cells present in clusters C0, C9 and C12 expressed only relatively low levels of both the core DC genes and the moDC genes. Yet, they shared with cDC2s relatively high levels of the genes encoding MHC-II molecules (*Cd74*, *H2-Aa*, *H2-Ab1*, *H2-Eb1*, *H2-DMa* and *H2-DMb1)* **(Figure 12)**, vertical orange bar) and of *Tmem176a* and *Tmem176b* (**Figure 12**, orange asterisks). Consistent with the gene module analysis **(Figure 11)**, they were especially characterized by a high expression of interferon-stimulated genes, including *Fcgr1*, *Ly6c2*, *Bst2*, *Irf7*, *Ifit3*, *Isg15*, *Rsad2*, *Cxcl10*, *Ly6i* and *Stat1* (**Figure 12**, vertical red bar), combined with high expression of *Irf8 and Csf1r* (**Figure 12**, red crosses), likewise to inflammatory cDC2s^21^. We will therefore refer to these cells as inflammatory moDCs, due to their sharing of characteristics with both moDCs and inf-cDC2s.

Both *in vitro*-derived and *ex vivo*-isolated moDCs did not express core cDC genes, including *Flt3*, *Zbtb46* and *Dpp4* (**Figure 12**, bold orange names). Both *in vitro*-derived and *ex vivo*-isolated moDCs shared with mature cDCs a high expression of genes encoding MHC-I molecules, *H2-D1*, *H2-K1*, *B2m*, or involved in antigen processing and presentation or the regulation of T cell activation, including *Lamp1*, *Ctss*, *Ctsz*, *Lgmn*, *Cd63*, *Pdcd1lg2*, *Ccl17* and *Batf3* (**Figure 12**, bold grey names), combined with lower levels of the genes encoding MHC-II than in immature cDCs (**Figure 12**, vertical orange bar). However, contrary to *in vitro*-derived moDCs, *ex vivo*-isolated moDCs did not express many of the other genes associated to cDC maturation, including *Ccr7*, *Fscn1*, *Tnfsf4*, *Icosl*, *Relb* and *Etv3*. In combination with their specific expression of *Arg1* and *Tgfbi*, this suggests that, in HDM allergy, moDCs are in a maturation state differing from that of cDC2s or of highly activated *in vitro*-derived moDCs, and might exert immunosuppressive or tolerogenic functions.

From the analysis of the various populations, *Mrc1* and *C5ar1* seemed to be enriched in Mo-derived cells, whereas *Dpp4* was expressed on cDCs and *Ms4a3*-Ai14^+^ *Zbtb46*^GFP-^ cells. When sorted, *Ms4a3*-Ai14^+^ *Zbtb46*^GFP+^ moDCs displayed higher levels of *Zbtb46*^GFP^ in comparison to *Ms4a3*-Ai14^+^ *Zbtb46*^GFP-^ cells, but they did not show substantial expression of *Zbtb46* or *Dpp4* at the single-cell level, with the exception of cluster 9. This may be due to the low copy number of these genes, or the higher background noise created by MF metabolic genes. It remains possible that there may be some discrepancies between mRNA and protein levels. Thus, to verify what we observed in scFB5P-seq data, we decided to assess, on the lung MNPs isolated from HDM-treated MAZ mice, the expression of CD88, CD206 and CD26, encoded respectively by *C5ar1*, *Mrc1* and *Dpp4*, in order to test whether the combination of these cell surface markers would allow to separate moDCs from MFs and cDC2Bs/DC3s. Firstly, we observed that CD26 could be used as a surrogate for *Zbtb46*^GFP^ expression as the levels of both proteins were tightly correlated (**Figure 13A**) (**Supplementary Figure 12**). Then, we examined CD88 expression within CD26^+^ cells and found that the majority of CD88^+^ cells were fate-mapped by *Ms4a3*. Only a minority of *Ms4a3*-Ai14^+^ cells did not express CD88, putatively corresponding to DC3s. Furthermore, the majority of CD88^+^ CD26^+^ cells were fate mapped by *Ms4a3* and expressed high levels of *Zbtb46*^GFP^ (**Figure 13B-C**). Concurrently, we found that CD206, a marker traditionally expressed by interstitial MFs, was expressed on GFP-expressing cells, both fate-mapped and non-fate-mapped by *Ms4a3* (**Figure 13C**). Considering our single-cell analysis, we reasoned that CD206-expressing cells would be comprised of putative moDCs, potentially resembling the ones of clusters 2 and 3, and DCs present in cluster 4 that are enriched for *Cd209a*, *Clec10a*, *Cd206* and *Mgl2* expression. Therefore, the combined use of CD26, CD206 and CD88 could be exploited to identify three main DC populations (**Figure 13D-E**) (**Supplementary Figure 12**): (i) CD26^+^ CD206^-^ CD88^-^ Sirpα^+^ cDCs resembling those found in cluster 1 and enriched in transcripts consisting of *Ccr7*, *Chil3*, *Fscn1*, *Relb*, *Irf8* and *Id2*, (ii) CD26^+^ CD206^+^ CD88^-^ Sirpα^+^ cDCs resembling those in cluster 4 and enriched for the expression of *Cd209a*, *Clec10a*, *Cd206* and *Mgl2*, and (iii) CD26^+^ CD206^+^ CD88^+^ moDCs selectively expressing CD36 and likely encompassing cells in different maturation states corresponding to the gene expression programs of clusters 2 and 9. It seems that cDC2s from cluster 4 show a more activated or migratory phenotype, whereas those from cluster 1 seem to be more immature.

**Figure 13.**
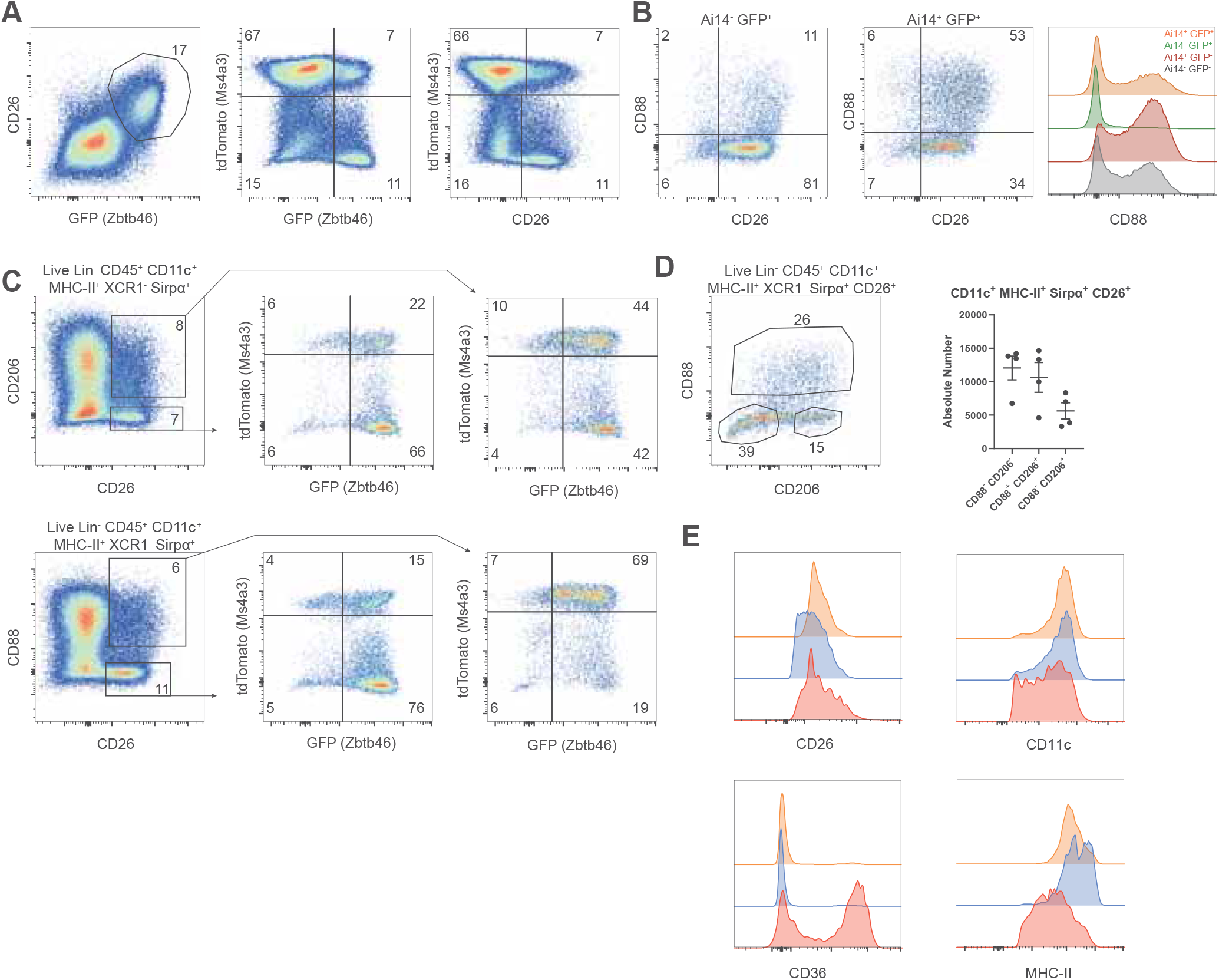
The combined use of CD26, CD88 and CD206 enables to identify moDCs in the lungs of C57BL/6J mice. **A)** Dot plots comparing the expression pattern of CD26 versus *Zbtb46-*GFP in *Ms4a3*-*Cre*;Ai14-*Zbtb46*GFP mice treated with HDM. The cells were gated as Live Lin^-^ CD45^+^ CD11c^+^ MHC-II^+^ XCR1^-^ Sirpα^+^. **B)** Dots plots comparing the distribution of CD88 and CD26 within Ai14^+^ GFP^+^ or Ai14^-^ GFP^+^ cells. Histograms assessing the levels of CD88 expression across the four different populations defined by Ms4a3-Cre fate-mapping and Zbtb46-GFP reporter expression. In all plots and histograms, the cells were gated as Live Lin^-^ CD45^+^ CD11c^+^ MHC-II^+^ XCR1^-^ Sirpα^+^. **C)** Assessment of relative abundance of Ai14^+^ GFP^+^ and Ai14^-^ GFP^+^ cells within CD26^+^ cells in combination or not with CD88 or CD206 expression. The cells were gated as Live Lin^-^ CD45^+^ CD11c^+^ MHC-II^+^ XCR1^-^ Sirpα^+^. **D)** Dot plot showing the Identification of three different populations within CD26-positive Sirpα^+^ DCs segregated by the expression of CD88 and CD206. Graph showing the absolute numbers corresponding to each of the three populations. **E)** Histograms showing the expression of CD26, CD11c, CD36 and MHC-II in CD88^-^ CD206^-^ (orange), CD88^-^ CD206^+^ (blue) and CD88^+^ CD206^+^ (red) cells. The cells were gated as Live Lin^-^ CD45^+^ CD11c^+^ MHC-II^+^ XCR1^-^ Sirpα^+^.

Overall, we observed that cDC2s and DC3s co-express the same surface markers and genes regardless of their origin and can be found co-clustering in different activation states, further supporting that they may correspond to the same DC type. With regard to moDCs, we confirm that they are a population of cells that arises *in vivo* upon HDM treatment and co-express canonical DC markers together with gene modules shared with either DCs or Mo/MFs.

We thus propose that moDCs in the allergic lung can be discriminated from cDC2s and DC3s on the basis of CD88 expression. In addition, we have a identified a fraction of cDC2s/DC3s that expresses CD206.

## Discussion

Here we showed that, likewise to human *in vitro*-derived moDCs, the majority of the murine DCs developing from total BM cultured in GM-CSF and IL-4 derive from Mos, do not express Flt3 or only at very low levels, are fate-mapped by *Ms4a3*^Cre^ but not *Clec9a*^Cre^, and do not express many of the canonical genes typical of the cDC lineages. Moreover, these cells align transcriptionally to human *in vitro*-derived moDCs, with a conserved selective expression of several genes including *Arg1*, *Slamf1*, *Sdc1* and *Dcstamp*. Hence, these cells do not belong to the cDC or DC3 lineages. We demonstrated that a significant fraction of the murine cells differentiated in vitro from total BM in the presence of GM-CSF and IL-4 are moDCs homologous to human moDCs.

In a previous publication, within BM cells with a Mo phenotype, a fraction was identified as Flt3^+^ CD11c^-^ cells (“R2 gate”), expressing *Cd209a* and giving rise to moDCs *in vivo*^101^. However, the nature of these cells was later reinterpreted as corresponding to progenitors of DC3s (pro-DC3s) differentiated from Ly6C^+^ MDPs without any Mo intermediate stage^20^. Our results may help understand this discrepancy. The R2 gate likely corresponds to a mixed population encompassing both pro-DC3s and Ly6C^+^ cMos, as previously suggested^20^, and this heterogeneity may have led to the opposite interpretations proposed for the nature of this cell population^20,101^. Here, we showed that *Cd209a* is not expressed in moDCs, neither *in vitro* nor *ex vivo*, and that cDC2Bs/DC3s require FLT3-L to develop but that their exposure to GM-CSF can induce their upregulation of *Cd209a*. This supports the interpretation that the splenic CD209a^+^ cells that have been reported to arise in mice bearing orthotopic B16-GM-CSF tumors^101^ may correspond to cDC2Bs/DC3s that have been exposed to GM-CSF. In contrast, the cells arising *in vitro*, from the culture of total BM explants or following the isolation of R2 cells, likely correspond to a mixture of CD209a^-^ moDCs and CD209a^+^ cDC2Bs/DC3s. Indeed, CD209a^+^ cells were shown to be a minor population in total BM cultures, while they constituted a sizeable fraction of the cells derived from isolated R2 cells^101^. This might be explained by the fact that R2 cells were cultured for only 2 days and in the presence MS-5 feeder cells, which could have provided an environment supportive of cDC2B/DC3 development. In contrast, total BM explants were cultured for 5 days; under conditions where most cDC2Bs and DC3s likely failed to differentiate or survive, due to the lack of supportive signals other than GM-CSF, resulting in ∼4% CD209a^+^ cells^101^. Moreover, in our own experiments, we used recombinant GM-CSF and IL-4, which may have been less permissive for cDC2B/DC3 survival than using GM-CSF-B16 supernatant that may contain supportive factors for this cell lineage.

We showed that cDC2As can be generated *in vitro* in FLT3-L cultures and we provide a series of genes they express selectively. To the best of our knowledge, we reported for the first time that murine DC3s can be generated *in vitro*, and that they develop in BM cultures supplemented with FLT3-L but not in GM-CSF and IL-4 cultures. Since cDC2Bs and DC3s showed very similar gene expression patterns irrespective of their stimulation conditions, we propose that they correspond to the same DC type differentiating though two different but converging ontogenetic paths. The notion that DC3s were FLT3-L-independent was partly cemented on experiments utilizing *Flt3l*^-/-^ mice^101^. However, their cytokine dependency was never evaluated *in vitro* in cultures supplemented with FLT3-L. We therefore propose that DC3s depend on FLT3-L for their development, but that they require GM-CSF to adopt an inflammatory phenotype that partially overlaps with the one of moDCs. DC3s share these properties with cDC2Bs. Given these results, the studies with human PBMCs reporting the conversion of CD1c^+^ CD14^-^ cDC2s into CD1c^+^ CD14^+^ DC3s^59,102^ will need to be re-evaluated. Based on their functional activity, surface marker expression and cytokine dependency, these cells might correspond to moDCs or MFs. Alternatively, the human CD1c^+^ CD14^+^ DC3s identified *in vivo* might constitute a unique lineage not present in mice. Future studies will be needed to compare *in vitro*-derived human and mouse moDCs with the human CD1c^+^ CD14^+^ cells derived *ex vivo* or in tumors from CD1c^+^ CD14^-^ cDC2s. Here we provide key transcriptomic modules to align human and mouse MNPs.

Studies with human immune cells further supported the requirement of GM-CSF for DC3 development, as human DC3s were present in B16 tumors implanted orthotopically in mice with a humanized immune system only when these tumors were engineered to express GM-CSF but not FLT3-L^27^. However, in this study, no side-by-side comparison was performed between the human DC3s observed in vivo in B16-GM-CSF tumors, and the CD14^+^ CD1c^+^ CD163^+^ cells arising from *in vitro* cultures of CD34^+^ hematopoietic progenitors cultured with MS5_FS12 cells and recombinant GM-CSF, or with human DCs differentiated in vitro from purified blood Mos in the presence of GM-CSF and IL-4 (e.g. classical human moDCs) ^27^. Our results thus raise the question whether *in vitro*-derived DC3s could have been confounded with moDCs, since we show that DC3s are FLT3-L-dependent and Mos can only survive and differentiate into moDCs in GM-CSF and IL-4 cultures but not in FLT3-L cultures.

To seek for *bona fide* moDCs *in vivo*, we have harnessed HDM-induced respiratory allergy as a model of type 2 inflammation inducing physiological co-production of the cytokines driving moDC differentiation *in vitro*, GM-CSF and IL-4/IL-13. To specifically trace the differentiation of Mos into moDCs *in vivo* during inflammation, we also harnessed another model that is easily amenable to Mo adoptive transfer: thioglycolate-induced peritonitis, which may rather resemble type 1 immunity.

For the model of HDM-induced respiratory allergy, previous publications have reported the development/recruitment of moDCs in the lungs^90^. However, at that time, no available tools enabled accurately discriminating moDCs from MFs and cDC2Bs/DC3s. Here, we show that CD11c^+^ MHC-II^+^ *Ms4a3*-Ai14^+^ *Zbtb46*^GFP+^ CD36^+^ candidate moDCs develop in the lungs upon HDM treatment and that they are different from CD11c^+^ MHC-II^+^ *Ms4a3*-Ai14^-/+^ *Zbtb46*^GFP+^ CD36^-^ cDC2Bs/DC3s, as well as from the CD11c^+^ MHC-II^+^ *Ms4a3*-Ai14^+^ *Zbtb46*^GFP-^ and CD11c^+^ MHC-II^+^ *Ms4a3*-Ai14^+^ *Zbtb46*^GFPint^ cells present in the lung parenchyma and MedLNs. Lung moDCs display a core transcriptional signature shared with *in vitro*-derived moDCs, encompassing the genes *Arg1*, *Atp6v0d2*, *Olr1*, *Mmp12* and *Gpnmb*, and express genes related to APC function including *H2-Aa, H2-Ab1, Batf3, Cd74, Cd83, B2m, Tapbp, Itgax, Lamp1, Ctss and Psmb8*, suggesting that they have the capacity to process antigens and present them to T cells. When compared to *Ms4a3*-Ai14^+^ *Zbtb46*^GFPlo^ cells, lung moDCs displayed a tissue repair tolerogenic profile resembling *Trem2* MFs^103,104^, including the expression of a specific gene module encompassing *Trem2*, *Chil3*, *Cstb*, *Lpl*, *Spp1* and *Apoe*, whereas *Ms4a3*-Ai14^+^ *Zbtb46*^GFP-^ cells from the lungs and *Ms4a3*-Ai14^+^ *Zbtb46*^GFP-^ and *Ms4a3*-Ai14^+^ *Zbtb46*^GFPint^ cells from the MedLNs harbored an interferon response transcriptomic signature encompassing inflammatory genes such as *Cxcl10*, *Cxcl9*, *Cd80*, *Cd86*, *Isg15*, *Stat1, Irf7*, *Il1b* and *Il1rn*.

Overall, *in vivo* in the lungs from HDM-treated mice, we have identified two distinct populations of Mo-derived cells with DC features. The first one, CD36^+^ *Ms4a3*-Ai14^+^ *Zbtb46*^GFP+^ cells, closely resembles *in vitro*-derived moDCs, although with a distinct maturation profile, strongly suggesting that they are *in vivo* moDCs. The other one, the *Ms4a3*-Ai14^+^ *Zbtb46*^GFP-/lo^ Mo-derived cells, does not show such a strong similarity to *in vitro*-derived moDCs, but is enriched for an inflammatory gene module, in particular ISGs, suggesting that they have undergone a different maturation program than the CD36^+^ moDCs. We annotated these cells as inf-moDCs as they bear striking similarities with the inf-cDC2s previously described^21^. Based on their phenotyping and on their gene expression profiling, both moDCs and inflammatory moDCs appear to have the potential to interact with T cells, considering their expression of high levels of MHC-II proteins and of genes encoding both costimulatory and co-inhibitory ligands. However, only inf-moDCs expressed *Ccr7* and were detected in the lymph nodes, suggesting that these cells may have migrated from the lungs and might contribute there to the induction of adaptive immunity, whereas CD36^+^ moDCs may rather play a local role in inflamed non-lymphoid tissues as previously proposed^16^.

One of the most striking differences observed between *in vitro*-derived moDCs differentiated in the presence of GM-CSF and IL-4 and putative moDCs that develop upon HDM sensitization, is their differential maturation state, with much lower expression if any of many maturation genes in the moDCs isolated *ex vivo* from the lung parenchyma. Exposure to M-CSF in the BM during Mo development may explain these differences. cMos may differentiate normally, egress the BM, circulate through the bloodstream and be eventually recruited to the lungs. It will only be at this stage that they would be exposed to the GM-CSF and IL-4/IL-13 produced locally. In these settings, only certain pathways may remain responsive to GM-CSF and IL-4, thus explaining the partial enrichment for some MF programs, as opposed to what could happen during *in vitro* BM cultures where progenitors would differentiate in Mos in the absence of M-CSF or at lower concentrations than what happens physiologically. Other cues different from M-CSF exposure may also favor the *in vivo* differentiation of Mos into MFs rather than moDCs, depending on the pathophysiological context. Much of the knowledge about MF differentiation comes from the study of their tissue-specific imprinting steered by locally produced mediators. However, it is possible that MF versus moDC specification already begins within the BM marrow during late Mo developmental stages and progresses further in the bloodstream. Future studies should explore this possibility. Another notable difference is that *in vitro*-derived cells are always subjected to some degree of shear stress through pipetting, high confluency conditions in the culture wells and the presence of some levels of endotoxins. These conditions could promote terminal maturation of the *in vitro*-derived moDCs. Alternatively, it remains to be evaluated in mice whether murine *in vivo* moDCs would be able to undergo further maturation upon the co-occurrence of high levels of GM-CSF, IL-4/13 and IFN-I, and whether these cells may align with murine *in vitro* moDCs polarized towards a type-1 functional polarization and human moDCs arising in lupus or tumor ascites^23,28,105,106^.

Not much is known about how microenvironments associated to type 1 immunity *in vivo*, particularly sterile inflammation, can ultimately shape Mo differentiation and activation towards states resembling *in vitro*-derived terminally mature moDCs. Murine moDCs have been reported to arise in thioglycolate-induced peritonitis, as CD226^+^ ICAM-2^-^ cells^91^. However, the peritoneal cavity is also known to house CD226^+^ SCMs that harbor mixed characteristics of MFs and DCs^94–100^, together with ICAM-2^+^ LCMs, and these populations have not been analyzed side-by-side at the single-cell level with peritoneal candidate moDCs and with the moDCs derived *in vitro*. The questions remained open whether a fraction of peritoneal CD226^+^ SCMs arising from MDP-derived Mos^9^ could correspond to peritoneal moDCs^91^. Here, we observed that *Ms4a3-*Ai14^+^ Ly6C^+^ GMP-derived Mos and *Ms4a3-*Ai14^-^ Ly6C^+^ MDP-derived Mos gave rise to an overlapping fraction of MFs with increased lipid metabolism, while also developing into two unique MF populations. GMP-derived Mos also selectively gave rise to a subset of LCMs enriched in genes related to tissue repair. MDP-derived Mos selectively gave rise to CD226^+^ MNPs enriched in the transcriptomic signature of SCMs and split into two clusters, close to LCMs versus to moDCs in the UMAP, and expressing distinct genes suggesting different functions, including genes encoding immunoregulatory molecules or ligands for lymphocyte co-receptors suggesting an enhanced capacity to interact with T cells.

We showed that GMP origin as assessed by *Ms4a3* fate-mapping was not sufficient to fully capture the identity of moDCs and DC3s. *In vitro*, we observed no major differences between *Ms4a3*-Ai14^+^ *Zbtb46*^GFP+^ versus *Ms4a3*-Ai14^-^ *Zbtb46*^GFP+^ moDCs in GM-CSF and IL-4 cultures, and between *Ms4a3*-Ai14^+^ *Zbtb46*^GFP+^ versus *Ms4a3*-Ai14^-^ *Zbtb46*^GFP+^ cDC2Bs/DC3s in FLT3-L cultures. *In vivo*, we report that *Ms4a3*-Ai14^+^ *Zbtb46*^GFP+^ cells are heterogeneous upon HDM sensitization, and encompass both moDCs and cDC2Bs/DC3s. Our deep side-by-side phenotypic and transcriptomic characterization of mouse CD11b^+^ MNPs is paving the way for the development of adequate fate-mapping models and phenotypic keys to accurately identify and target moDCs *in vivo* in future studies in order to determine their functions in comparison to those of cDC2Bs/DC3s and MFs.

## Materials and methods

### Mice

C57BL/6 mice were purchased from Janvier, France. All other mouse strains were bred at the Centre d’ImmunoPhénomique (CIPHE) or the Centre d’Immunologie de Marseille-Luminy (CIML), Marseille, France, under specific pathogen free-conditions and in accordance with animal care and use regulations. Mice were housed under a 12 h dark:12 h light cycle, with a temperature range of 20–22 °C and a humidity range of 40–70%. The animal care and use protocols were designed in accordance with national and international laws for laboratory animal welfare and experimentation (EEC Council Directive 2010/63/EU, September 2010). They were approved by the Marseille Ethical Committee for Animal Experimentation (registered by the Comité National de Réflexion Ethique sur l’Expérimentation Animale under no. 14; authorization #11–09/09/2011 and APAFIS #51110-2024091711569098 v3).

### Preparation of single cell suspensions

Lung single cell suspensions were prepared by digesting the lung tissue with Collagenase IV (Wortington biochemicals) and DNAse I (Roche Diagnostics) in RPMI using the lung dissociation program 37_m_LDK_1 in the gentle GentleMacs from Miltenyi. When performing depletion of CD3, CD19, Ly6G and Siglec-F, lung single cell preparations were first incubated with Fc-block and later incubated with biotinylated antibodies. Afterwards, cells were enriched via negative selection by using anti-biotin microbeads (Miltenyi) and collecting the resulting flow-through utilizing LS columns (Miltenyi). Harvest of peritoneal immune cell populations was performed by washing the peritoneal cavity twice and using a syringe with a 26 Gx needle. Red blood cells (RBCs) lysis was performed by using RBC lysis buffer (Life Technologies) for the lung cell suspensions, BM cell preparations and peritoneal washes.

### BM cultures

For cDC differentiation, BM cells were isolated and cultured for 8 to 10 days at 2.5*10^6^ cells/ml in 6-well plates using RPMI supplemented with 10 % FCS, 1 % glutamine (Gibco), 100 U/mL penicillin streptomycin, and 50 μM β-mercaptoethanol, with a 1/10 dilution of B16-FLT3-L supernatant.

For moDC differentiation in GM-CSF and IL-4, BM cells were isolated and cultured at a final concentration of 2.5*10^6^ cells/ml in a 6 well plates using RPMI supplemented with 10 % FCS, 1 % glutamine (Gibco), 100 U/mL penicillin streptomycin, and 50 μM β-mercaptoethanol, with 20 ng/ml of GM-CSF and 5 ng/ml of IL-4. At day 2 half of the supernatant was discarded and replaced by fresh medium with 40 ng/ml of GM-CSF and 10 ng/ml of IL-4. At day 4, 70 to 80 % of the supernatant was discarded and replaced by fresh medium with 20 ng/ml GM-CSF and 5 ng/ml of IL-4 up to 4 ml. The cells were harvested at days 5, 6 or 8. If going for 8 days culture, at day 6, half of the supernatant was discarded and replaced by fresh medium with 20 ng/ml GM-CSF and 5 ng/ml of IL-4.

moDC differentiation from ex vivo sorted BM Ai14^+^ Ly6C^+^ monocytes was performed in FACS tubes coated overnight with RPMI supplemented as mentioned above. Afterwards, 10^6^ Ai14^+^ Ly6C^+^ monocytes were plated in combination with 9*10^6^ CD45.1 BM cells in 6-well plates. They were cultured with RPMI as mentioned above, supplemented with 20 ng/ml of GM-CSF and 5 ng/ml of IL-4. At day 2 add 2 ml of fresh medium with 40 ng/ml GM-CSF and IL-4 at 10 ng/ml was added. The cells were harvested at day 4.

### Treatments

Asthma was induced by intranasal delivery on 40 μg of *D. pteronyssinus* extract FD (02.01.85.10 CITEQ Biologics) diluted in PBS, which was performed on day 1 of treatment and repeated on days 7 to 11. Lungs of HDM-treated mice were harvested at day 14.

Peritonitis induction was performed using 3.8 % thioglycolate diluted in sterile PBS. Each mouse was injected intraperitoneally with 1 ml of thioglycolate solution at day 1. At day 4, peritoneal cells were harvested. When performing adoptive transfer of monocytes, mice were treated with thioglycolate 18 hours prior to the adoptive transfer and immune cells were harvested 3 days after performing the adoptive transfer.

### DC stimulation

cDCs and moDCs were cultured *in vitro* as described above. cDCs were harvested at day 9 and moDCs at the 5. Afterwards, they were FACS sorted using sterile PBS as sheath fluid and plated in a 96-well plate with medium supplemented with FLT3-L for cDCs and medium with GM-CSF and IL-4 for moDCs, seeding 20000 or 15000 cells per well. Later, they were stimulated for 18 hours with either 5 μg/ml of Poly (I:C) (Invivogen), 1 μg/ml of LPS, 10 μg/ml of R848 or 5 ng/ml of IFN-γ (Peprotech). The readout was performed by collecting the supernatant and performing cytokine quantification using the LEGENDplex™ Mouse Inflammation Panel (13-plex) kit from BioLegend.

### Monocyte adoptive transfer to mice treated with thioglycolate

BM cells of MAZ mice were harvested. Later, Ms4a3Ai14^+^ Ly6C^+^ BM marrow monocytes were FACS sorted as Zbtb46GFP^-^ Ly6G^-^ Ms4a3Ai14^+^ Ly6C^hi^ c-Kit^-^. Afterwards, each mouse was injected intraperitoneally with Ms4a3Ai14^+^ Ly6C^+^ 10^6^ monocytes resuspended in 200 μl of PBS. For the adoptive transfer of Ms4a3Ai14^-^ Ly6C^+^ BM marrow monocytes BM cells of MAZ mice were harvested and monocytes were purified using the EasySep™ Mouse Monocyte Isolation Kit (StemCell) and FACS sorted as Zbtb46GFP^-^ Ly6G^-^ Ms4a3Ai14^-^ Ly6C^hi^ c-Kit^-^ Ly6D^-^ B220^-^ CD115^+^ CD11b^+^. Each mouse was intraperitoneally injected with 10^5^ Ms4a3Ai14^-^ Ly6C^+^ monocytes resuspended in 200 μl of PBS.

### Flow cytometry analysis

All stainings were done in PBS 1x supplemented with 2 mM EDTA (Sigma-Aldrich) and 10 % FCS except when staining for CLEC4A4 which was conducted in HBSS (with calcium and magnesium) supplemented with 10 % FCS. Before adding the antibody cocktail, cell suspensions were Fc-blocked with 2.4G2 for 15 minutes at 4 °C. All antibody stainings were performed for 30 min at 4 °C. Dead cell were discriminated in all experiments using LIVE/DEAD™ Fixable Blue Dead Cell Stain Kit, for UV excitation. Acquisition was performed on an Aurora five lasers (Cytek Biosciences) using SpectroFlo v.3.0.1 software (Cytek Biosciences) or on a BD Symphony A5 44 using Diva. Cell sorting was conducted on a FACS ARIA III (BD Biosciences) or on a FACS ARIA SORP using BD Diva. All data was analyzed with Flowjo.

### Antibodies

| Antibody name | Dilution | Reference |
| --- | --- | --- |
| BD Pharmingen™ Biotin Rat Anti-Mouse CD19 | 1/200 | 553784 |
| Biotin anti-mouse CD3ε Antibody BioLegend | 1/200 | 100304 |
| Biotin anti-mouse CD170 (Siglec-F) Antibody BioLegend | 1/200 | 155512 |
| Biotin anti-mouse Ly-6G Antibody BioLegend | 1/200 | 127603 |
| BD Horizon™ BUV737 Hamster Anti-Mouse CD11c BioLegend | 1/300 | 612796 |
| Ly-6D Monoclonal Antibody (49-H4), eFluor™ 450, eBioscience™ | 1/200 | 48-5974-80 |
| Brilliant Violet 510™ anti-mouse Ly-6G Antibody BioLegend | 1/200 | 127633 |
| BD Horizon™ BV605 Mouse Anti-Mouse CD45.2 | 1/200 | 563051 |
| PerCP anti-mouse CD90.2 (Thy1.2) Antibody BioLegend | 1/200 | 105322 |
| Alexa Fluor® 647 anti-mouse/rat XCR1 Antibody BioLegend | 1/200 | 148214 |
| Alexa Fluor® 700 anti-mouse I-A/I-E Antibody BioLegend | 1/300 | 107622 |
| APC/Cyanine7 anti-mouse CD172a (SIRPα) Antibody BioLegend | 1/150 | 144017 |
| APC anti-mouse CD26 (DPP-4) Antibody BioLegend | 1/200 | 137807 |
| FITC anti-mouse CD206 (MMR) Antibody BioLegend | 1/200 | 141703 |
| PE/Dazzle™ 594 anti-mouse CD206 (MMR) Antibody BioLegend | 1/200 | 141731 |
| PE/Cyanine7 anti-mouse CD88 (C5aR) Antibody BioLegend | 1/200 | 135809 |
| DCIR2 Antibody, anti-mouse APC Myltenyi (Staining in HBSS with calcium and magnesium) | 1/50 | 130-102-177 |
| PE/Cyanine7 anti-mouse CD36 Antibody BioLegend | 1/200 | 102615 |
| Biotin anti-mouse CD36 Antibody BioLegend | 1/200 | 102604 |
| Alexa Fluor® 700 anti-mouse Ly-6G Antibody BioLegend | 1/200 | 127621 |
| BD Pharmingen™ APC-Cy™7 Rat Anti-Mouse Ly-6C | 1/400 | 560596 |
| BD Pharmingen™ PE-Cy™7 Rat anti-Mouse CD117 | 1/200 | 558163 |
| BD Pharmingen™ Alexa Fluor® 647 Rat Anti-Mouse CD45R | 1/200 | 557683 |
| BD Horizon™ BUV395 Rat Anti-CD11b | 1/200 | 563553 |
| Brilliant Violet 711™ anti-mouse CD115 (CSF-1R) Antibody BioLegend | 1/200 | 135515 |
| APC anti-mouse/rat XCR1 Antibody BioLegend | 1/200 | 148205 |
| Spark PLUS UV395™ anti-mouse/rat XCR1 Antibody BioLegend | 1/200 | 148239 |
| APC anti-mouse CD226 (DNAM-1) Antibody BioLegend | 1/200 | 128809 |
| F4/80 Monoclonal Antibody (BM8), eFluor™ 450, eBioscience™ | 1/200 | 48-4801-82 |
| BD Horizon™ BV421 Rat anti-Mouse CD371 (Clec12A) | 1/200 | 564795 |
| APC anti-mouse CD45.1 Antibody BioLegend | 1/200 | 110713 |
| BD Horizon™ BUV395 Mouse Anti-Mouse CD45.2 | 1/200 | 564616 |
| BD Horizon™ BUV737 Mouse Anti-Mouse CD45.2 | 1/200 | 612779 |
| Brilliant Violet 711™ anti-mouse CD11c Antibody BioLegend | 1/200 | 117349 |
| F4/80 Monoclonal Antibody (BM8), PE-Cyanine7, eBioscience™ | 1/200 | 25-4801-82 |
| Brilliant Violet 711™ anti-mouse CD45.1 Antibody BioLegend | 1/200 | 110739 |
| Brilliant Violet 785™ Streptavidin BioLegend | 1/200 | 405249 |

### Mini-bulk RNA-seq and scRNA-seq

For the generation of the mini-bulk- and scRNA-seq data, we followed the FB5P-seq method previously published^82^. For mini-bulk RNA-seq 100 cells were sorted per well into ice-cold 96-well PCR plates (Thermo Fisher Scientific) containing 4 µl of lysis mix per well. For scRNA-seq, single-cells were FACS sorted into ice-cold 96-well PCR plates (Thermo Fisher Scientific) containing 2 µl of lysis mix per well.

Immediately after cell sorting, each plate was covered with an adhesive film (Thermo Fisher Scientific), briefly spun down in a benchtop plate centrifuge and frozen on dry ice. The reverse transcription (RT) reaction was performed with SuperScript IV (Thermo Fisher Scientific) in the presence of RNase inhibitor (Takara Bio), triton X 10 % (Thermo Fisher Scientific), betaine (Sigma-Aldrich), MgCl_2_ (Sigma-Aldrich) and well-specific template-switching oligonucleotides. For complementary DNA amplification, KAPA HiFi HotStart ReadyMix (Roche Diagnostics) was used with adapted primers and the program was run performing 16 PCR reaction cycles for mini-bulk RNAseq and 22 PCR reaction cycles for scRNAseq following one hour of reverse transcription. For library preparation, amplified cDNA from each well of a 96-well plate was pooled, purified with one round for mini-bulk RNAseq and two rounds for scRNAseq of 0.6x solid-phase reversible immobilization beads (AmpureXP, Beckman or CleanNGS, Proteigene) and finally eluted in nuclease-free water. After tagmentation and neutralization, tagmented cDNA was amplified with Nextera PCR Mastermix containing Nextera i5 primer (Illumina) and customized i7 primer mix. Libraries generated from multiple 96-well plates of single-cells and carrying distinct i7 barcodes were pooled for sequencing on an Illumina NextSeq2000 platform, with 100 cycles of P1, P2 or P3 flow cells, targeting 5*10^5^ reads per cell in paired-end, single-index mode with the following cycles: Read1 (Read1_SP, 67 cycles), Read i7 (i7_SP, 8 cycles) and Read2 (Read2_SP, 16 cycles). For mini-bulk RNAseq samples sorted in different days and from different cultures and from different sequencing runs were pooled together in the same Seurat object. For scRNAseq in lungs and MedLNs of HDM treated MAZ mice, two pools of 5 mice were generated sorting at least one plate per population from each pool. Cells coming from the lungs and MedLNs were sorted in two separate days and belong to different mice treated at different times, with the same HDM regimen. Plates loaded with cells from the two organs were sequenced in the same run. For scRNAseq in peritonitis, cells developing from adoptively transferred *Ms4a3*Ai14^+^ Ly6C^+^ monocytes were pooled from 4 C57BL6 mice. On a separate day the cells developing from adoptively transferred Ms4a3Ai14^-^ Ly6C^+^ monocytes were pooled from two CD45.1 mice. Plates loaded with cells from the two adoptive transfer systems were sequenced in the same run.

### BubbleGUM analysis

The BubbleGUM software^84^ was used to generate DC type-specific transcriptomic signatures from the *in vitro* atlas of DCs and macrophages to assess their enrichment across cell types from other datasets. BubbleGUM is an open-source software composed of two modules; i) GeneSign, which generates statistically significant gene signatures and ii) BubbleMap, which automatically assesses the enrichment of input gene signatures between all possible pairs of conditions from independent datasets, based on gene set enrichment analysis (GSEA) methodology^107^, and which generates an integrated graphical display.

### Statistical analysis and reproducibility

No statistical tests were conducted for murine experiments as our readouts were mainly descriptive. For mini-bulk-RNAseq at least 16 wells divided in two replicate plates (8 per plate) were loaded per cell type and condition. All scRNAseq experiments were performed by loading at least two plates per cell type. For the experiments measuring cytokine secretion by the various DC types the supernatant from each cell type and condition was measured by making at least three replicate wells. For the statistical analysis a one-way ANOVA was performed, ensuring homoscedastic variance and normality of residues.

## Acknowledgements

We thank CIPHE as well as the staff of the CIML mouse houses for their assistance in the breeding of mice. We thank F. Ginhoux (Institut Gustave Roussy, France), for providing *Ms4a3*^Cre^ mice. This research was funded by the French National Research Agency (ANR; no. ANR-22-CE14-0023-02, ‘nRegSKIN’, and ANR-23-CE15-0001, ‘RIPIREVI’, to M.D.). We also acknowledge institutional support from CNRS (French National Centre for Scientific Research), INSERM (French National Institute of Health and Medical Research) and Aix-Marseille Université. We thank Stoyan Ivanov and Alexandre Gallerand (Université Côte d’Azur, CNRS, LP2M, Nice, France) for scientific discussions and feedback on the manuscript. We also thank Bart Lambrecht and Lukas Amon (VIB Center for Inflammation Research, Ghent, Belgium) for scientific discussions.

## Supplementary material

**Supplementary Figure 1.**
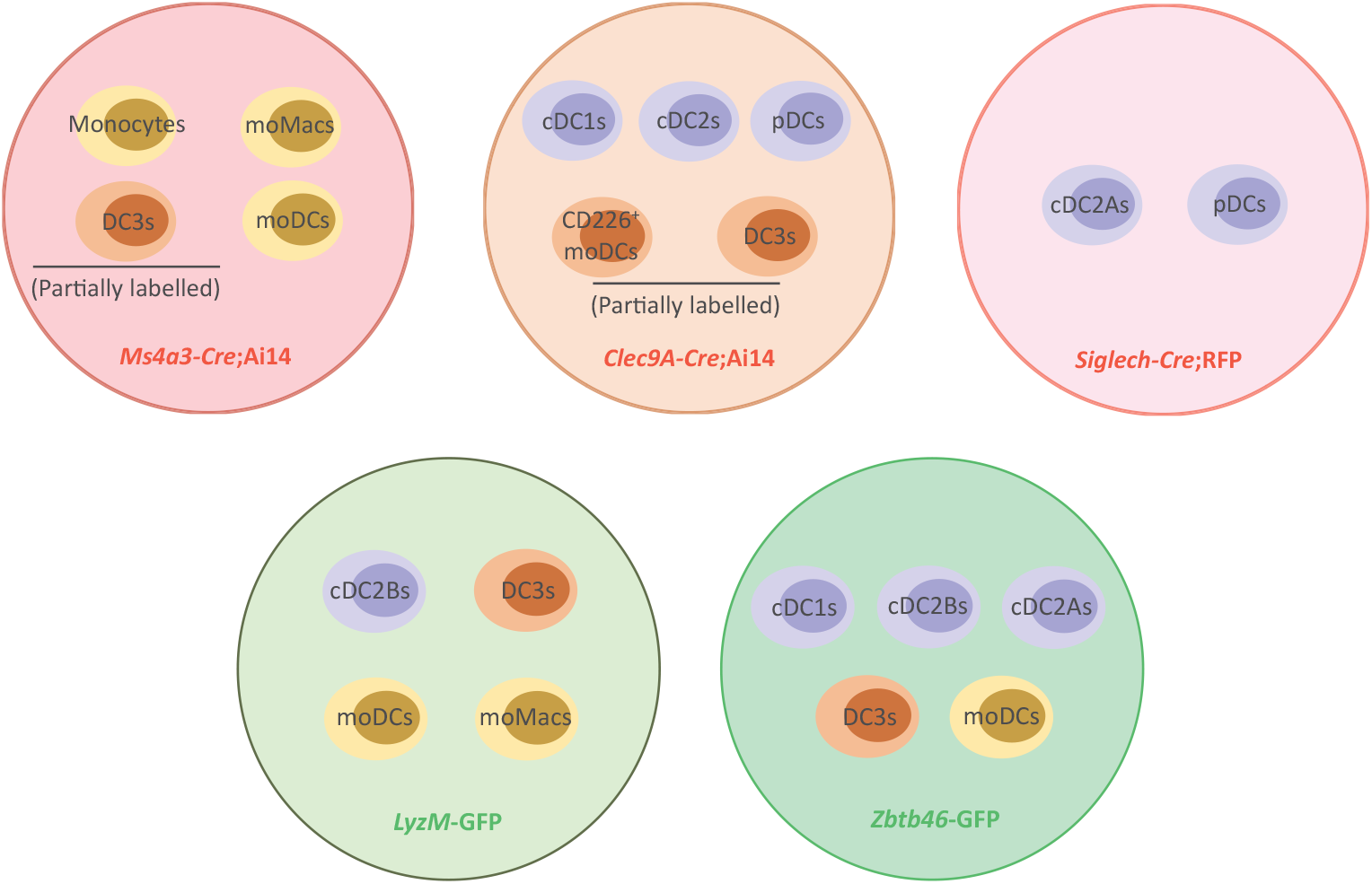
Graphical summary highlighting the specific cell populations labelled by each fate-mapping and constitutive reporter constructs used in the study.

**Supplementary Figure 2.**
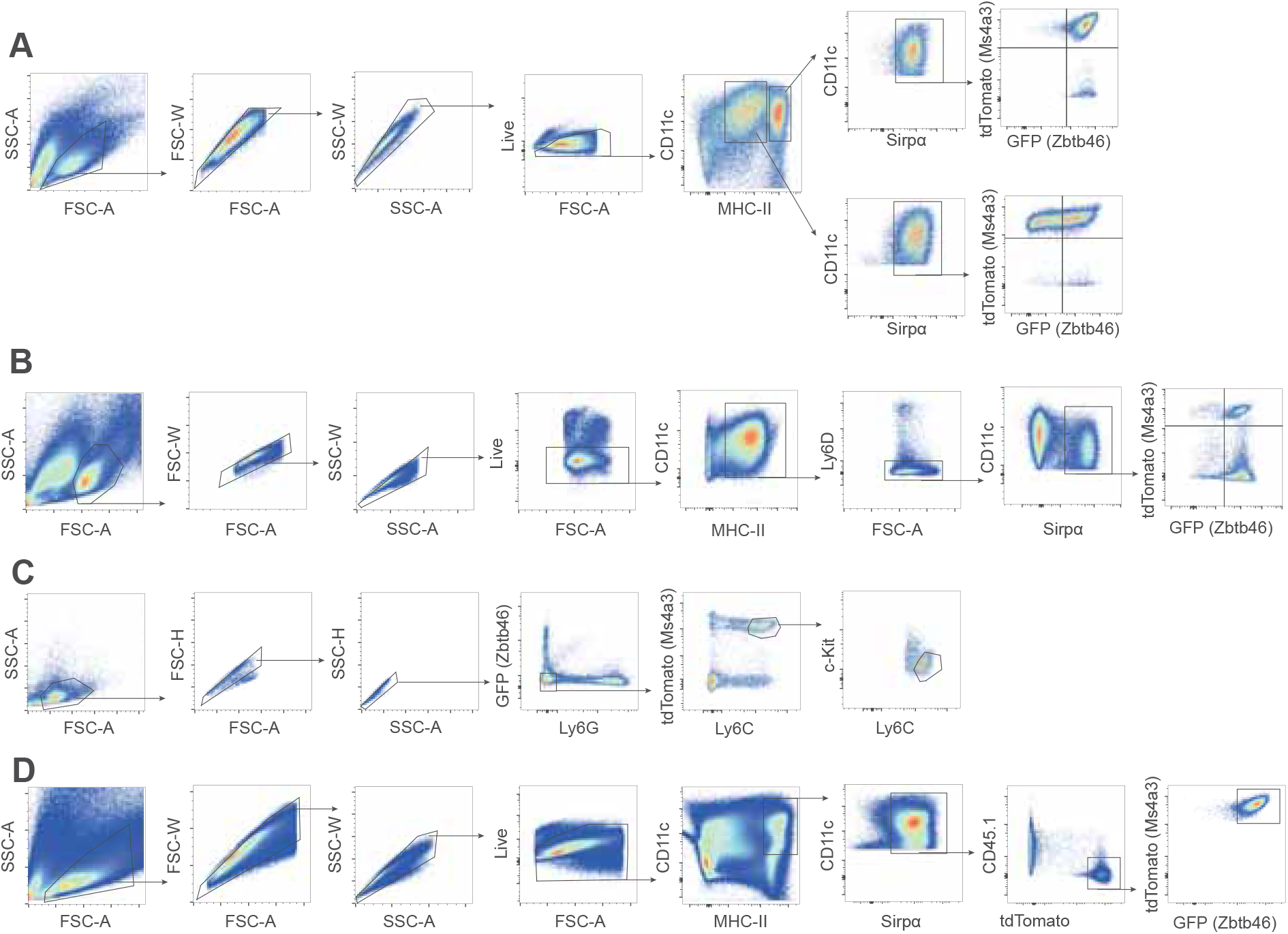
Main gating strategies employed to differentiate cDCs and moDCs in vitro. A) Gating strategy to identify moDCs in total bone marrow cultures supplemented with GM-CSF and IL-4 at day 8. B) Gating strategy to identify DC3s in total bone marrow cultures supplemented with FLT3-L supernatant at day 8. C) Gating strategy to FACS sort Ly6C^+^ Ai14^+^ classical bone marrow monocytes for moDC differentiation in vitro and adoptive transfer experiments. D) Gating strategy to identify moDCs derived from FACS sorted Ly6C^+^ Ai14^+^ classical bone marrow monocytes cultured in vitro in the presence of CD45.1 congenic bone marrow cells and GM-CSF and IL-4 for 4 days.

**Supplementary Figure 3.**
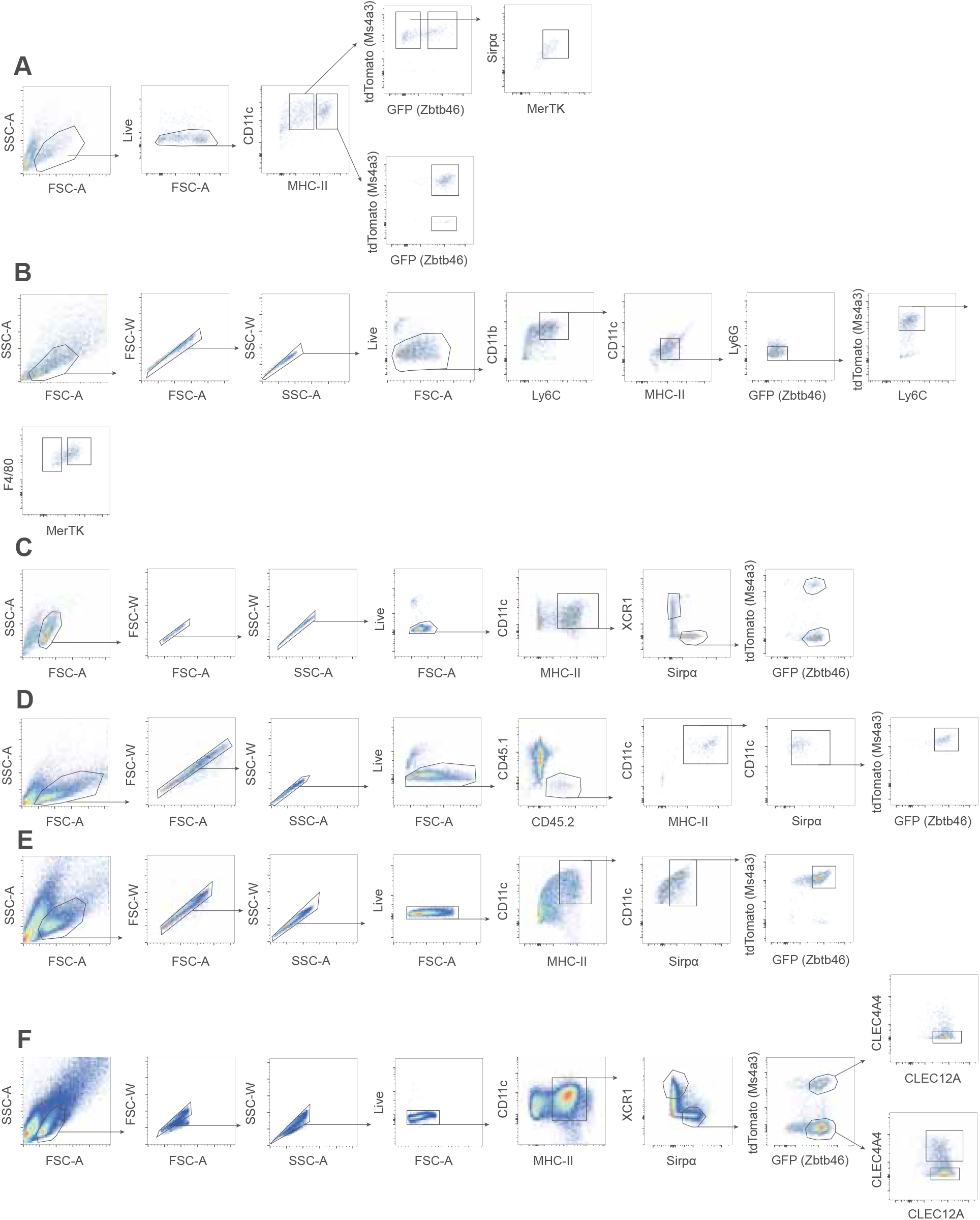
Gating strategies used to sort mononuclear phagocytes included in the various mini-bulk experiments. A) Gating strategy to sort moDCs and macrophages from total bone marrow cultures supplemented with GM-CSF and IL-4 at day 8. B) Gating strategy to sort macrophages from total bone marrow cultures supplemented with M-CSF supernatant at day 6. C) Gating strategy to sort cDC2s and DC3s from total bone marrow cultures supplemented with FLT3-L supernatant at day 8. D) Gating strategy to sort moDCs derived from FACS sorted Ly6C^+^ Ai14^+^ classical bone marrow monocytes cultured in vitro in the presence of CD45.1 congenic bone marrow cells and GM-CSF and IL-4 for 4 days. E) Gating strategy to sort moDCs from total bone marrow cultures supplemented with GM-CSF and IL-4 at day 8. F) Gating strategy to sort cDC2As, cDC2Bs and DC3s from total bone marrow cultures supplemented with FLT3-L supernatant at day 10 in the presence or absence of maturation cues.

**Supplementary Figure 4.**
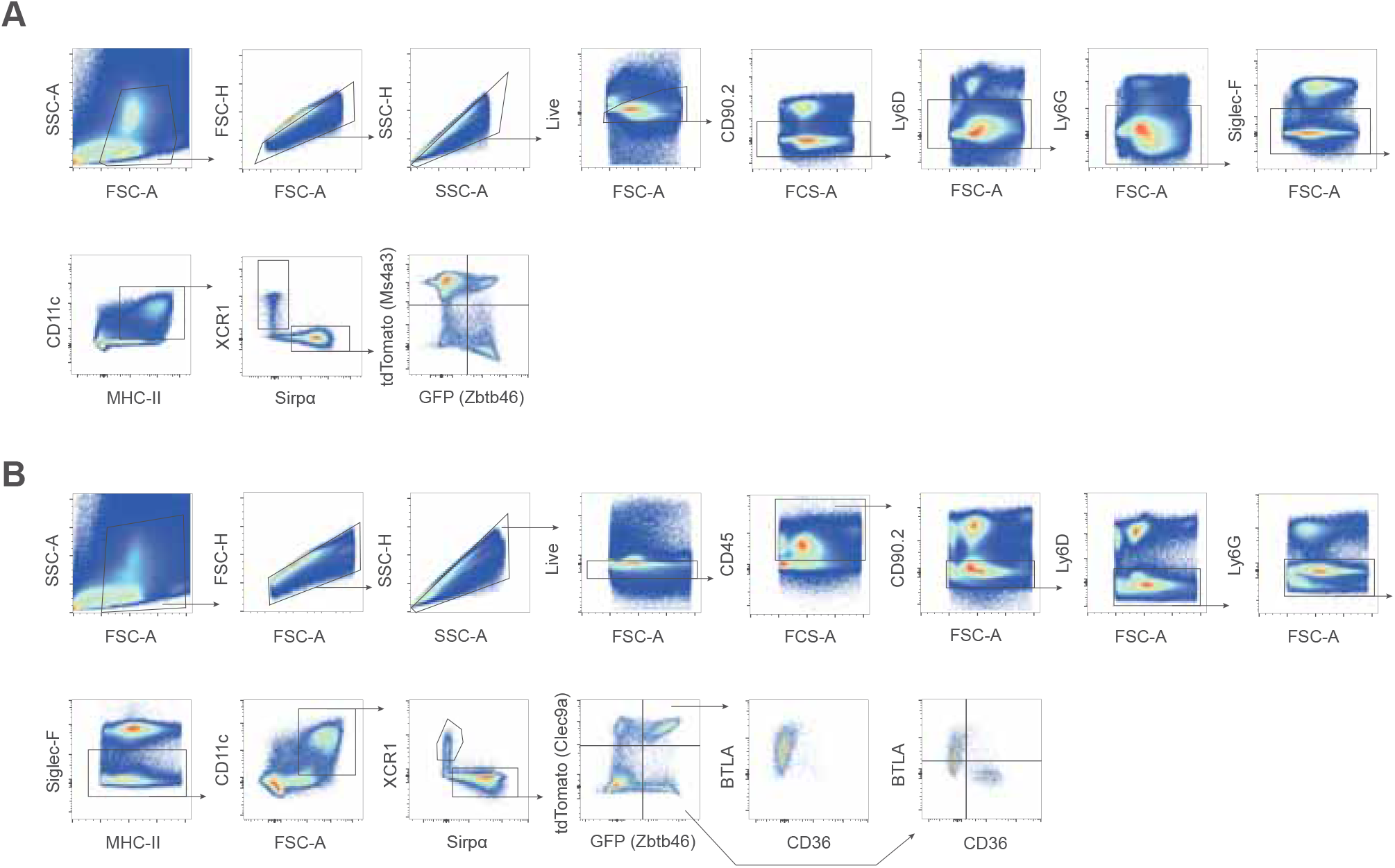
Gating strategies used to identify cellular fractions encompassing moDCs in *Ms4a3*-*Cre*;Ai14-*Zbtb46-*GFP and *Clec9a*-Cre;*Ai14*;*Zbtb46*-GFP mice treated with HDM. A) Gating strategy employed in *Ms4a3*-*Cre*;Ai14-*Zbtb46-*GFP mice. B) Gating strategy employed in *Clec9a*-Cre;*Ai14*;*Zbtb46*-GFP mice.

**Supplementary Figure 5.**
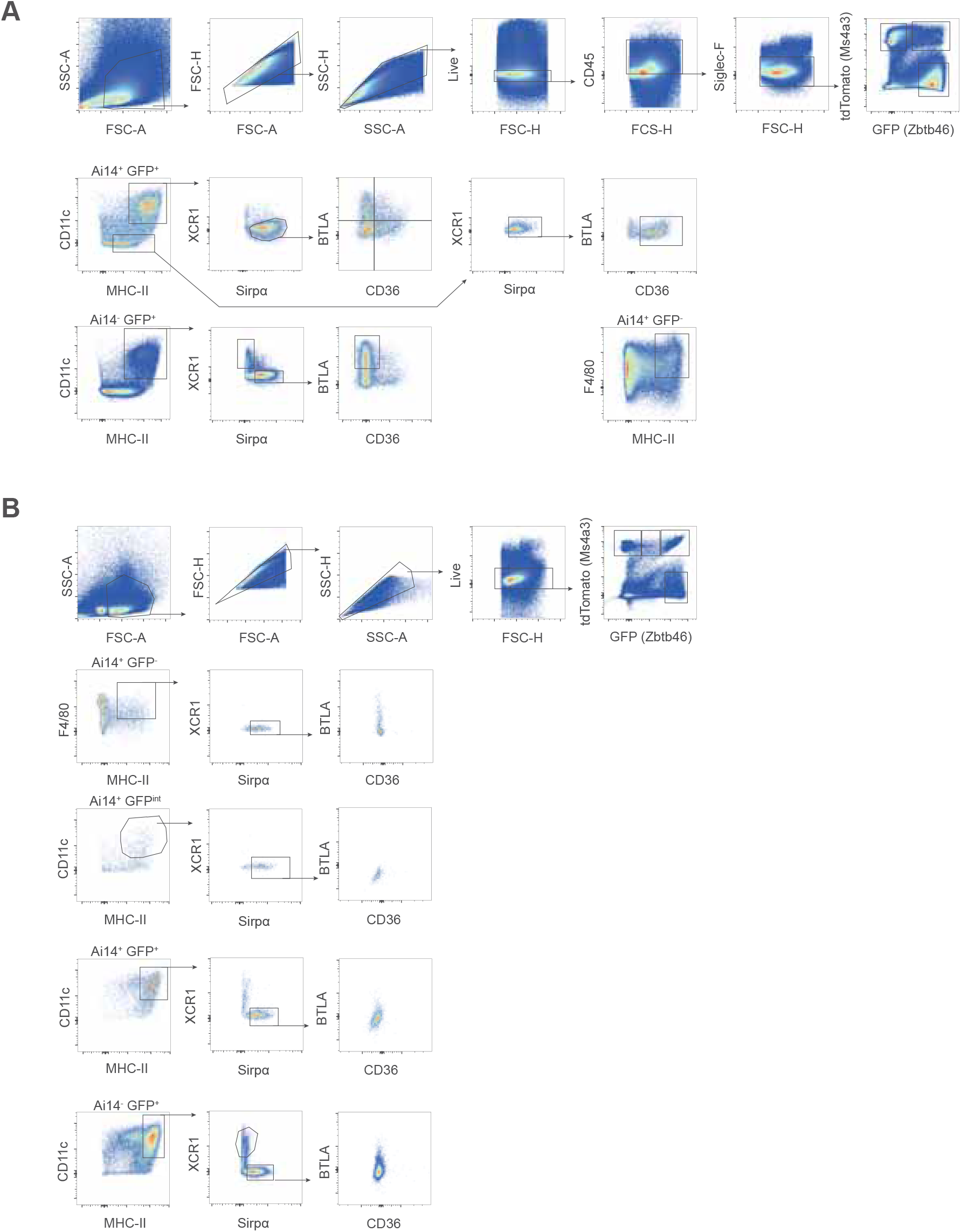
Identification of putative moDCs using BTLA versus CD36 expression in *Ms4a3*-*Cre*;Ai14-*Zbtb46-*GFP treated with HDM. A) Gating strategy used to identify various DC and macrophage populations in the lungs. B) Gating strategy employed various DC and macrophage populations in the lungs in the MedLNs.

**Supplementary Figure 6.**
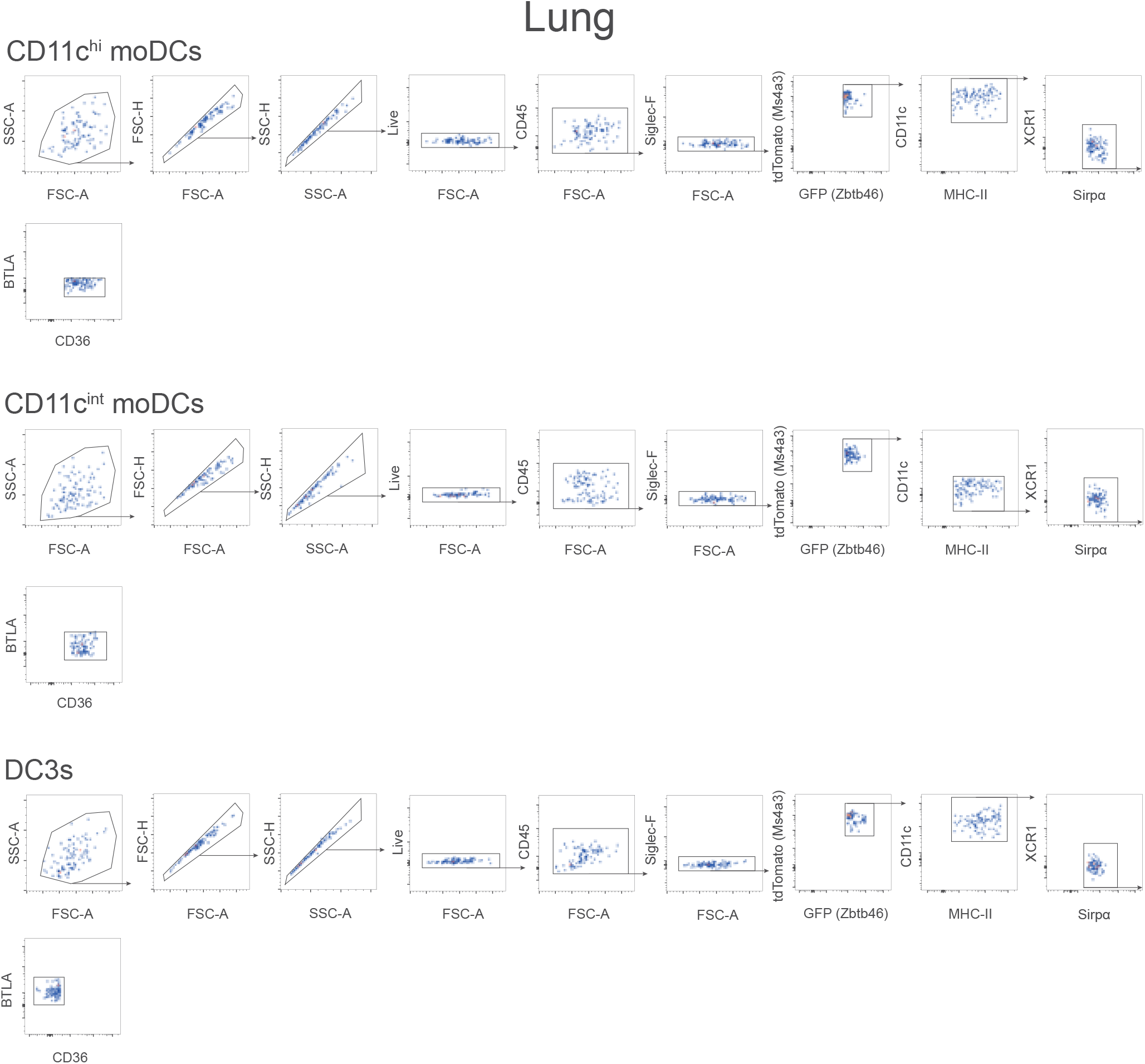
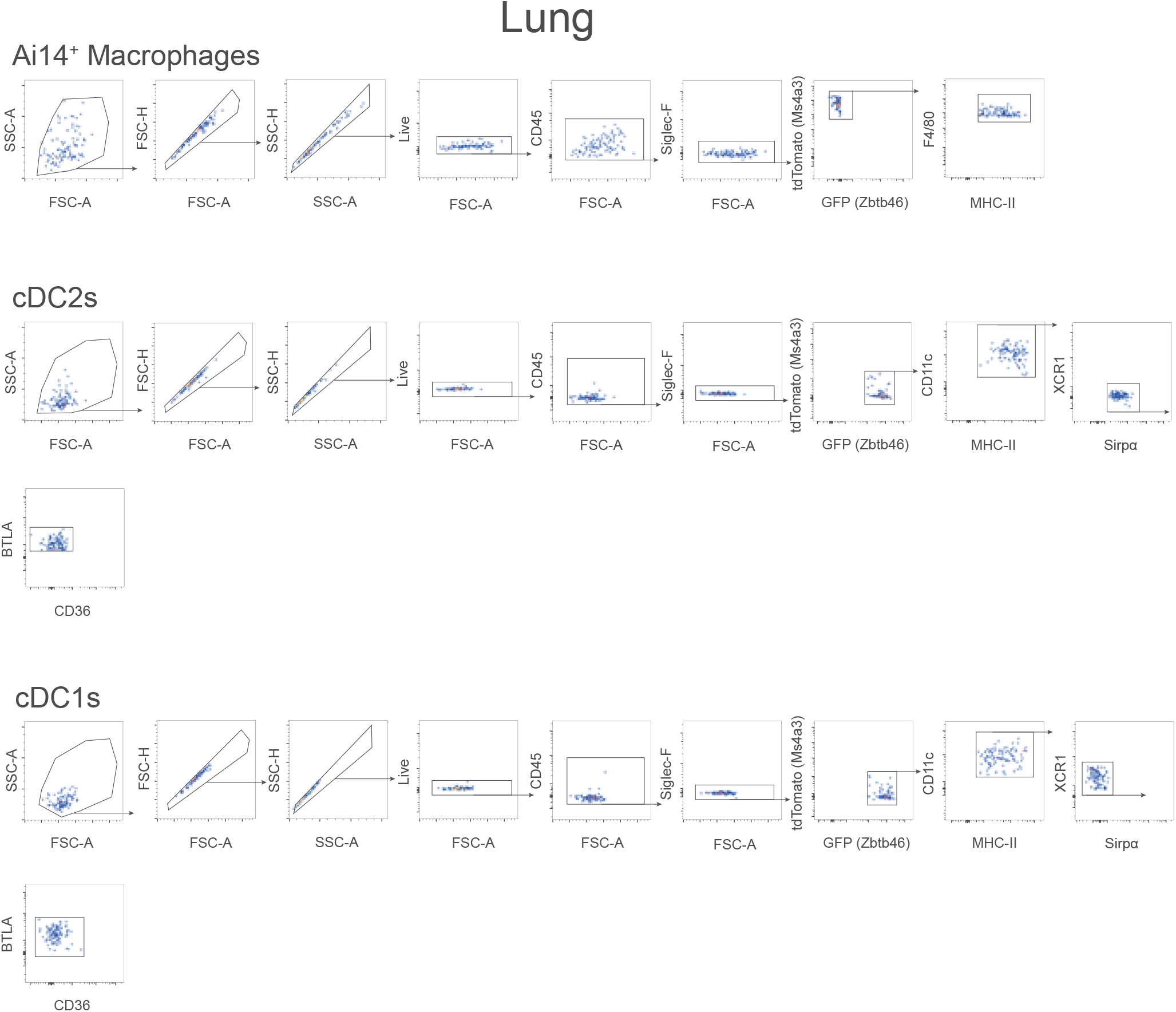
Index sorting flow cytometry data of various mononuclear phagocyte populations isolated from the lungs of *Ms4a3*-*Cre*;Ai14-*Zbtb46-*GFP treated with HDM for single cell RNAseq.

**Supplementary Figure 7.**
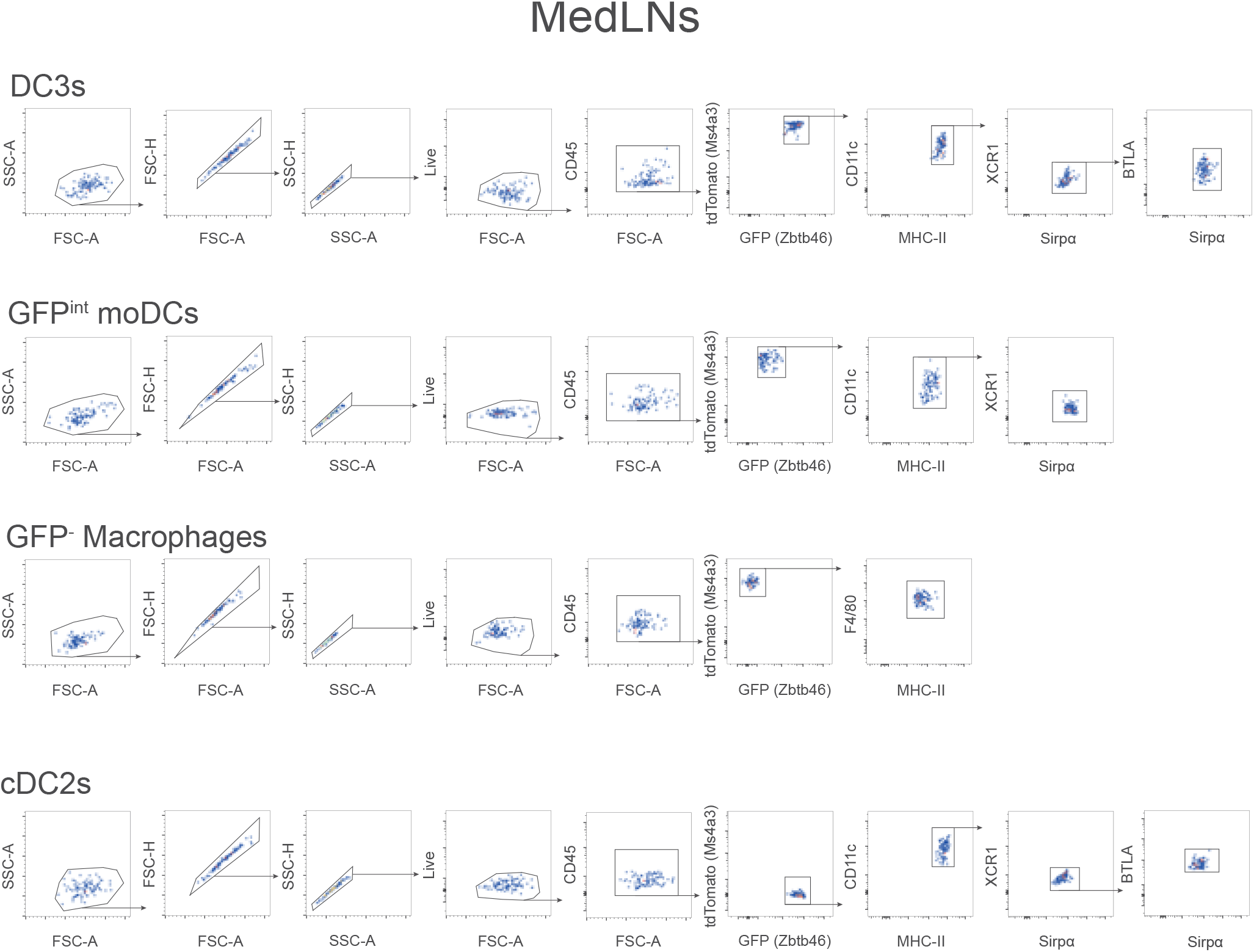
Index sorting flow cytometry data of various mononuclear phagocyte populations isolated from the MedLNs of in *Ms4a3*-*Cre*;Ai14-*Zbtb46-*GFP treated with HDM for single cell RNAseq.

**Supplementary Figure 8.**
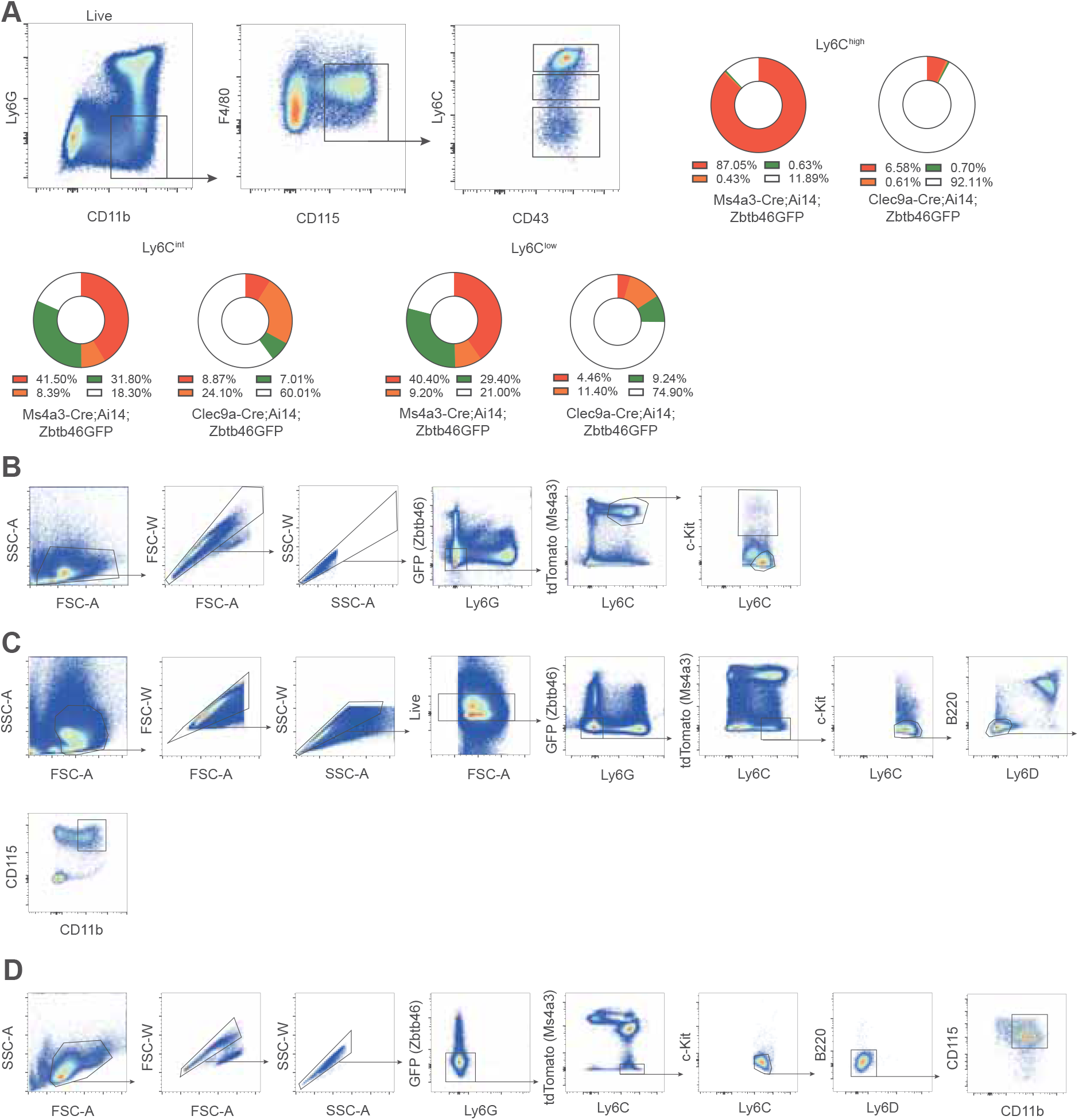
Ly6C expressing cells in the bone marrow are heterogeneous. A) Assessment of fate-mapped cells and cells expressing Zbtb46 using the gating strategy employed in Yanez et al., 2017 in bone marrow cells coming from *Ms4a3*-*Cre*;Ai14-*Zbtb46-*GFP and *Clec9a*-Cre;*Ai14*;*Zbtb46*-GFP mice. B) Ai14^+^ Ly6C^+^ monocytes in the bone marrow encompass a fraction of c-Kit expressing progenitors. C) Gating strategy to identify bone marrow Ai14^-^ Ly6C^+^ monocytes in the absence of purification kit. D) Gating strategy used to FACS sort, for adoptive transfer, Ai14^-^ Ly6C^+^ monocytes from bone marrow cell preparations coming from *Ms4a3*-*Cre*;Ai14-*Zbtb46-*GFP mice and enriched for the monocyte fraction using the EasySep mouse monocyte isolation kit from Stemcell Technologies.

**Supplementary Figure 9.**
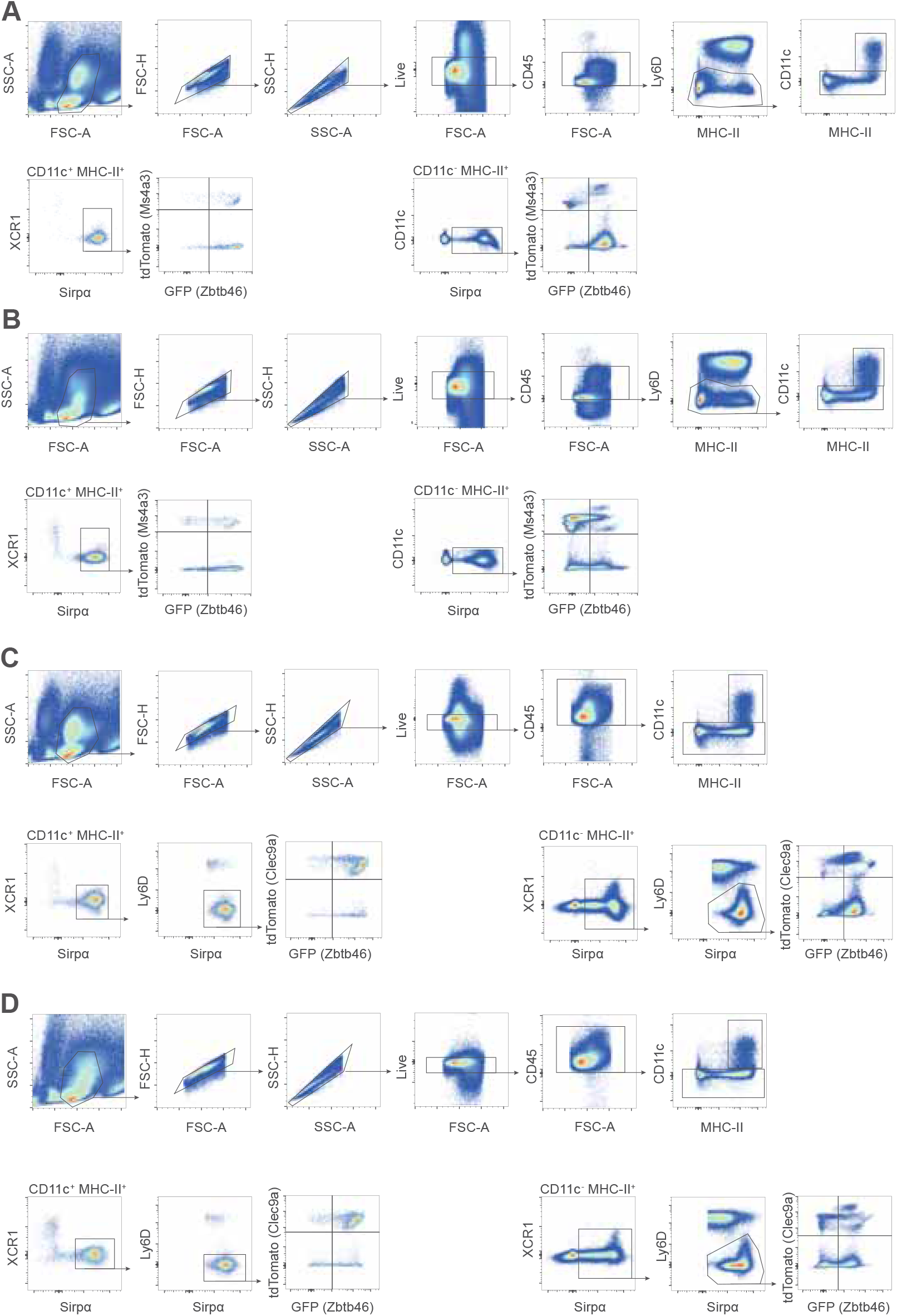
Assessment of fate-mapped cells and cells expressing Zbtb46 comparing untreated to *Ms4a3*-*Cre*;Ai14-*Zbtb46-*GFP and *Clec9a*-Cre;*Ai14*;*Zbtb46*-GFP mice challenged with thioglycolate. A) Peritoneal immune cell fractions from non-treated *Ms4a3*-*Cre*;Ai14-*Zbtb46-*GFP mice. B) Peritoneal immune cell fractions from *Ms4a3*-*Cre*;Ai14-*Zbtb46-*GFP mice treated with thioglycolate. C) Peritoneal immune cell fractions from non-treated *Clec9a*-Cre;*Ai14*;*Zbtb46*-GFP mice. D) Peritoneal immune cell fractions from *Clec9a*-Cre;*Ai14*;*Zbtb46*-GFP mice treated with thioglycolate.

**Supplementary Figure 10.**
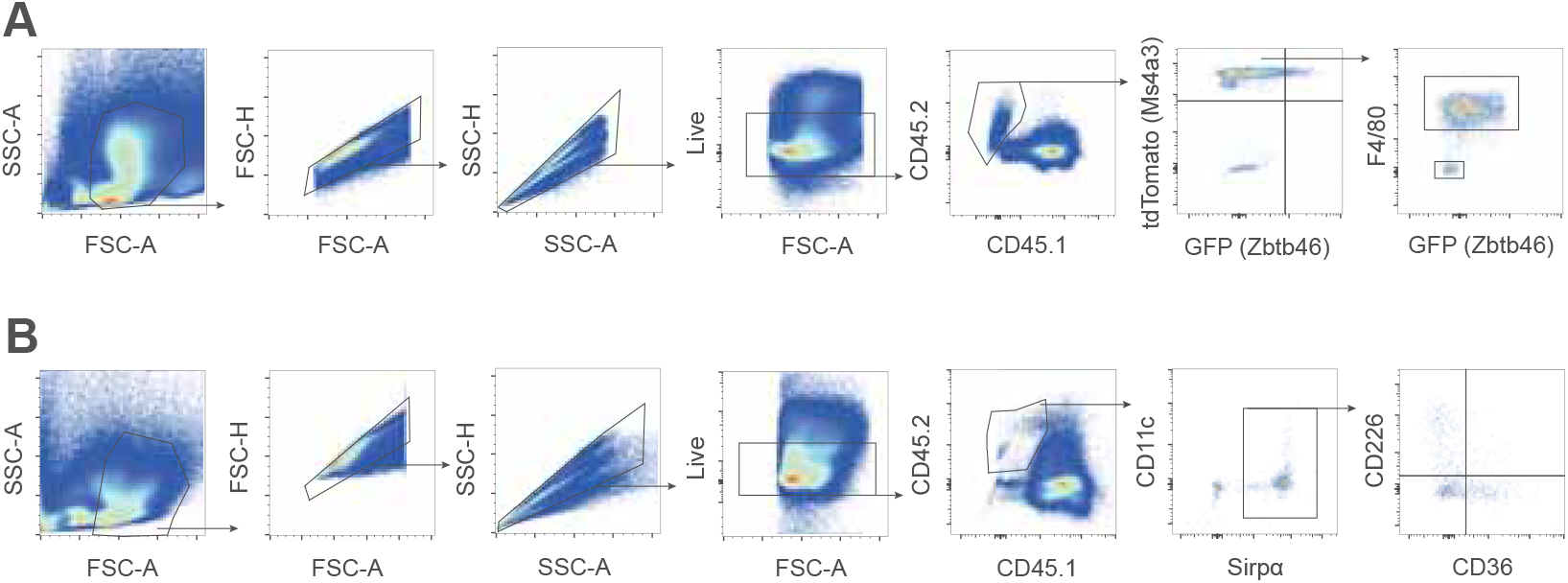
Adoptively transferred Ai14^+^ Ly6C^+^ or Ai14^-^ Ly6C^+^ monocytes give rise to various populations in the inflamed peritoneum. A) Gating strategy employed to identify the progeny of adoptively transferred Ai14^+^ Ly6C+ monocytes to CD45.1 mice treated with thioglycolate. B) Gating strategy employed to identify the progeny of adoptively transferred Ai14^-^ Ly6C^+^ monocytes to CD45.1 mice treated with thioglycolate.

**Supplementary Figure 11.**
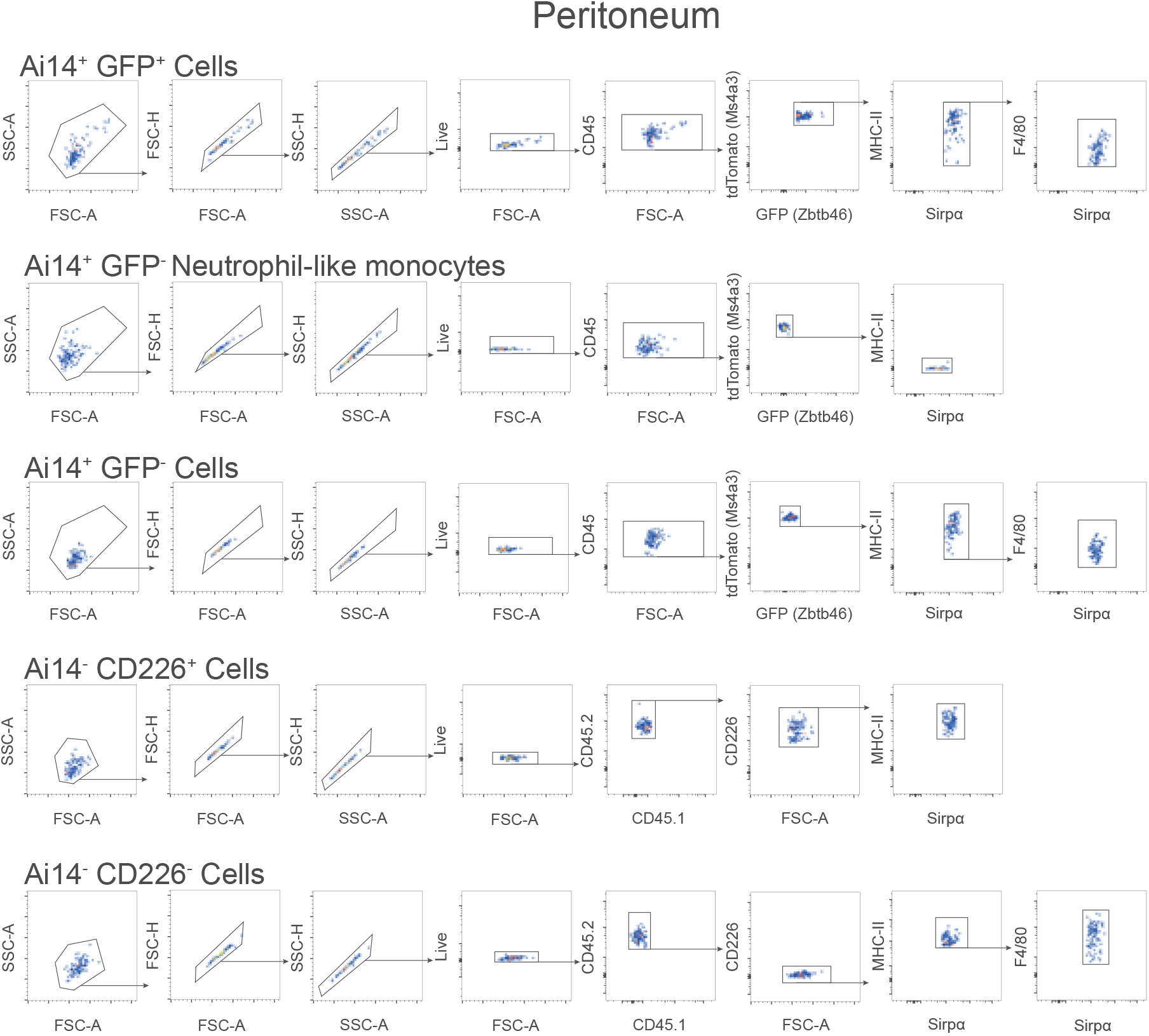
Index sorting flow cytometry data of various mononuclear phagocyte populations isolated from the peritoneum of thioglycolate treated mice following the adoptive transfer of Ms4a3;Ai14^+^ Ly6C^+^ or Ms4a3;Ai14^-^ Ly6C^+^ monocytes to C57BL6/J mice and to CD45.1 congenic mice respectively.

**Supplementary Figure 12.**
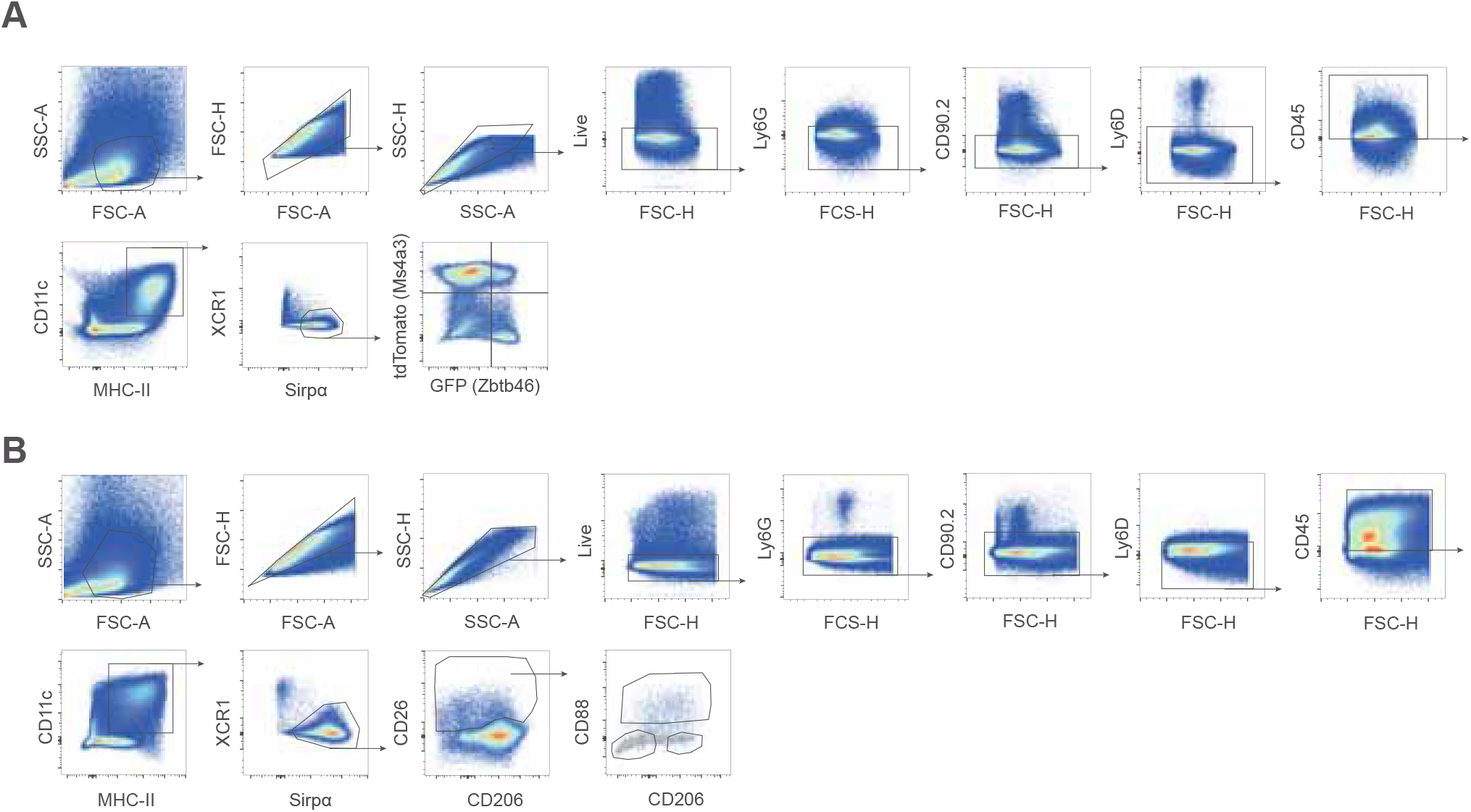
CD88 CD26 CD206 expression in moDCs in *Ms4a3*-*Cre*;Ai14-*Zbtb46-*GFP and C57BL/6 mice. A) Gating strategy used to test CD88, CD26 and CD206 in the lungs of *Ms4a3*-*Cre*;Ai14-*Zbtb46-*GFP treated with HDM. B) Validation of the gating strategy to identify moDCs in the lungs of C57BL/6 mice treated with HDM using the combination of CD88, CD26 and CD206.

## Notes

### Competing Interest Statement

The authors have declared no competing interest.

